# Mathematical Modelling of the Mitochondrial Dicarboxylate Carrier (SLC25A10)

**DOI:** 10.1101/2025.10.15.680887

**Authors:** Ramin Nashebi, Yingying Lyu, Elías Vera-Sigüenza, Daniel A. Tennant, Fabian Spill

## Abstract

The mitochondrial dicarboxylate carrier SLC25A10 mediates reversible exchange among succinate, malate, and phosphate, contributing to mitochondrial metabolic regulation. Structural studies establish a ping–pong mechanism, but most mathematical models still assume sequential binding, lacking mechanistic justification and overlooking the alternation of a single binding site. Here, we present the first mechanistically derived and thermodynamically consistent model of SLC25A10 based on a ping–pong framework. The model incorporates competitive binding of succinate, malate, and phosphate, heteroexchange, reversibility, and electroneutrality, and is calibrated using experimental datasets from intact mitochondria and reconstituted proteoliposomes. To estimate kinetic parameters and quantify their uncertainty, we employed Bayesian inference, enabling statistically rigorous calibration to uptake and competition assays. The model introduces new terms that quantify which substrate and from which side of the membrane is most likely to start the transport cycle. Beyond reproducing experimentally observed exchange kinetics, the model resolves non-equilibrium transport dynamics that are difficult to access directly in classical uptake assays. In particular, the simulations reveal a two-phase response in which an initial phosphate-driven high-flux uptake regime for malate and succinate is followed by a slower redistribution phase in which the two dicarboxylates continue to readjust primarily against each other. The model also predicts that mitochondrial morphology modulates early transport behaviour, with matrix swelling increasing and matrix condensation decreasing the initial SLC25A10 flux magnitude. More broadly, the framework provides a quantitative basis for studying how substrate competition, thermodynamic driving forces, and compartment geometry shape SLC25A10-mediated exchange, and it offers a transferable modelling strategy for other carriers in the SLC25 family.

## 2 Introduction

The solute carrier family 25 member A10 (SLC25A10), also known as the dicarboxylate carrier, is a transporter located in the inner mitochondrial membrane (IMM) [1, 2, 3, 4, 5, 6]. As its name suggests, this carrier facilitates the exchange of substrates containing two carboxyl groups, such as succinate and malate, together with other anions including inorganic phosphate, sulfate and thiosulfate [2, 4, 7, 8]. This exchanger plays a crucial role in the replenishment of intermediates of the tricarboxylic acid (TCA) cycle, supports oxidative phosphorylation, and maintains redox balance by enabling bidirectional exchange of metabolites [5, 4].

Under conditions such as cancer or ischaemia, SLC25A10 can become increasingly important in regulating succinate and malate levels [5, 2, 4, 9, 10, 11]. Evidence points towards SLC25A10 being a key player in the regulation of carbon metabolism under these circumstances, where its activity may contribute to the induction of a pseudo-hypoxic cellular state that promotes tumourigenesis [12, 13, 14].

Mechanistically, SLC25A10 functions as a secondary active antiporter. It does not consume ATP directly, but instead operates in response to transmembrane substrate gradients, particularly those involving phosphate and dicarboxylates [15, 16, 17, 2]. Transport is reversible, meaning that substrates can move in either direction depending on relative gradients, and it supports both homoexchange, the exchange of identical metabolites (e.g., succinate-for-succinate), and heteroexchange, the exchange of different metabolites (e.g., phosphate-for-succinate or malate-for-succinate). Since similar-charged molecules are exchanged in a strict 1:1 stoichiometry, the process is electroneutral and therefore insensitive to mitochondrial membrane potential [15]. Importantly, competition among transported substrates is well established, with malate generally exhibiting higher binding affinity than succinate and phosphate under classical assay conditions [3, 15, 16, 17].

Recent structural studies by Ruprecht et al. [18, 4] and Mavridou et al. [19] have shown that mitochondrial carriers of the SLC25 family, including SLC25A10, function as monomers with a single substrate binding site located in a water-filled cavity. This site is accessible from one side of the inner mitochondrial membrane at a time, consistent with an alternating-access mechanism. Complementary kinetic analyses by Cimadamore-Werthein et al. [1] demonstrated that these carriers operate through a ping–pong exchange mechanism, in which a substrate is bound, translocated and released before the counter-substrate can bind. Together, these structural and kinetic results support a single-displacement alternating mechanism, while challenging earlier sequential models that proposed that phosphate and dicarboxylates could bind simultaneously at distinct sites and be co-transported through conformational changes [3, 20, 21, 22, 23, 24, 15, 16, 17]. This combined evidence provides a strong motivation for revisiting the mathematical modelling of SLC25A10 within a ping–pong kinetic framework.

An additional but underexplored layer of regulation is the effect of mitochondrial morphology on SLC25A10 transporter fluxes. Both experimental and computational studies have demonstrated that alterations in the relative volumes of the mitochondrial matrix (MM) and intermembrane space (IMS) strongly influence the diffusion of metabolites and the exchange rates mediated by SLC25 [25, 26, 27, 28, 29]. In particular, swelling of the matrix or IMS dilutes metabolite pools and reduces effective concentration gradients. Because SLC25A10 transport relies on substrate gradients rather than ATP hydrolysis for energy, the relative volumes of the matrix and IMS directly determine the magnitude of its fluxes. Thus, it is worth exploring how volume changes modulate SLC25A10 activity at a coarse-grained level.

Several mathematical models have been developed to explore the kinetics of SLC25A10 [15, 16, 17]. While these models incorporate important features such as electroneutral heteroexchange, competitive inhibition, and thermodynamic constraints, they are generally based on a sequential mechanism and rely on the King–Altman method [30]—a graphical approach that simplifies kinetics but lacks mechanistic depth. As a result, these models are considered conceptual or proxy representations rather than mechanistically derived systems.

In this study, we present the first mechanistically derived and thermodynamically validated kinetic model of the mitochondrial dicarboxylate carrier SLC25A10 based on the ping–pong mechanism. Our framework integrates competitive binding of succinate, malate, and phosphate, together with reversibility, heteroexchange, and electroneutrality, and is calibrated against experimental data using Bayesian inference. In contrast to earlier sequential formulations, our model introduces conformational bias weights that quantify how different substrates bias the initiation of the ping–pong cycle, providing a mechanistic way to capture directional preferences in transporter activity. Beyond reproducing classical uptake and competition assays, the model provides access to non-equilibrium exchange dynamics that are difficult to measure directly, including transient two-phase behaviour and the modulation of early transport fluxes by mitochondrial compartment geometry. In this way, the present framework offers a quantitative and transferable basis for studying SLC25A10 and related mitochondrial carriers within a structure-informed transport modelling setting.

## 3 Model and Methods

### 3.1 Model Construction

The ping-pong kinetic mechanism describes an alternating-access process in which the transporter cycles between two distinct conformational states, each exposing a single substrate-binding site to one side of the membrane. This mechanism prevents simultaneous binding of counter-substrates and ensures strictly alternating translocation.

The SLC25A10 transporter is located in the IMM and mediates exchange of dicarboxylates and inorganic anions (e.g., succinate, malate, and phosphate) between the MM and the extra-matrix space. In vivo, these substrates originate from the cytosol (C) and can access the IMS by passing through the outer mitochondrial membrane (OM), which is highly permeable to small metabolites due to the presence of the voltage-dependent anion channel (VDAC) (see Fig. 1). Accordingly, in this study we treat the external compartment as the combined IMS and cytosol space (IMS+C) and the internal compartment as the MM, with transport occurring across the IMM (Fig. 2). For notational simplicity, we refer to this external compartment as the external side (*c*) throughout the manuscript.

**Figure 1:**
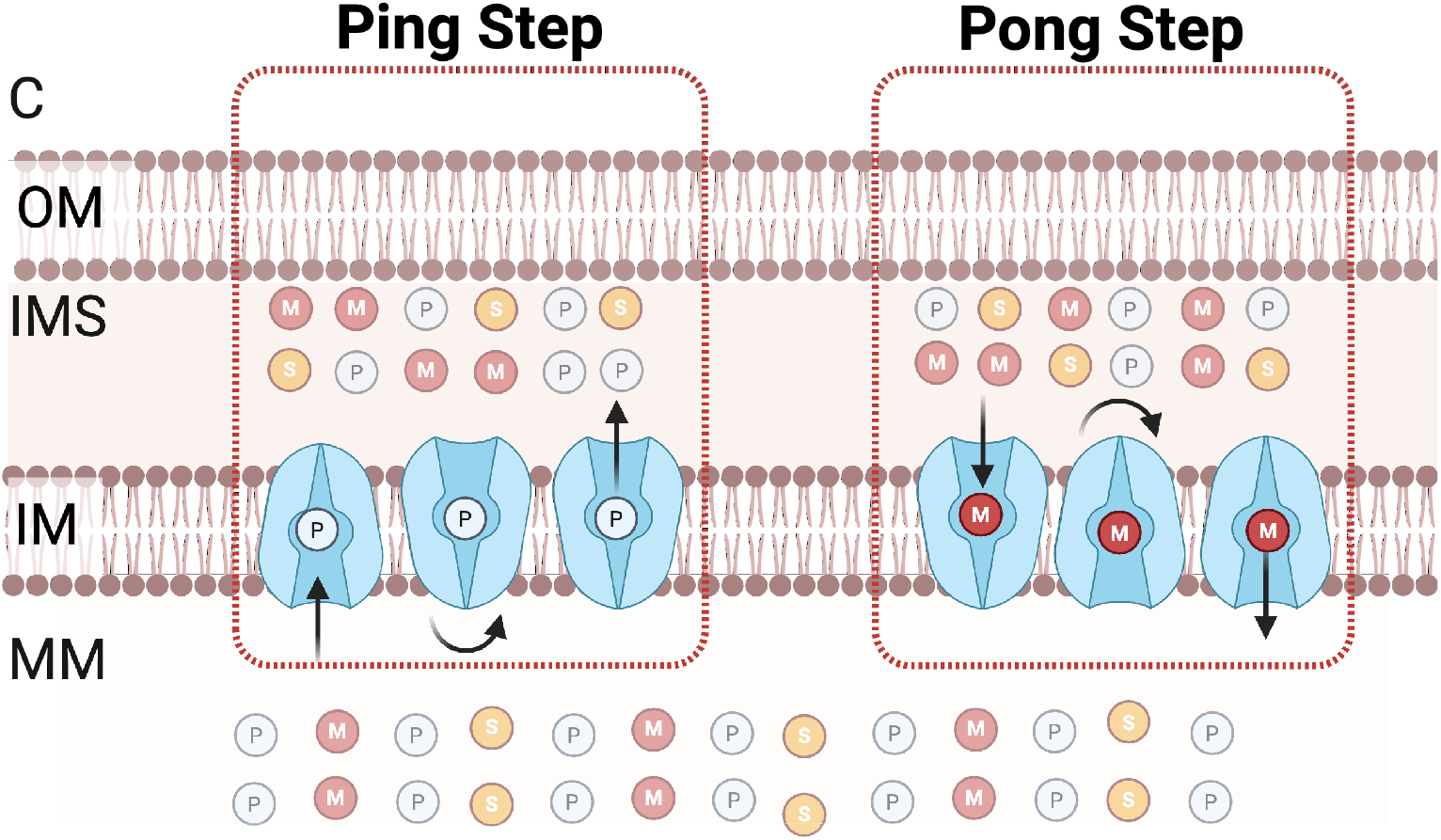
Schematic representation of the SLC25A10 transporter in the inner mitochondrial membrane (IMM), illustrating one representative phosphate-coupled malate exchange mode within the ping–pong mechanism. The dicarboxylates succinate (*S*, yellow) and malate (*M*, red) can, in general, be exchanged competitively with phosphate (*P*, white) across the IMM via SLC25A10. For clarity, this schematic shows only a malate/phosphate exchange example as an illustration of the single-binding-site alternating-access mechanism. In the *ping* step, phosphate is exported from the matrix to the external side; phosphate binding is followed by a conformational transition and substrate release. In the *pong* step, the vacant binding site is occupied by malate, which is then imported into the matrix; malate binding is followed by a conformational transition and release into the matrix. This schematic is not intended to imply that SLC25A10 operates intrinsically only in this mode; rather, the dominant exchange mode depends on substrate availability and transmembrane gradients. Created with BioRender.com.

**Figure 2:**
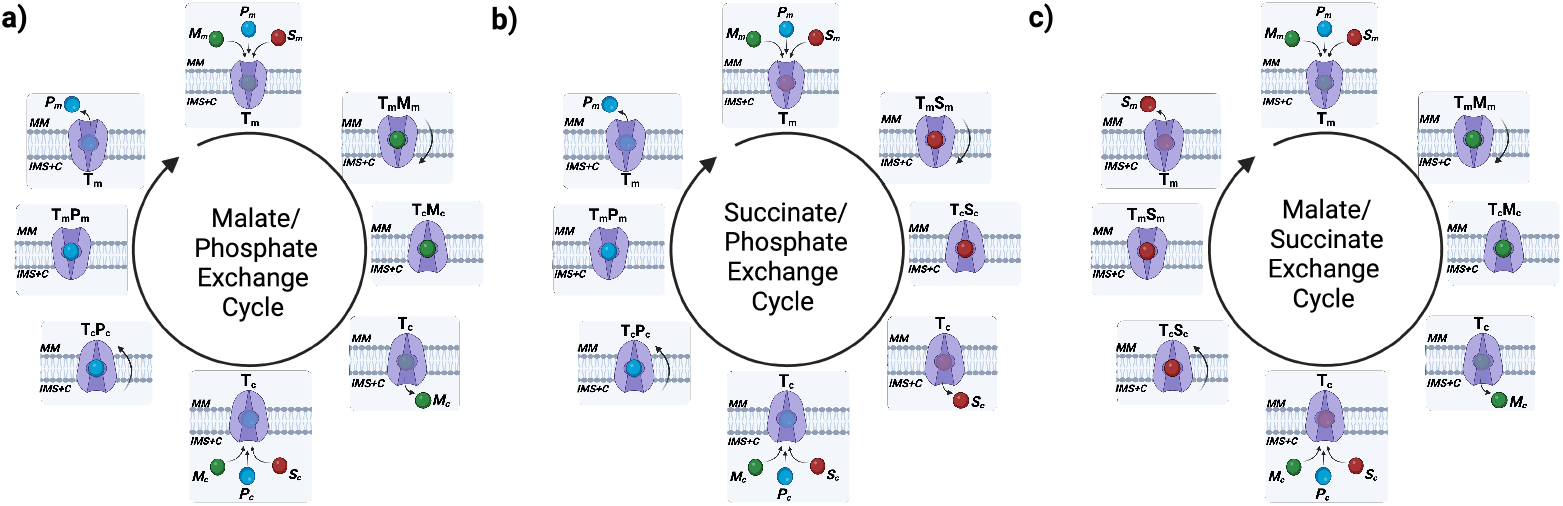
Ping–pong mechanism of the SLC25A10 transporter. The carrier alternates between a matrix-facing conformation (*T*_*m*_) and an externally facing conformation (*T*_*c*_), binding and translocating one substrate at a time through a single binding site. The external compartment (*c*) denotes the combined IMS+cytosol space (IMS+C). (a) Malate/Phosphate exchange cycle. (b) Succinate/Phosphate exchange cycle. (c) Malate/Succinate exchange cycle. In all cases, net directionality is not assumed a priori and is determined by the prevailing concentration gradients and reversible kinetics. Created with BioRender.com.

This transporter operates through a ping–pong mechanism. A central feature of this mechanism is that the net antiport direction is determined by the prevailing electrochemical driving forces (i.e., the relative substrate gradients and binding-site occupancy) [31, 32, 33]. In the ping–pong cycle, the carrier can bind a substrate on the side to which its binding site is exposed, undergo a conformational reorientation, and release that substrate on the opposite side, after which the counter-substrate binds and is translocated in the reverse direction (Fig. 2). Importantly, the cycle is fully reversible, so transport can proceed in either direction depending on the prevailing gradients, ensuring thermodynamic consistency and adaptability to changing metabolic demands [1]. Experimentally, the dicarboxylate carrier can support both heteroexchange and homoexchange among available substrates; in this work, we focus on the heteroexchange mode relevant to the calibration datasets.

In the mitochondrial matrix, we consider a system where mitochondrial succinate (*S*_*m*_), mitochondrial malate (*M*_*m*_), and mitochondrial phosphate (*P*_*m*_) compete for binding to the carrier. The transporter in its initial state, with the binding site facing the mitochondrial matrix, is denoted *T*_*m*_ (Fig. 2).

Binding of succinate, malate, or phosphate to *T*_*m*_ depends on their concentrations and binding affinities. When one substrate binds, a transient substrate–transporter complex (*T*_*m*_*S, T*_*m*_*M*, or *T*_*m*_*P* ) is formed [34, 1]. After binding, the carrier can undergo a conformational transition—driven by thermal fluctuations—that reorients the binding site toward the external side, yielding the state *T*_*c*_. In this configuration, the bound substrate can dissociate into the external compartment according to the modelled reversible binding/unbinding kinetics and be replaced by a counter-substrate for the return step [35].

Accordingly, three dominant heteroexchange modes can emerge under different gradient regimes: a malate/phosphate exchange cycle, in which malate and phosphate are the dominant transported substrates on the matrix-facing and external-facing sides, respectively (Fig. 2a); a succinate/phosphate exchange cycle, in which succinate and phosphate are dominant on the matrix-facing and external-facing sides, respectively (Fig. 2b); and a malate/succinate exchange cycle, in which malate and succinate are dominant on the matrix-facing and external-facing sides, respectively (Fig. 2c).

#### 3.1.1 Chemical Reaction Network of the Ping-Pong Mechanism

The reaction begins with the competitive binding of *S*_*m*_, *M*_*m*_, and *P*_*m*_ to the transporter in its matrix-facing conformation *T*_*m*_. Succinate binds to form the complex *T*_*m*_*S*, or malate binds to form *T*_*m*_*M*, while phosphate binds to form *T*_*m*_*P* . Thus, all three major substrates compete for a single binding site, and the dominant exchange behaviour emerges from the prevailing substrate gradients and kinetic parameters. These reversible reactions are characterised by forward 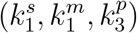 and backward 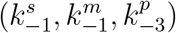 rate constants:

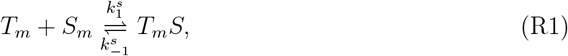

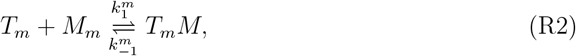

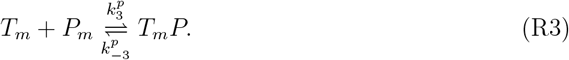

Once formed, the transporter–substrate complexes undergo conformational changes to expose the binding site to the external side. For succinate and malate, these transitions occur with rate constants 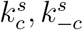 and 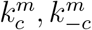, and for phosphate with rate constants 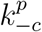 and 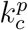, respectively:

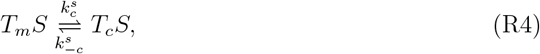

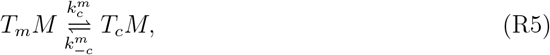

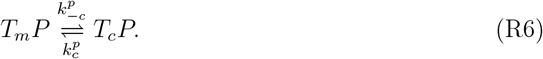

At the external side, the transporter releases the bound substrate, regenerating the free transporter in its external-facing state. These reactions leave the transporter in its modified state (*T*_*c*_), poised for subsequent interactions in the alternating-access cycle:

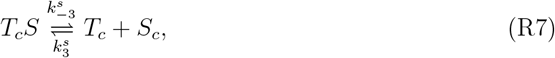

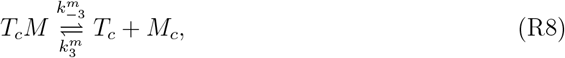

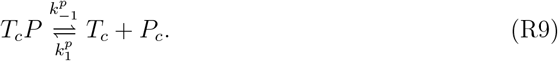

Consequently, the full cycle is reversible and can support both homoexchange and heteroexchange, depending on substrate availability and gradient direction.

### 3.2 System of Ordinary Differential Equations (ODEs)

The dynamics of the transporter system are described by a set of ordinary differential equations (ODEs) that capture the temporal evolution of transporter states, substrate complexes, and metabolite concentrations. To distinguish between MM and external side, we explicitly include their respective compartment volumes, *V*_*m*_ and *V*_*c*_, in the model. This ensures that fluxes are appropriately scaled by compartment size and that concentration changes are proportional to the fluxes across compartments. We further assume mass conservation and a spatially homogeneous system, since on the timescales considered, diffusion within each compartment is much faster than the transport processes being modelled. The complete ODE system is provided in Appendix A.

### 3.3 Model Assumptions and Derivation of Rate Equations for the SLC25A10 Transporter

We assume a symmetric kinetic parameterisation for SLC25A10 as a simplifying modelling assumption [31, 32, 33]. In this framework, the rate constants governing substrate binding, conformational transitions, and product release are taken to be equal in magnitude for transport in both directions. For example, in the case of succinate, the binding and release rate constants on the matrix side 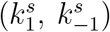 are assumed identical to the corresponding rate constants on the external side 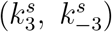. The same symmetry applies to malate and phosphate. This assumption reduces the number of independent kinetic parameters, thereby simplifying parameter estimation, while also ensuring that the transporter satisfies microscopic reversibility and detailed balance at equilibrium. Under this symmetry, the complete system of ODEs can be further reduced to a compact form, provided in Appendix C.

We further assume that substrate binding and dissociation for the transporter– substrate complexes (*T*_*m*_*S, T*_*m*_*M, T*_*m*_*P, T*_*c*_*S, T*_*c*_*M, T*_*c*_*P* ) occur on a faster timescale than the conformational transitions between matrix-facing and external-facing states [36]. This separation of timescales motivates treating binding and dissociation as quasi-equilibrated relative to the slower conformational changes, enabling a systematic reduction of the full carrier-state model by eliminating the fast binding/unbinding dynamics (see Appendix C).

Accordingly, we apply the rapid equilibrium assumption (REA) [37], which provides a principled way to reduce a mechanistic ping–pong transporter model to closed-form flux expressions while retaining reversibility, saturation, and substrate competition; the full mathematical derivation is provided in Appendix C.

Within this framework, we denote the net flux contributions of succinate, malate, and phosphate by *J*_suc_, *J*_mal_, and *J*_pho_, respectively, and adopt the sign convention that positive flux denotes net transport into the matrix (negative flux denotes net transport into the external compartment). Under this convention, *J*_suc_ is:

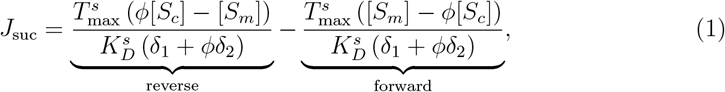

*J*_mal_ is:

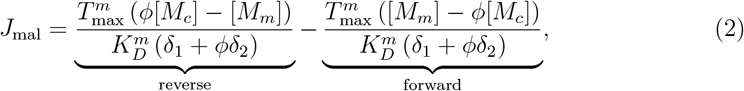

and *J*_pho_ is:

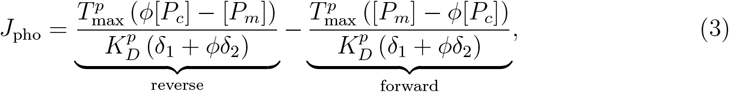

where

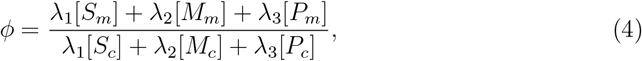

The dimensionless ratio *ϕ* compares the weighted presence of substrates on the two sides of the membrane, thus modulating the extent to which the matrix side pulls the transporter relative to the external side or vice versa. The parameters *λ*_1_, *λ*_2_, and *λ*_3_ are conformational bias weights assigned to succinate, malate, and phosphate, respectively. These quantify how strongly each bound substrate lowers the effective energy barrier for thermally driven conformational transitions and thereby biases the transport cycle. Specifically, *λ*_1_ corresponds to succinate, *λ*_2_ to malate, and *λ*_3_ to phosphate. Biologically, these parameters capture the competitive influence of each substrate at the single binding site of SLC25A10, where succinate, malate, and phosphate compete for access. The *λ*-weights thus scale the contribution of each substrate to the conformational bias term (*ϕ*).

It is important to clarify that, in the SLC25A10 ping–pong mechanism, the transport cycle is reversible. Depending on the prevailing concentration gradients, the transporter can initiate the cycle from either the matrix or the external side, and with either a dicarboxylate (succinate or malate) or phosphate as the starting substrate. For instance, if the cycle begins with phosphate, the counter-substrate may be phosphate, malate, or succinate; likewise, if it begins with a dicarboxylate, the counter-substrate may be phosphate, the other dicarboxylate, or the same dicarboxylate. The ratio *ϕ*, together with the weighting parameters *λ*_1_, *λ*_2_, *λ*_3_, governs this directional choice by determining which substrate and which side of the membrane most strongly bias the initiation of the transport cycle.

The terms *δ*_1_ and *δ*_2_ in Eqs. 1–3 are dimensionless partition functions that describe the availability of the transporter for binding on the matrix and external sides, respectively:

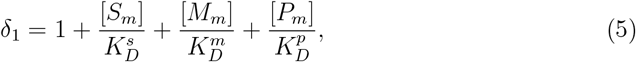

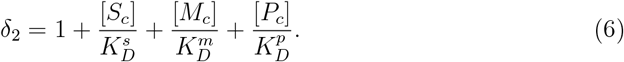

These include contributions from all three substrates, ensuring that succinate, malate, and phosphate compete for the same single binding site on whichever side of the membrane is exposed. The “1” term corresponds to the unbound transporter, while each substrate term (e.g., 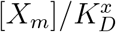 for the matrix or 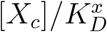 for the external side) represents the relative occupancy weight contributed by substrate *X*. Here, 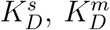, and 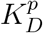 are dissociation constants representing the binding affinities of the respective substrates to the carrier.

Conceptually, *δ*_1_ and *δ*_2_ have the same form as classical binding polynomials used in enzyme kinetics [14]. Each *δ* represents the sum of all possible occupancy states on the matrix or external side. Dividing an individual term (e.g. 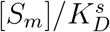) by *δ* gives the probability that the transporter is bound by that substrate. This highlights that our model treats transporter occupancy analogously to enzyme–ligand binding, but extended here to the ping–pong transport mechanism.

In Eqs. 1–3, 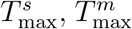, and 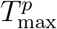 represent the apparent maximal transport capacities for succinate, malate, and phosphate, respectively. Each *T*_max_ reflects the product of the amount of active transporter (per gram of protein) and its turnover rate (per second) under saturating substrate conditions, thereby setting the upper bound of the corresponding flux (reported in mmol*/*s*/*g protein) in the absence of competitive inhibition or thermodynamic constraints. Differences among *T*_max_ values may arise from distinct translocation rates for each substrate or from asymmetries in binding and release kinetics across the two halves of the ping–pong cycle.

Since no leak pathways are considered and SLC25A10 is assumed to have a single binding site, each complete ping–pong cycle exchanges one bound substrate for one counter-substrate, with the identity of the exchanged pair determined by substrate availability and transmembrane gradients. At steady state, the flux through both half-cycles is therefore identical. Using the sign convention defined above, we can write:

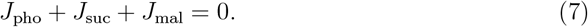

Finally, the overall transport magnitude of the dicarboxylate carrier is expressed as:

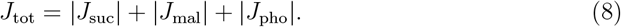

### 3.4 Reduced Model

Using the approximate flux expressions (*J*_mal_, *J*_suc_, and *J*_pho_), the full carrier-state ODE system can be reduced to a compartment-level mass-balance model for metabolite concentrations in the MM and the external compartment. Because the transporter fluxes reported in the experimental datasets used to calibrate the model are normalised per gram of protein, the concentration dynamics are written in terms of the compartment volumes per gram of protein, denoted 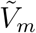 and 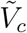. The reduced model is

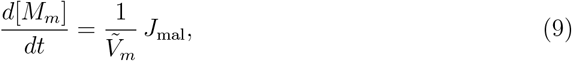

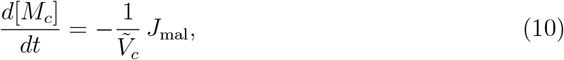

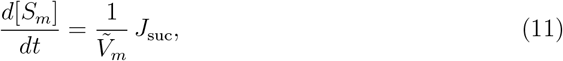

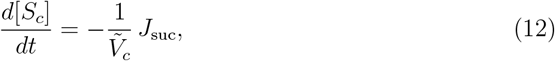

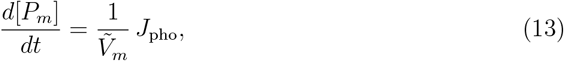

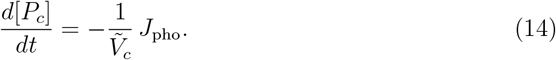

Here, positive fluxes are defined as transport into the matrix, such that the corresponding external compartment terms carry the opposite sign. This reduced model preserves mass balance across the membrane while capturing the essential dynamics of succinate, malate, and phosphate exchange through the SLC25A10 transporter.

### 3.5 Calibration Datasets and Experimental Conditions

In this study, we calibrated our mechanistic model using a combination of experimental datasets derived from both reconstituted proteoliposomes and intact rat liver mitochondria. Our study focuses on heteroexchange conditions in which a dicarboxylate (succinate or malate) is exchanged with inorganic phosphate. Consequently, we selected experimental datasets that specifically represent these heteroexchange conditions to constrain the model parameters.

For the core transport kinetics, we used data from reconstituted proteoliposomes reported by Indiveri et al. [23, 39], where the substrate composition on opposing sides of the membrane is strictly controlled. In particular, we selected datasets describing malate uptake against varying internal phosphate and phosphate uptake against varying internal malate (see Fig. 9).

To account for substrate competition and inhibition in a physiological setting, we incorporated data from intact rat liver mitochondria reported by Palmieri et al. [3]. Specifically, the datasets include succinate uptake rates measured over a range of external succinate concentrations, malate uptake rates measured over a range of external malate concentrations in the presence of external succinate, and phosphate uptake rates measured over a range of external phosphate concentrations (see Fig. 8). These measurements reveal competitive inhibition between malate, succinate, and phosphate for transport through the carrier and therefore provide key constraints on the competitive-binding component of the model.

Unlike the proteoliposome system, the internal substrate concentrations in intact mitochondria are not explicitly specified in these kinetic datasets. In our model, the external compartment concentrations of succinate, malate, and phosphate (*S*_*c*_, *M*_*c*_, *P*_*c*_) were set according to the experimental conditions in each dataset. The corresponding matrix concentrations (*S*_*m*_, *M*_*m*_, *P*_*m*_) were fixed based on the supporting literature values (see Table 1). Finally, the volumes of the matrix and external compartment (*V*_*m*_ and *V*_*c*_), together with the corresponding protein-normalised volumes 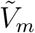 and 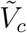, were fixed and calculated from the experimental sources used in this study (see Appendix F and Appendix G) and are summarised in Table 1.

**Table 1:** General Model Parameter Values.

| Parameter | Description | Value | Source |
| --- | --- | --- | --- |
| $\tilde{V}_m$ | Intraliposomal aqueous volume normalised per gram of protein (proteoliposomes) | $2.436 \text{ L g}^{-1}$ | Calculated; see Appendix F |
| $\tilde{V}_c$ | Extraliposomal assay medium volume normalised per gram of protein (proteoliposomes) | $59.4 \text{ L g}^{-1}$ | Calculated; see Appendix F |
| $V_m$ | Intraliposomal aqueous volume per assay (proteoliposomes) | $3.94 \mu\text{L}$ | Calculated; see Appendix F |
| $V_c$ | Extraliposomal assay medium volume (proteoliposomes) | $96.06 \mu\text{L}$ | Calculated; see Appendix F |
| $\tilde{V}_m$ | Sucrose-inaccessible aqueous matrix volume normalised per gram of protein (intact mitochondria) | $0.70 \mu\text{L mg}^{-1}$ | [38] |
| $\tilde{V}_c$ | External incubation-medium volume normalised per gram of protein (intact mitochondria) | $0.49 \text{ L g}^{-1}$ | Calculated; see Appendix G |
| $V_m$ | Sucrose-inaccessible aqueous matrix volume per assay (intact mitochondria) | $1.42 \mu\text{L}$ | Calculated; see Appendix G |
| $V_c$ | External incubation-medium volume per assay (intact mitochondria) | $1.0 \text{ mL}$ | Assumed |
| $[P_m]$ | Matrix phosphate concentration | $2 \text{ mM}$ | [14] |
| $[S_m]$ | Matrix succinate concentration | $0.2 \text{ mM}$ | [14] |
| $[M_m]$ | Matrix malate concentration | $0.2 \text{ mM}$ | Assumed |

### 3.6 Structural Parameter Identifiability Analysis and Reparameterisation

As a first step in model calibration [40], we analysed whether the model parameters can, in principle, be determined uniquely from perfect, continuous-time, noise-free measurements of the observables, based solely on the model structure and the available measurements. Here, the observables are the experimentally measurable model outputs, namely the net fluxes *J*_suc_, *J*_mal_, and *J*_pho_. To expose the structural symmetries of the reduced transport law, we first performed a local structural identifiability analysis of the reduced SLC25A10 ping–pong transport model written in terms of the nine quantities

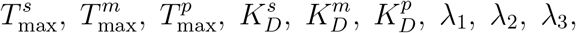

using the STRIKE-GOLDD method [41].

This analysis showed that the reduced model is not fully observable and that not all parameters are structurally identifiable under the current output configuration. Two main structural features underlie this loss of identifiability. First, in the reduced flux expressions, 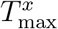 and 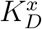 enter in coupled form for each substrate *x*, so that the observable transport law constrains their combined contribution to the fluxes rather than all underlying microscopic quantities independently. Second, in the *ϕ* expression, the parameters *λ*_1_, *λ*_2_, and *λ*_3_ enter only as proportional weights on succinate, malate, and phosphate concentrations. Consequently, scaling all three *λ*-parameters by a common constant leaves *ϕ* unchanged, so that only their relative combinations are structurally identifiable, not their absolute values.

To address this symmetry, we reparameterised the model by introducing the ratios

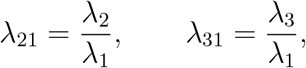

in place of the absolute *λ*-parameters in the *ϕ* term. Under the reduced flux formulation (see Appendix C), these ratios can be written as

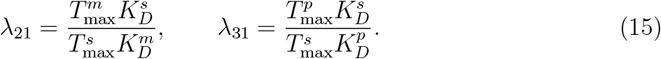

Thus, *λ*_21_ and *λ*_31_ are obtained as derived quantities from the transport-capacity and dissociation parameters, rather than being estimated independently. These ratios represent the relative weighting of malate and phosphate with respect to succinate in the conformational-bias term *ϕ*.

This identifiability analysis motivated the final inferential parameterisation used for Bayesian calibration. Rather than estimating the absolute *λ*_*i*_ values or a separate weighting-based parameter set, Bayesian inference was performed on the six core kinetic parameters

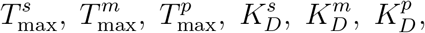

from which *λ*_21_ and *λ*_31_ were subsequently calculated as derived quantities. In this way, the structural-identifiability analysis was used to diagnose the symmetry of the original reduced formulation and to motivate the final six-parameter inferential model adopted throughout the remainder of the manuscript.

Although this reparameterisation resolves the symmetry in the *λ* terms, the practical identifiability of the remaining kinetic parameters still depends on the information content of the experimental data. Therefore, in settings where certain parameters are not sufficiently constrained by the available measurements, literature-based values or informative priors can be used to constrain the corresponding transport-capacity and dissociation parameters.

### 3.7 Local Sensitivity Analysis

To estimate the local sensitivity coefficients (LSCs), we compute the partial derivative of the flux with respect to a given parameter *θ*. This describes how small perturbations in that parameter change the flux. Each parameter *θ*_*i*_ in 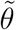 was perturbed individually by a factor *ε* = 0.1 (a 10% change), while keeping all other parameters fixed. The partial derivative was approximated using the central finite difference:

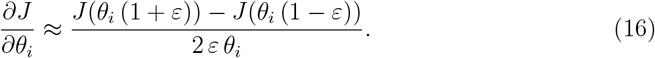

Since raw sensitivities depend on the units and magnitudes of both the flux and the parameter, they are not directly comparable across parameters. To address this, we use normalised LSCs. In this normalisation, each sensitivity is scaled relative to both the corresponding parameter value and the flux value. The normalised LSC is then defined as:

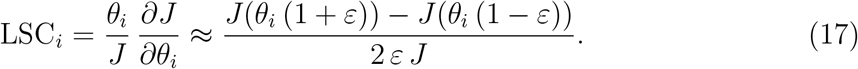

This renders the coefficients dimensionless, allowing the influence of parameters with different units (e.g. maximum transport rates versus dissociation constants) to be compared on the same scale. Intuitively, the normalised LSC represents the percentage change in a flux that results from a one percent change in a given parameter.

As a numerical accuracy check for the finite-difference sensitivities used throughout this work, we also compute the same local sensitivity coefficients using the complex-step method [42], which avoids subtractive cancellation and typically provides near machine-precision derivatives for analytic model evaluations.

### 3.8 Bayesian Framework for Parameter Estimation

To estimate the unknown parameters and quantify their uncertainties in the SLC25A10 transporter flux equations, we apply Bayesian inference using the Metropolis–Hastings Markov Chain Monte Carlo (MCMC) algorithm [43, 44]. The parameter vector 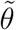 defined as

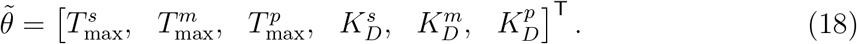

The weighting quantities are calculated as derived parameters from the inferred six-parameter set according to

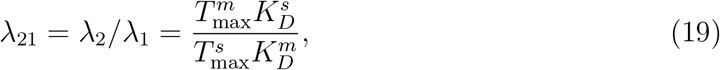

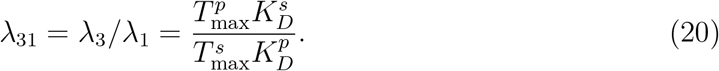

According to Bayes’ theorem, the conditional probability of the parameter vector 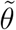 given the measured data *Y*, known as the posterior distribution, is expressed as:

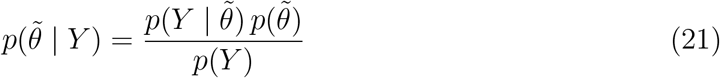

where 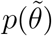 is the prior distribution of the unknown parameters, encoding available knowledge or plausible ranges before considering the experimental data. 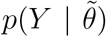 is the likelihood function, describing the probability of observing the experimental data *Y* given a specific parameter set 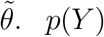 is the marginal likelihood (or evidence), acting as a normalising constant ensuring 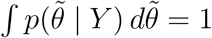. In practice, *p*(*Y* ) is difficult to compute directly, so inference is carried out up to a proportionality constant:

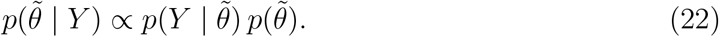

#### 3.8.1 Likelihood Function

The results of our SLC25A10 model are always subject to error because it is only an approximation of the underlying transport process. If we consider the experimental measurements, *Y*, as the observation of the transport fluxes, then the model predictions with parameters 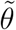, denoted 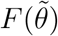, can be expressed as

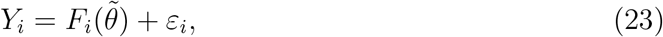

where *ε*_*i*_ represents the error in the model and *i* indexes the experimental data points. The error is assumed to follow an independent Gaussian distribution with zero mean and constant variance *σ*^2^. Under these assumptions, the likelihood function for the data *Y* given the parameters 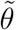 and variance *σ*^2^ can be written as

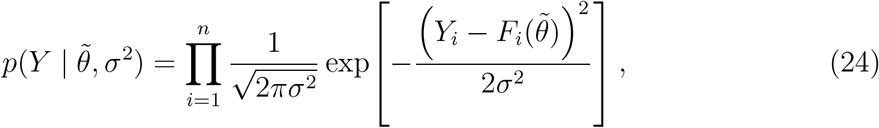

where *n* is the total number of flux measurements across all experiments.

Since the error variance *σ*^2^ is unknown, it is treated as an additional parameter to be inferred jointly with 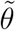 in the Bayesian framework. For a calibration with *d*_*θ*_ kinetic parameters, we introduce the overall parameter vector 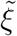, defined as:

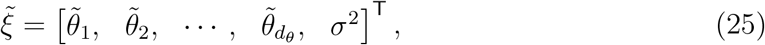

To simplify our computations, we take the natural logarithm of the likelihood function. The log-likelihood is then proportional to the sum of squared residuals between the measured and model-predicted fluxes:

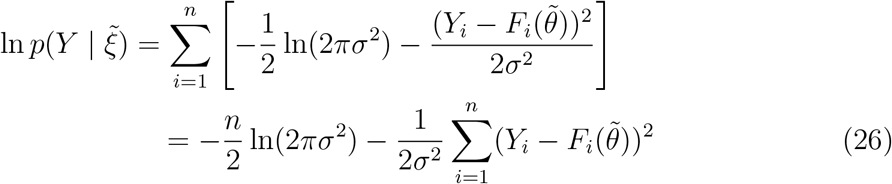

We emphasise that our assumption of independent, identically distributed Gaussian errors is not intended to fully capture the complexity of experimental uncertainty or model–data discrepancy. Instead, we regard this as a proof-of-concept choice that enables a tractable Bayesian calibration of a mechanistic model grounded in real physiological dynamics. The Gaussian assumption allows for closed-form likelihoods and efficient computation, which is essential at this stage.

#### 3.8.2 Prior

The prior distributions of the unknown parameters 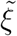 were assigned based on the results of the structural identifiability analysis and available experimental data.

For the intact-mitochondria transport assays, we assigned independent uniform priors to the six kinetic parameters over physiologically reasonable ranges, reflecting the absence of strong prior constraints:

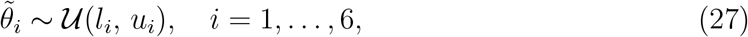

with probability density function

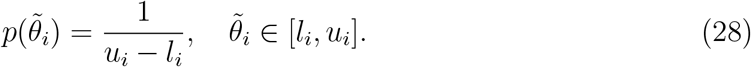

For the reconstituted proteoliposome transport assays, the experimental system probes only malate–phosphate exchange, with succinate absent. Accordingly, Bayesian inference in this setting was restricted to four kinetic parameters,

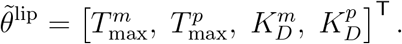

The observation-error variance *σ*^2^ was inferred separately in both experimental settings. For the four proteoliposome kinetic parameters, we assigned Gaussian priors using literature values where available, as summarised in Table 2:

**Table 2:** Literature values of kinetic parameters for reconstituted proteoliposome transport assays, with mean, standard deviation, 95% confidence intervals, and references.

| Parameter | Mean | Std | 95% CI (Low–High) | Reference |
| --- | --- | --- | --- | --- |
| $T_{\max}^m$ (mmol/min/g protein) | 6.00 | 2.40<sup>a</sup> | 1.30 – 10.70 | [23] |
| $T_{\max}^p$ (mmol/min/g protein) | 6.00 | 2.40<sup>a</sup> | 1.30 – 10.70 | [23] |
| $K_D^m$ (mM) | 0.49 | 0.05 | 0.392 – 0.588 | [23] |
| $K_D^p$ (mM) | 1.41 | 0.35 | 0.724 – 2.096 | [23] |

<sup>a</sup> Standard deviations for $T_{\max}^m$ and $T_{\max}^p$ were assumed for the Gaussian priors.

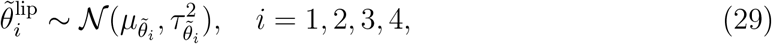

with probability density function:

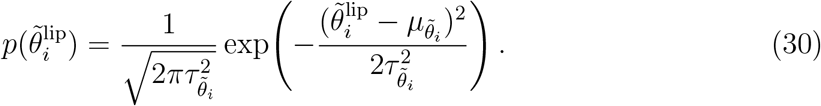

In this proteoliposome setting, the weighting parameters *λ*_21_ and *λ*_31_ are not estimated separately. Instead, their identifiable ratio is computed deterministically from each posterior sample as

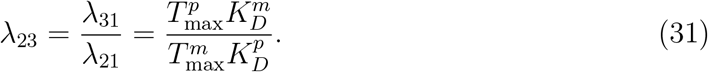

For both experimental settings, the observation-error variance *σ*^2^ was assigned an inverse-gamma prior with shape parameter *α* and scale parameter *β*:

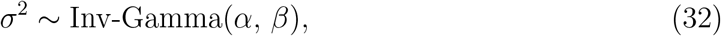

with probability density function:

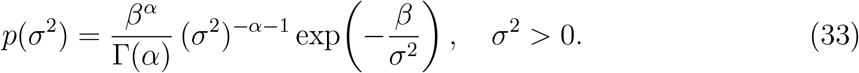

Assuming a priori independence, the joint prior distribution factorizes as:

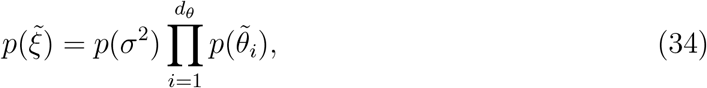

where *d*_*θ*_ = 6 for the intact-mitochondria calibration and *d*_*θ*_ = 4 for the proteoliposome calibration, with each term defined according to the distributional choices described above. To simplify computations, we used the logarithm of the prior distribution:

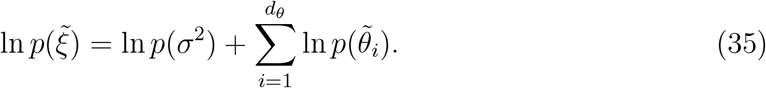

#### 3.8.3 Posterior Distribution

Given the likelihood function 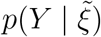 in Eq. (24) and the independent priors 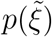 defined in Eq. (34), the posterior distribution of the parameter vector is obtained using Bayes’ theorem (Eq. (22)).

For computational convenience and to ensure positivity, we work with the logarithm of the posterior distribution. Combining Eq. (26) and Eq. (35), the log-posterior can be written as:

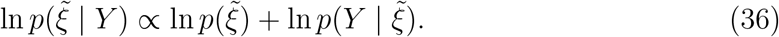

#### 3.8.4 Markov Chain Monte Carlo (MCMC) Method

Direct analytical evaluation of the posterior distribution 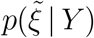 is generally intractable due to the complexity of the nonlinear transporter model. Therefore, we employed a Markov Chain Monte Carlo (MCMC) approach to generate samples from the posterior distribution and estimate the kinetic parameters [43].

We used the Metropolis–Hastings algorithm to construct the Markov chain 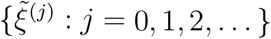, where *j* denotes the sampling iteration. At each iteration *j*, a new candidate 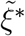 was generated using a log-normal random walk proposal:

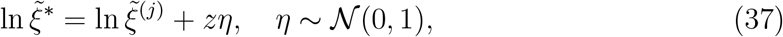

which ensures positivity of all parameters. Here, *z* is the step size, set to *z* = 0.1 in our calculations.

The acceptance probability is given by:

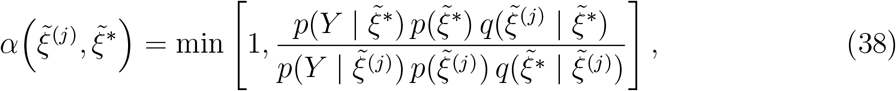

where *q*(·|·) denotes the log-normal proposal density. Since the log-normal proposal distribution is asymmetric, the Hastings correction (the ratio of proposal densities) is included to guarantee detailed balance and ensure that the posterior distribution remains the stationary distribution of the chain [44].

If *α* ≥ *r*, where *r* ∼ U(0, 1), the candidate is accepted:

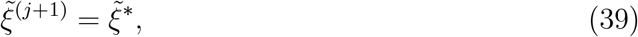

otherwise, the current state is retained:

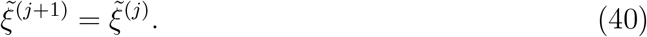

At each iteration, the fluxes *J*_suc_, *J*_mal_, and *J*_pho_ were computed using the transporter flux equations. We used all three datasets adopted from [3], which are described above. These datasets together comprise *n* = 18 data points and were used as the observation vector *Y* .

#### 3.8.5 Estimation of Monte Carlo Standard Error (MCSE)

After generating a sequence of samples from the posterior distribution using the Metropolis–Hastings MCMC algorithm, we used these samples to estimate posterior quantities such as the mean, median, and credible intervals. However, because the number of samples is finite and they are often correlated due to the Markov chain structure, each sample depends on the previous one [45, 46, 47, 48], these estimates have some variability. The MCSE quantifies the error in an MCMC estimate (e.g. the posterior mean) that arises from sampling variability.

To calculate the MCSE of a kinetic parameter of interest *θ*_*i*_ from a chain of *N* samples, we first compute the sample mean:

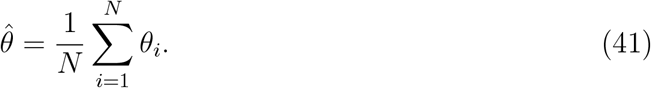

We then compute the sample variance of the MCMC samples:

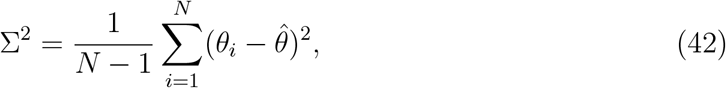

which represents the variability of the parameter values in the chain.

Since MCMC samples are correlated, the effective sample size (*N*_eff_) is typically smaller than *N* . To estimate *N*_eff_, we compute the autocorrelation at lag *k*, denoted *ρ*_*k*_, using the autocorrelation function (ACF) [45]:

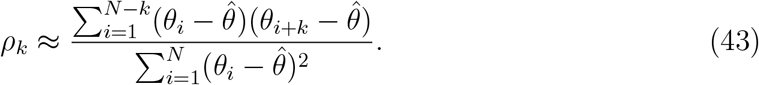

The effective sample size is then estimated as:

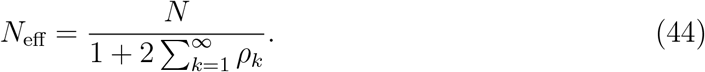

In practice, the sum is truncated at a lag where *ρ*_*k*_ becomes negligible [48].

Finally, the MCSE for the posterior mean is given by:

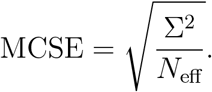

This formula accounts for correlation among the MCMC samples. A small MCSE indicates that the MCMC estimate is precise, meaning that the chain length is sufficient to approximate the true posterior statistic. In this study, we aimed for an MCSE smaller than 10% of the posterior standard deviation (Σ), which reflects the intrinsic uncertainty of the parameter under the posterior distribution. While we cannot guarantee the correctness of the finite set of *N* samples, by calculating the MCSE-to-SD ratio and targeting values below 0.10, we empirically assessed the convergence and reliability of the chains.

#### 3.8.6 Estimation of 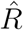 Statistic

To assess the convergence of our chains to the equilibrium or stationary distribution, we estimate the Gelman–Rubin statistic, 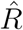 [49, 50]. This diagnostic evaluates whether independently initialised chains target the same stationary posterior. It does so by comparing between-chain to within-chain variability for each parameter.

Suppose there are Ω chains, each with *N* draws for a parameter *θ*. Let the draws from chain *ω* ∈ Ω be denoted *θ*_*ω,i*_, where *i* indexes the samples. The chain mean for chain *ω* is

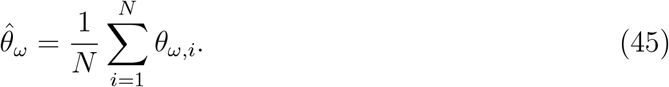

The overall mean across all chains is

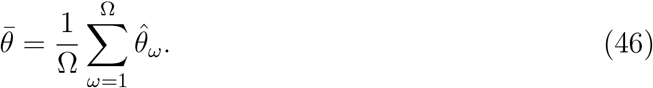

The between-chain variance is

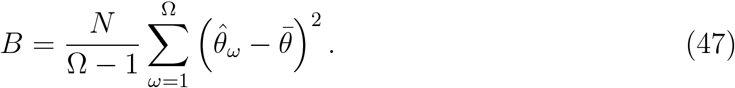

The within-chain variance for each chain is

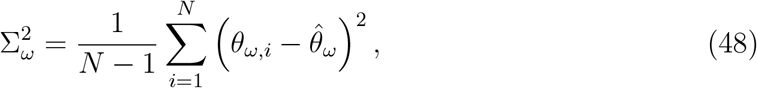

and the average within-chain variance is

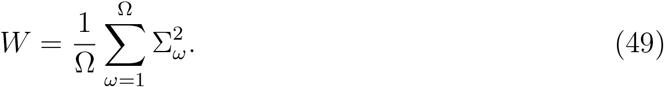

The pooled variance estimator is defined as

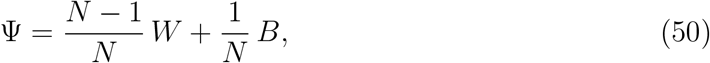

and the Gelman–Rubin statistic is then computed as

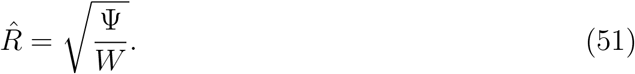

In practice, values of 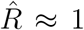 (e.g., *<* 1.1) indicate acceptable convergence, whereas 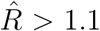 suggests potential convergence issues [49]. In this study, we require 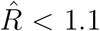 for all parameters.

### 3.9 Free-Energy Calculations for SLC25A10 Antiport processes

The free-energy change (Δ*G*) for the electroneutral antiport processes mediated by the SLC25A10 transporter is calculated for the following exchange reactions:

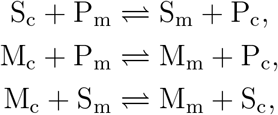

following the methodology of Jol et al. (2010) [51]. The standard chemical-thermodynamics expression for the free-energy change is:

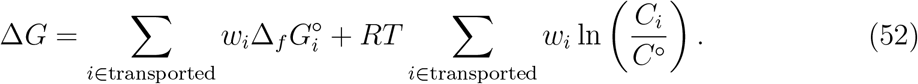

Here, *w*_*i*_ are the stoichiometric coefficients for the transported species (e.g. *S*_*m*_, *P*_*m*_, *S*_*c*_, *P*_*c*_), with values of −1 for reactants on the matrix side and +1 for products on the external side. For each transported species *i*, 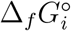 denotes the standard Gibbs energy of formation, and *C*_*i*_ denotes its concentration. The logarithmic terms are defined relative to the standard concentration *C*^◦^, taken here as 1 M [52], so that ln(*C*_*i*_*/C*^◦^) is dimensionless. Moreover, *R* is the gas constant, and *T* is the absolute temperature.

For succinate exchange, this yields

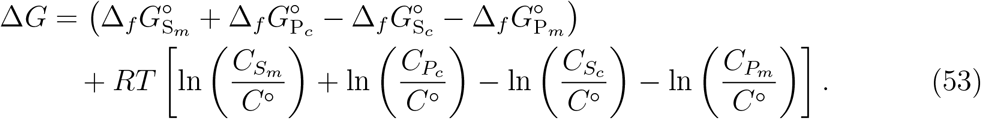

Using our notation, with [*S*_*m*_], [*S*_*c*_], [*P*_*m*_], [*P*_*c*_] in place of concentrations *C*, we write

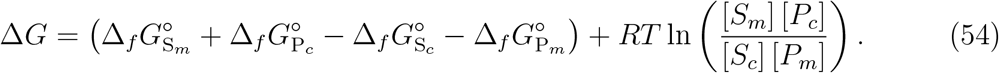

Similarly, for malate exchange,

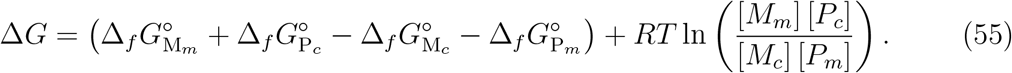

For malate/succinate exchange, the corresponding reaction free-energy change is

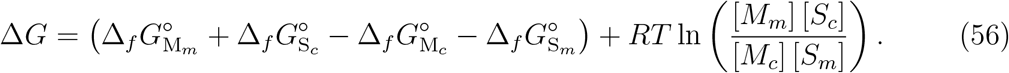

At thermodynamic equilibrium, Δ*G* = 0, giving

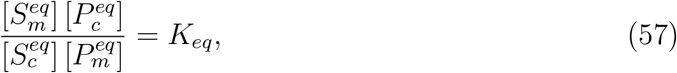

where 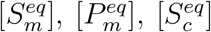, and 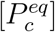 are the equilibrium concentrations of matrix and external succinate and phosphate, respectively. The equilibrium constant (*K*_*eq*_) is given by

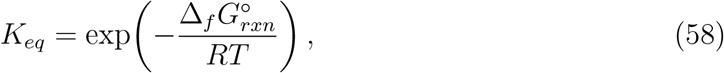

where

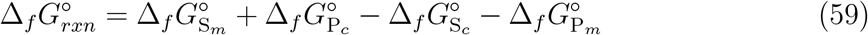

is the standard reaction free-energy change. Since the species are chemically identical across compartments, 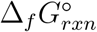 is zero, and the equilibrium distribution is dominated by the concentration ratios.

From conservation of mass in each metabolite pool, the analytical equilibrium concentrations can be written as explicit functions of initial metabolite concentrations. Let the initial mitochondrial concentrations for succinate, and phosphate be 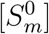, and 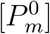, respectively, and let the external initial concentrations be 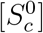, and 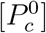, respectively. Then, the resulting closed-form equilibrium concentrations are

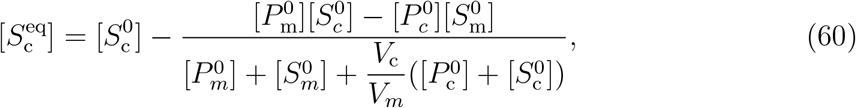

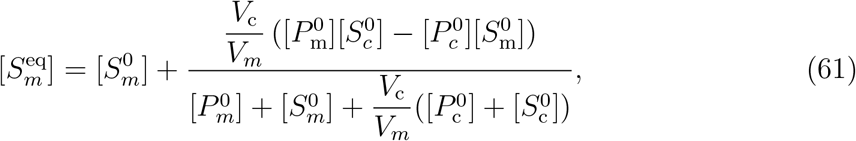

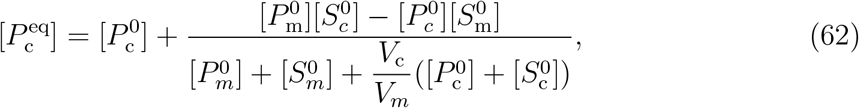

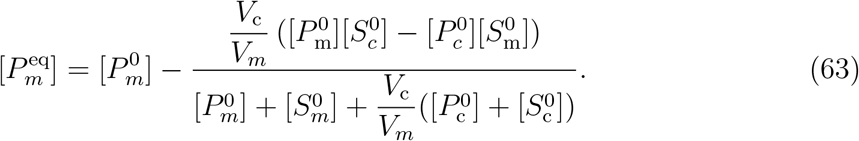

These closed-form expressions allow direct analytical validation of the simulated equilibrium state of the reduced transporter model against thermodynamic constraints. The full derivation of these expressions is provided in Appendix E.

## 4 Results

### 4.1 Local Sensitivity Analysis

We investigated how the kinetic parameters influence the dynamics of the SLC25A10 transporter. To this end, we computed normalised local sensitivity coefficients (LSCs) for three key output fluxes: succinate (*J*_suc_), malate (*J*_mal_), and phosphate (*J*_pho_). Positive values indicate that an increase in the parameter enhances the flux, while negative values indicate suppression.

The analysis was performed on the transporter model with six kinetic parameters: the maximum transport rates for succinate 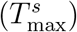 and malate 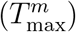, and phosphate 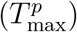; the dissociation constants for succinate 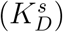, malate 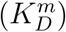, and phosphate 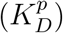.

Parameter values were initialised according to Table 1. External compartment metabolite concentrations were set to match the experimental data of Palmieri et al. [3], specifically the dataset measuring malate uptake in the presence of 0.5 mM external succinate. This condition provides a direct test of competitive inhibition. Here, the external malate concentration was fixed at 0.9077 mM, external succinate at 0.5 mM, and other conditions matched the experimental setup.

We compare finite-difference and complex-step sensitivity coefficients as a numerical accuracy check by reporting the absolute and relative differences between the two estimates (see Appendix H). The results show close agreement across all parameters (maximum discrepancy *<* 6 × 10^−4^), supporting the numerical reliability of our finite-difference implementation for the chosen perturbation size.

For the succinate flux (Fig. 3a), the dominant positive sensitivity is to the maximal succinate transport capacity 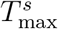, as expected for a capacity-controlled contribution. A strong positive sensitivity is also observed for 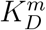, indicating that weaker malate binding reduces competitive inhibition and thereby increases *J*_suc_. By contrast, the strongest negative sensitivity is to 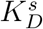, showing that reduced succinate affinity substantially suppresses succinate flux. The malate transport capacity 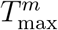 and phosphate dissociation constant 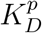 also contribute negatively, whereas 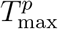 has a moderate positive influence. Overall, *J*_suc_ is governed primarily by succinate-side capacity and affinity, with additional modulation from substrate competition and phosphate-coupled exchange.

**Figure 3:**
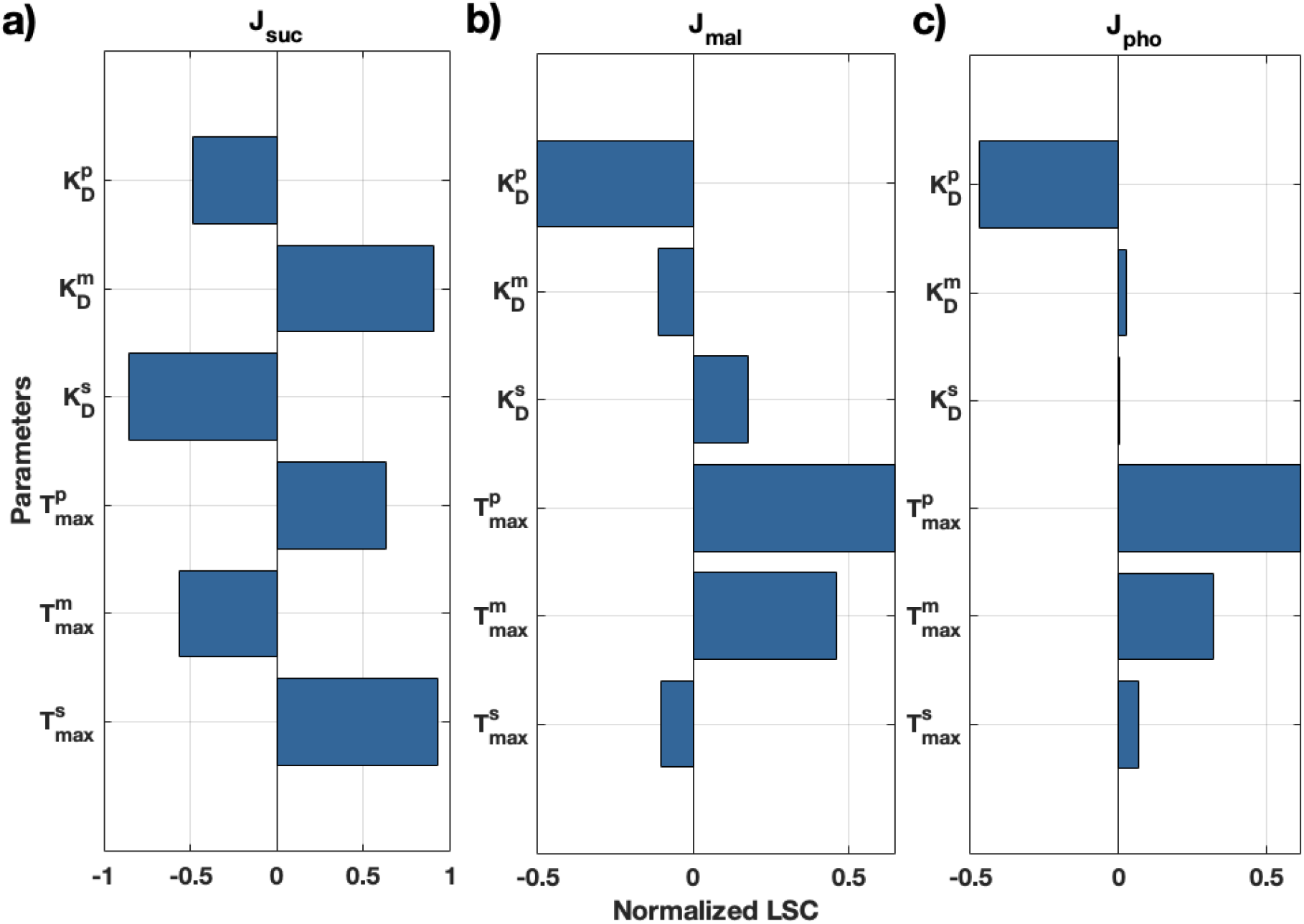
**Normalised local sensitivity coefficients (LSCs)** for (a) succinate flux *J*_suc_, (b) malate flux *J*_mal_, and (c) phosphate flux *J*_pho_ with respect to model parameters. Here, “normalised” means that each sensitivity is scaled relative to both the value of the parameter and the corresponding flux, making the coefficients dimensionless. This allows the influence of parameters with different units and magnitudes to be compared on the same scale. Positive values indicate that increasing a parameter increases the fluxes, whereas negative values imply the opposite.

For the malate flux (Fig. 3b), the largest positive sensitivity is to 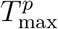, followed by 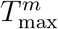, indicating that malate turnover in this regime is strongly coupled to phosphate exchange and to the intrinsic malate transport capacity. The dominant negative contribution arises from 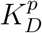, showing that weaker phosphate affinity substantially reduces *J*_mal_. Smaller effects are observed from the remaining parameters: 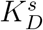 contributes weakly and positively, while 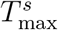 and 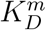 contribute weakly and negatively. These results indicate that, under the chosen assay conditions, malate flux is influenced most strongly by the phosphate-coupled branch of the antiport cycle.

For the phosphate flux (Fig. 3c), 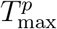 is the strongest positive driver, with 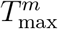 providing the next largest positive contribution. The dominant negative sensitivity is to 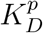, confirming that phosphate affinity is a key determinant of phosphate turnover in the antiport cycle. The effects of 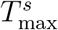 and 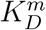 are comparatively small but positive, whereas is negligible in this setting. Taken together, these results show that phosphate flux is controlled primarily by the phosphate transport capacity and affinity, while still reflecting coupling to the malate branch of the transporter cycle.

### 4.2 Bayesian Parameter Estimation

Following the structural identifiability and local sensitivity analyses, we employed a Bayesian inference framework to quantitatively estimate the kinetic parameters and assess the uncertainty of the SLC25A10 transporter model. The model was then validated against experimental datasets from both reconstituted proteoliposomes and intact rat liver mitochondria.

We ran 1,000,000 Metropolis-Hastings MCMC iterations, discarding the first 10,000 as burn-in to reduce dependence on the initial guess. Posterior means, standard deviations, modes, and 95% credible intervals are computed from the retained samples for all parameters, including *σ*^2^, for each uptake-assay dataset. A statistical summary of the posterior parameter estimates for the intact mitochondria calibration is provided in Table 4. The corresponding posterior summary for the proteoliposome calibration is reported separately in Table 5.

**Table 3:** Thermodynamic parameters and constants for reaction free-energy calculations of SLC25A10 antiport reactions.

| Species / Constant | Charge ($z$) | $\Delta_f G^\circ$ (kJ/mol) | $w_i^{M/P}$ | $w_i^{S/P}$ | $w_i^{M/S}$ | Reference |
| --- | --- | --- | --- | --- | --- | --- |
| External Malate ($M_c$) | -2 | -842.66 | -1 | 0 | -1 | [17, 53] |
| Matrix Malate ($M_m$) | -2 | -842.66 | +1 | 0 | +1 | [17, 53] |
| External Succinate ($S_c$) | -2 | -690.44 | 0 | -1 | +1 | [51, 53] |
| Matrix Succinate ($S_m$) | -2 | -690.44 | 0 | +1 | -1 | [51, 53] |
| External Phosphate ($P_c$) | -2 | -1069.1 | +1 | +1 | 0 | [51, 53] |
| Matrix Phosphate ($P_m$) | -2 | -1069.1 | -1 | -1 | 0 | [51, 53] |
| Gas Constant (R) | - | 0.008314 kJ/K | - | - | - | [51] |
| Temperature (T) | - | 298.15 K (25 °C) | - | - | - | [51] |

**Table 4:** Statistical summary of posterior parameter estimates inferred from isolated intact mitochondria uptake assays, including 95% credible intervals.

| Parameter | Mean | Median | Mode | Std | 95% Credible Interval |  |
| --- | --- | --- | --- | --- | --- | --- |
|  |  |  |  |  | Low | High |
| $T_{\max}^s$ | 146.4 | 135.68 | 115.53 | 53.499 | 74.874 | 284.46 |
| $T_{\max}^m$ | 216.98 | 156.5 | 112.99 | 170.42 | 76.444 | 743.99 |
| $T_{\max}^p$ | 304.24 | 227.36 | 130.82 | 213.05 | 82.389 | 850.74 |
| $K_D^s$ | 2.4566 | 2.2352 | 1.8071 | 1.0553 | 1.084 | 5.2137 |
| $K_D^m$ | 0.38049 | 0.31821 | 0.26966 | 0.19167 | 0.19224 | 0.95612 |
| $K_D^p$ | 2.4324 | 2.139 | 1.5845 | 1.1712 | 0.97533 | 5.4296 |
| $\lambda_{21}$ (derived) | 8.5505 | 8.1676 | 7.5545 | 2.091 | 5.644 | 13.456 |
| $\lambda_{31}$ (derived) | 1.9137 | 1.7405 | 1.4680 | 0.64385 | 1.1562 | 3.5359 |
| $\sigma^2$ | 1.1447 | 1.0411 | 1.0509 | 0.46105 | 0.56415 | 2.301 |

**Table 5:** Statistical summary of posterior parameter estimates inferred from proteoliposome transport assays, including 95% credible intervals.

| Parameter | Mean | Median | Mode | Std | 95% Credible Interval |  |
| --- | --- | --- | --- | --- | --- | --- |
|  |  |  |  |  | Low | High |
| $T_{\max}^m$ | 9.0682 | 9.0117 | 8.9317 | 1.6217 | 6.0510 | 12.397 |
| $T_{\max}^p$ | 8.3548 | 8.2754 | 7.8945 | 1.6590 | 5.3364 | 11.816 |
| $K_D^m$ | 0.47927 | 0.47930 | 0.48180 | 0.049438 | 0.38219 | 0.57606 |
| $K_D^p$ | 1.3709 | 1.3667 | 1.3530 | 0.32503 | 0.74742 | 2.0224 |
| $\lambda_{23}$ (derived) | 0.35364 | 0.32545 | 0.28488 | 0.14516 | 0.16236 | 0.70920 |
| $\sigma^2$ | 1.5799 | 1.5288 | 1.4173 | 0.36543 | 1.0194 | 2.4427 |

For convergence diagnostics, we ran four independent MCMC chains initialised from dispersed starting values, each for 1,000,000 iterations with the first 10,000 discarded as burn-in for both uptake-assay calibrations. For each parameter, we computed the split-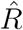 statistic using the post–burn-in draws across chains. All parameters satisfied 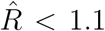. Table 6 shows the MCMC diagnostics for intact mitochondria assays, with values close to 1.0 (maximum 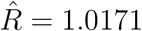). Table 7 shows the MCMC diagnostics for the proteoliposome calibration, with values very close to 1.0 (maximum 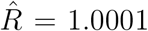). Both tables indicate good mixing and strong evidence of convergence to a common stationary posterior. The Metropolis–Hastings transition kernel used in our sampler satisfies detailed balance with respect to the posterior target by construction; the small 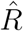 values confirm that the chains reached this stationary regime in practice.

**Table 6:**
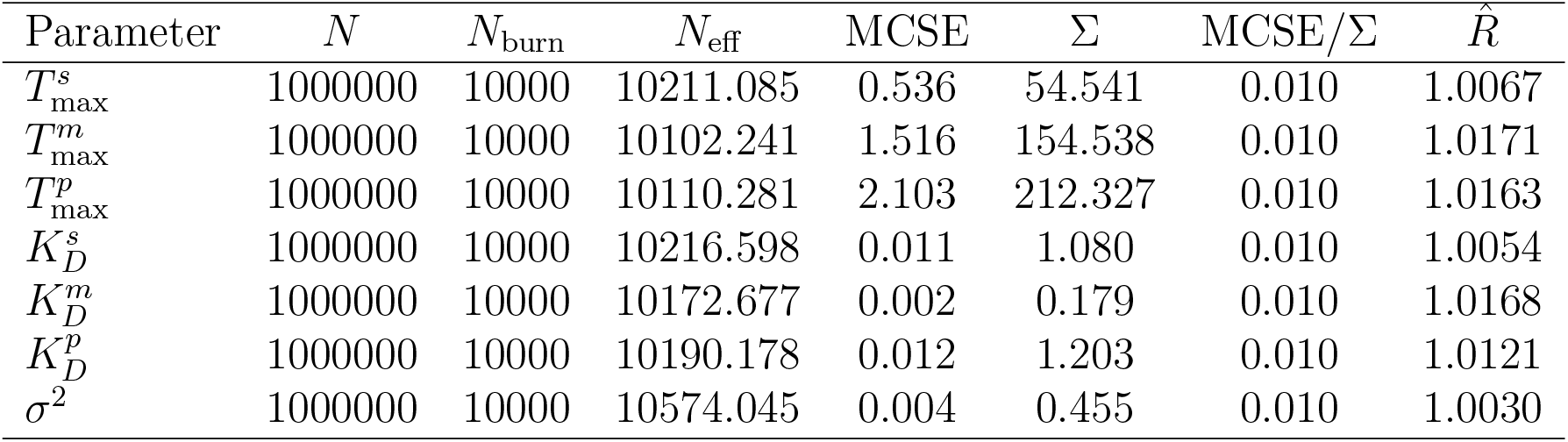
MCMC diagnostics for the intact mitochondria uptake-assay calibration. Diagnostics are reported per parameter with respect to total draws (*N* ), burn-in draws (*N*_burn_), effective sample size (*N*_eff_), Monte Carlo standard error (MCSE), posterior standard deviation (Σ), relative Monte Carlo error (MCSE/Σ), and 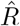.

**Table 7:** MCMC diagnostics for the proteoliposome uptake-assay calibration. Diagnostics are reported per parameter with respect to total draws (*N* ), burn-in draws (*N*_burn_), effective sample size (*N*_eff_), Monte Carlo standard error (MCSE), posterior standard deviation (Σ), relative Monte Carlo error (MCSE/Σ), and 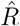.

| Parameter | $N$ | $N_{\text{burn}}$ | $N_{\text{eff}}$ | MCSE | $\Sigma$ | MCSE/$\Sigma$ | $\hat{R}$ |
| --- | --- | --- | --- | --- | --- | --- | --- |
| $T_{\max}^m$ | 1000000 | 10000 | 31012.386 | 0.012 | 2.164 | 0.006 | 1.0000 |
| $T_{\max}^p$ | 1000000 | 10000 | 31138.646 | 0.009 | 1.654 | 0.006 | 1.0000 |
| $K_D^m$ | 1000000 | 10000 | 92380.352 | 0.000 | 0.049 | 0.003 | 1.0000 |
| $K_D^p$ | 1000000 | 10000 | 26090.371 | 0.002 | 0.325 | 0.006 | 1.0001 |
| $\sigma^2$ | 1000000 | 10000 | 30184.533 | 0.002 | 0.403 | 0.006 | 1.0000 |

**Table 8:** State variables in the SLC25A10 transport model.

| Symbol | Description | Units |
| --- | --- | --- |
| $S_m$ | Succinate concentration in matrix | $\text{mol L}^{-1}$ |
| $M_m$ | Malate concentration in matrix | $\text{mol L}^{-1}$ |
| $P_m$ | Phosphate concentration in matrix | $\text{mol L}^{-1}$ |
| $S_c$ | Succinate concentration in external compartment | $\text{mol L}^{-1}$ |
| $M_c$ | Malate concentration in external compartment | $\text{mol L}^{-1}$ |
| $P_c$ | Phosphate concentration in external compartment | $\text{mol L}^{-1}$ |
| $T_m$ | Free transporter facing matrix | $\text{mol g}^{-1}$ |
| $T_c$ | Free transporter facing external compartment | $\text{mol g}^{-1}$ |
| $T_m S$ | Transporter–succinate complex facing matrix | $\text{mol g}^{-1}$ |
| $T_m M$ | Transporter–malate complex facing matrix | $\text{mol g}^{-1}$ |
| $T_c S$ | Transporter–succinate complex facing external side | $\text{mol g}^{-1}$ |
| $T_c M$ | Transporter–malate complex facing external side | $\text{mol g}^{-1}$ |
| $T_c P$ | Transporter–phosphate complex facing external side | $\text{mol g}^{-1}$ |
| $T_m P$ | Transporter–phosphate complex facing matrix | $\text{mol g}^{-1}$ |
| $T_{\text{total}}$ | Total transporter amount per gram protein | $\text{mol g}^{-1}$ |

*Note:* $L$ denotes liters, $g$ denotes grams of protein, and $\text{mol}$ denotes moles.

For the intact-mitochondria calibration, the effective sample sizes (*N*_eff_) ranged from 10,117.6 to 10,608.1 across the sampled parameters and derived coupling ratios. MCSE/Σ ratios were approximately 0.01 for all reported quantities, indicating that Monte Carlo error was small relative to posterior uncertainty. For the proteoliposome calibration, *N*_eff_ ranged from 26,090.4 to 92,380.4, and MCSE/Σ ratios ranged from 0.003 to 0.006. All parameters meet the conventional benchmark MCSE/Σ *<* 0.10, suggesting that Monte Carlo variability contributes negligibly to posterior uncertainty. Overall, these diagnostics indicate that the posterior summaries for both the intact-mitochondria and proteoliposome calibrations are numerically stable and suitable for inference.

Figures 4 and 5 summarise the MCMC trace plots and posterior distributions for parameters inferred from the intact-mitochondria uptake-assay calibration and the proteoliposome calibration, respectively. In both cases, the trace plots indicate stable exploration of the posterior after burn-in, and the marginal densities highlight which parameters are well constrained by the available data.

**Figure 4:**
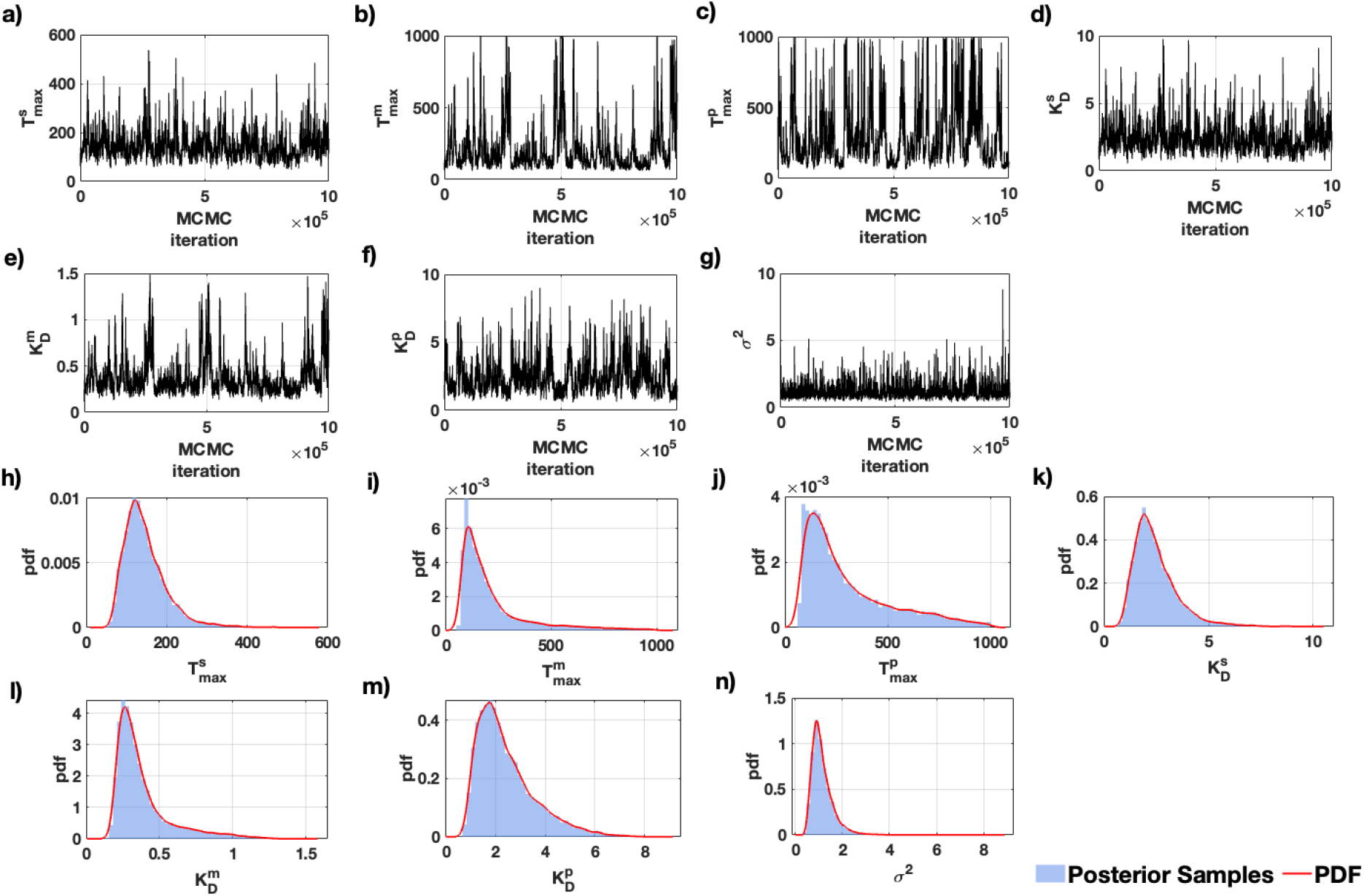
MCMC posterior summaries for the intact mitochondria uptake-assay calibration. Trace plots of the sampled parameter chains are shown in panels (a–g): (a) 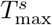, (b) 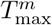, (c) 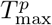, (d) 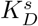, (e) 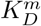, (f) 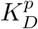, and (g) *σ*^2^. Posterior density summaries for the same parameters are shown in panels (h–n). In the density panels, the blue histograms denote posterior samples and the red curves denote kernel density estimates (KDEs) used to provide smoothed posterior summaries for all parameters.

**Figure 5:**
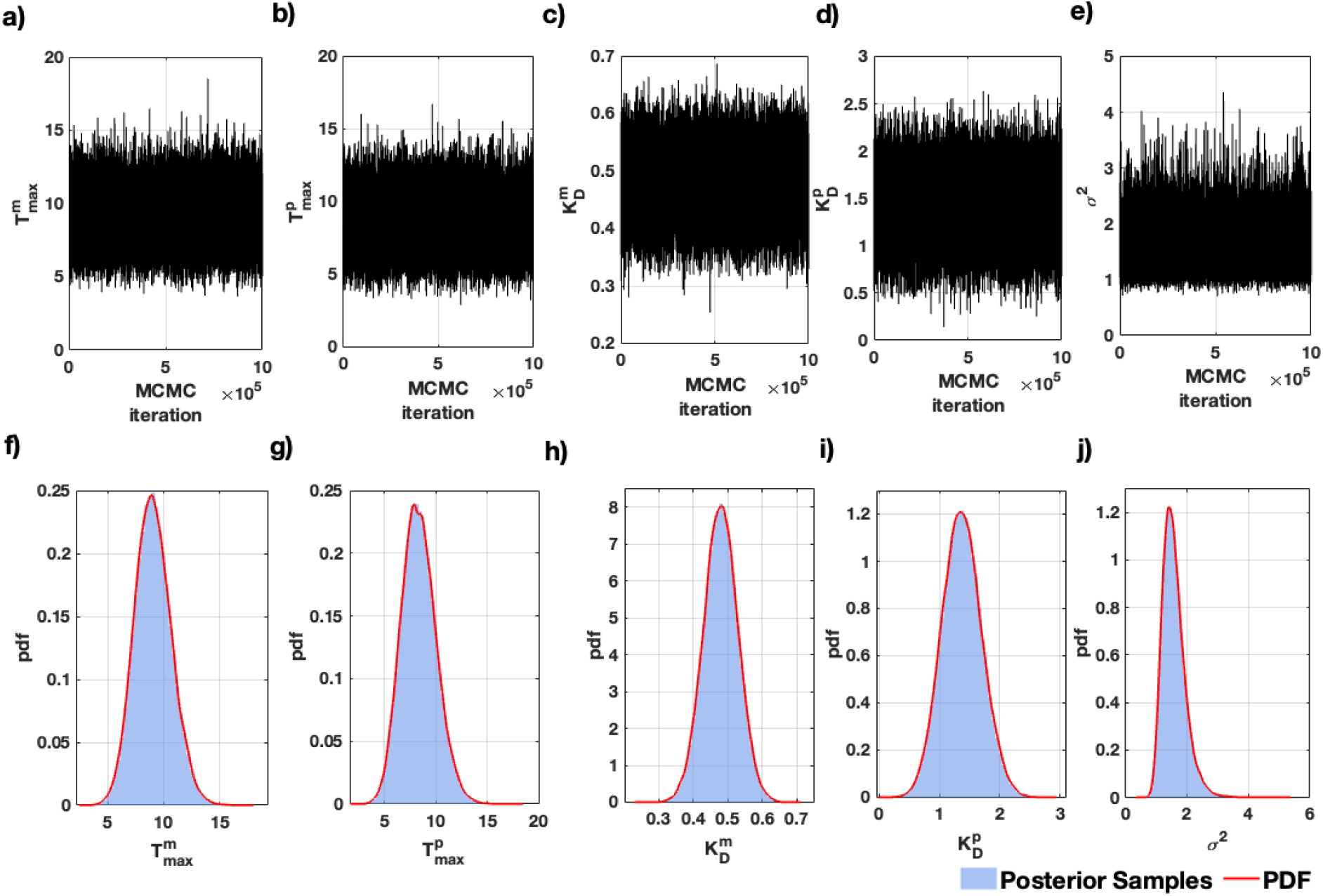
MCMC posterior summaries for the proteoliposome calibration. Trace plots of the sampled parameter chains are shown in panels (a–e): (a) 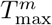, (b) 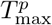, (c) 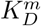, (d) 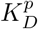, and (e) *σ*^2^. Posterior density summaries for the same parameters are shown in panels (f–j). In the density panels, the blue histograms denote posterior samples and the red curves denote kernel density estimates (KDEs) used to provide smoothed posterior summaries for all parameters.

For the intact-mitochondria dataset (Fig. 4), the reduced posterior supports a broad but physiologically coherent range of transport capacities and affinities. The inferred means are 146.4, 216.98, and 304.24 *µ*mol min^−1^ g^−1^ for 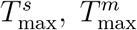, and 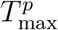, respectively, and 2.4566, 0.38049, and 2.4324 mM for 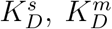, and 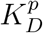, respectively. Thus, under the intact-mitochondria uptake conditions, the posterior favors a high-capacity malate/phosphate branch and a comparatively low 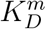, consistent with strong malate binding relative to succinate and phosphate. The derived coupling ratios have posterior means *λ*_21_ = 8.5505 and *λ*_31_ = 1.9137, indicating that malate contributes more strongly than phosphate to the conformational-bias term relative to succinate under this calibration. Because these ratios are derived from the kinetic parameters, their uncertainty reflects propagated uncertainty from the reduced posterior rather than independent sampling.

For the proteoliposome dataset (Fig. 5), the posterior is more tightly concentrated around the literature-informed kinetic priors and the reconstituted transport measurements. The posterior means are 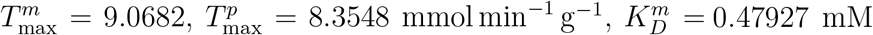, and 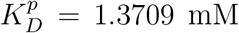. The derived ratio *λ*_23_ = 0.35364 indicates that, in the malate–phosphate proteoliposome setting with succinate absent, the identifiable phosphate/malate weighting is below one. This is consistent with the posterior estimate 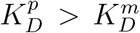, meaning that phosphate-related terms remain less strongly constrained by affinity than the malate branch. The observation-noise posterior for *σ*^2^ remains well behaved in both calibrations, supporting a stable Gaussian error model for the fitted uptake rates.

Overall, the Bayesian calibration estimates transport capacities and dissociation constants as the primary kinetic parameters, while substrate-weighting quantities are computed from these estimates according to the mechanistic relations. This parameterisation preserves the mechanistic interpretation of substrate weighting while linking it directly to the estimated transport capacities and dissociation constants, providing a consistent basis for posterior summaries and downstream simulations.

Figures 6 and 7 compare the prior and posterior distributions for the inferred parameters under the intact-mitochondria and proteoliposome calibrations, respectively. In the intact-mitochondria uptake assays (Fig. 6), the posterior distributions contract strongly relative to the uniform priors for all six inferred kinetic parameters, namely 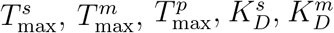, and 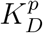. This indicates that the intact-mitochondria dataset is informative for constraining both transport-capacity and affinity parameters in the reduced model. In the proteoliposome setting (Fig. 7), inference is restricted to the malate–phosphate branch, and the posterior distributions for 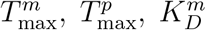, and 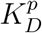 show varying degrees of departure from their Gaussian priors. The clearest posterior shift is observed for 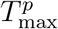, whereas 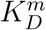 and 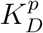 remain more strongly influenced by the prior specification. In both calibrations, the posterior distribution of *σ*^2^ concentrates toward relatively small values, supporting a low inferred model–data discrepancy under the assumed Gaussian noise model.

**Figure 6:**
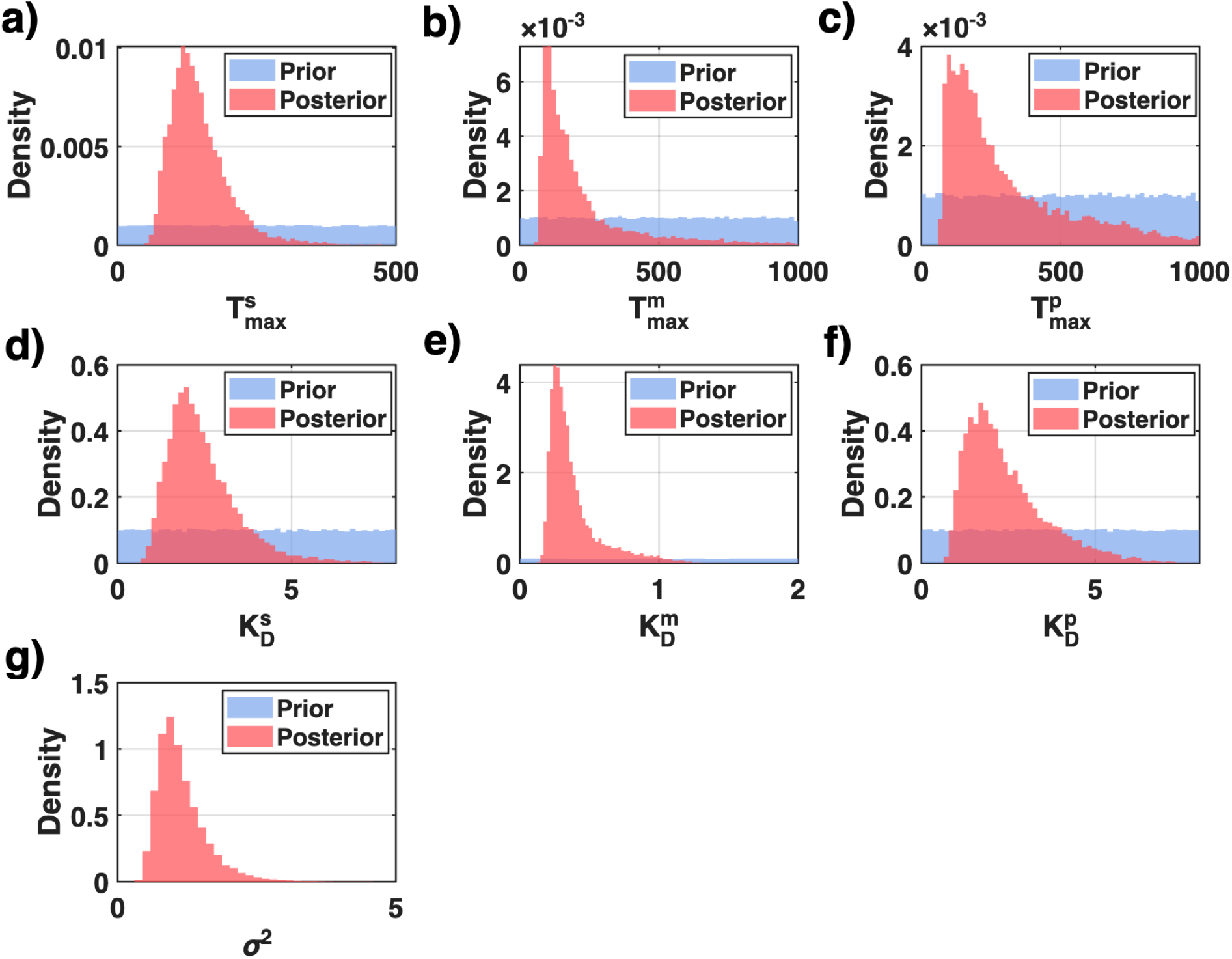
Prior–posterior comparison for the intact-mitochondria uptake-assay calibration. Overlaid density plots compare the prior (blue) and posterior (red) distributions for all inferred model parameters. Panels (a–c) show the maximal transport capacities 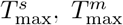, and 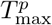; panels (d–f) show the dissociation constants 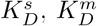, and 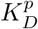; and panel (g) shows the observation-noise variance *σ*^2^. Uniform priors were used for the six kinetic parameters, and an inverse-gamma prior was used for *σ*^2^.

**Figure 7:**
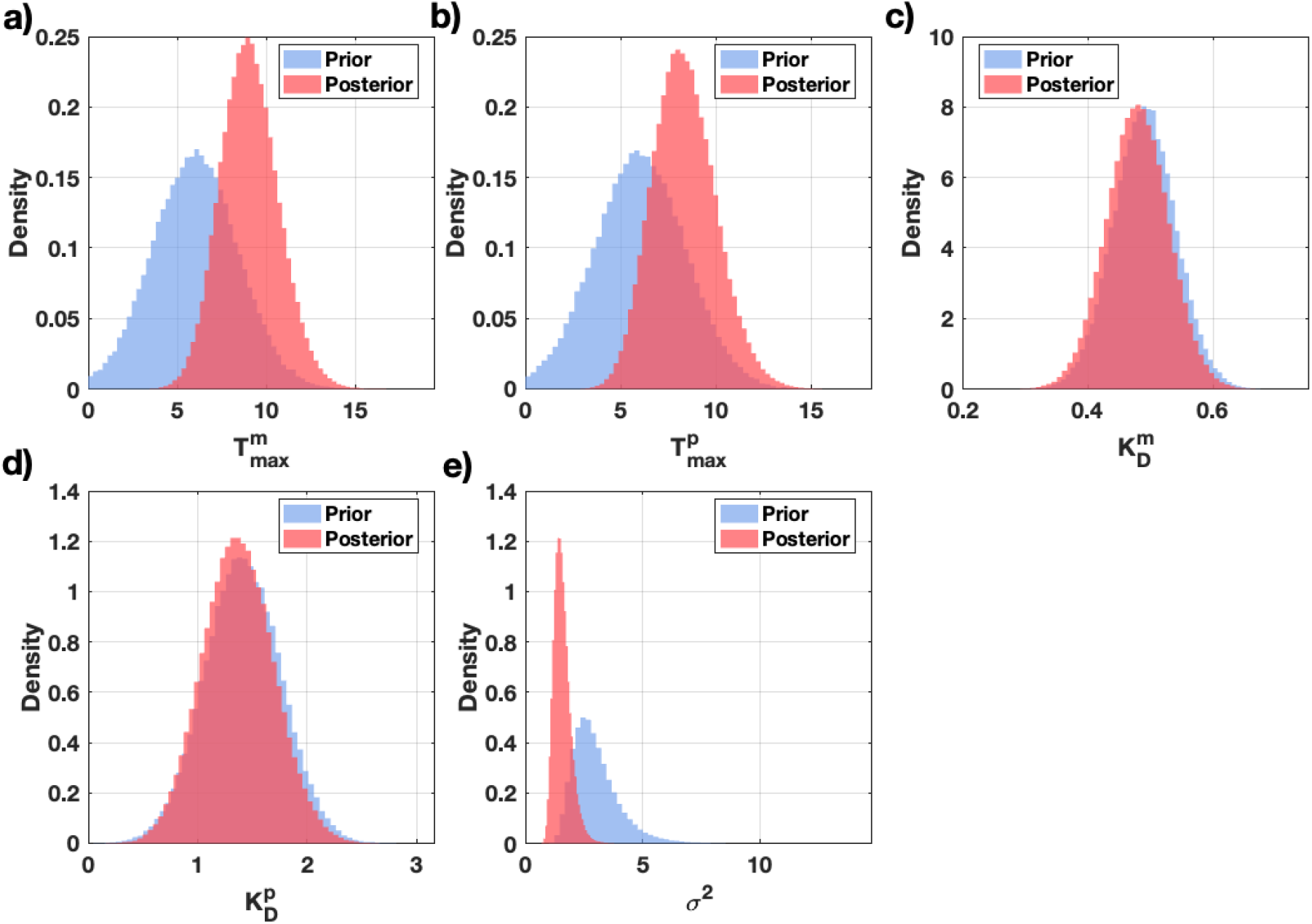
Prior–posterior comparison for the proteoliposome calibration. Overlaid density plots compare the prior (blue) and posterior (red) distributions for the inferred model parameters. Panels (a–b) show the maximal transport capacities 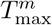 and 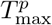; panels (c–d) show the dissociation constants 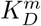 and 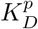; and panel (e) shows the observation-noise variance *σ* . Gaussian priors were used for the four kinetic parameters, and an inverse-gamma prior was used for *σ*^2^.

Figure 8 compares Bayesian model predictions (posterior mean and 95% credible intervals) against the experimental uptake-assay flux measurements. For the intact-mitochondria datasets (Fig. 8a–c), the inferred model closely tracks the observed trends across all tested conditions. Specifically, succinate uptake *J*_suc_ (Fig. 8a) increases monotonically with external succinate [*S*_*c*_], and the posterior mean follows the experimental curve with all points lying within the 95% credible band, indicating that the calibrated transport capacity and succinate affinity are well constrained. Malate uptake *J*_mal_ under competitive conditions (Fig. 8b), with external succinate fixed at 0.5 mM, exhibits the expected increasing dependence on external malate concentration [*M*_*c*_], and the model reproduces this behaviour well, indicating that the inferred reduced ping–pong formulation captures substrate competition consistently. Phosphate uptake *J*_pho_ (Fig. 8c) increases with external phosphate concentration [*P*_*c*_], and although the uncertainty band is somewhat broader at higher concentrations, the model still captures the overall trend and magnitude of the measurements.

**Figure 8:**
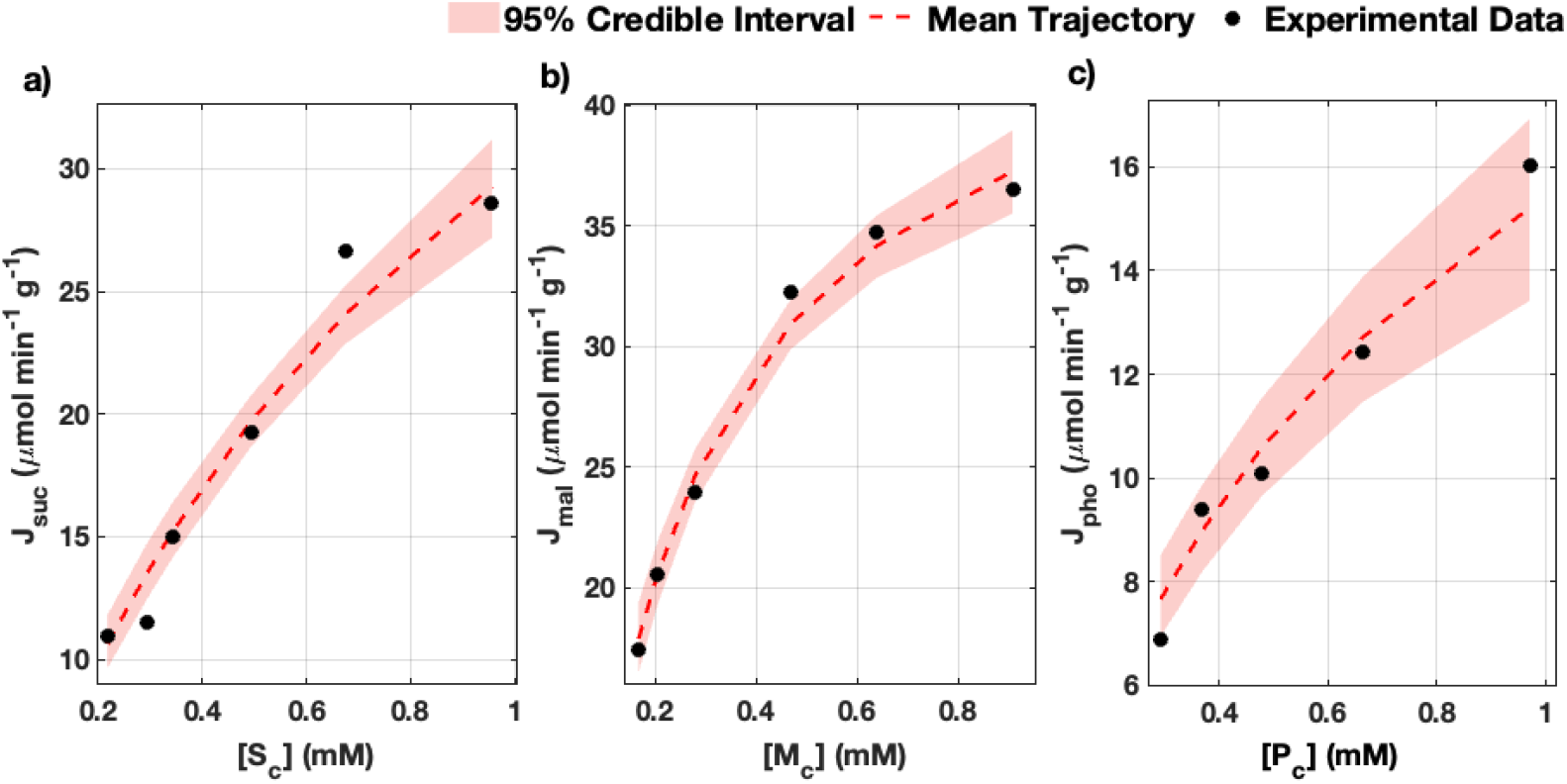
Model validation against isolated intact-mitochondria uptake assays. Experimental uptake fluxes (black dots) are compared with the Bayesian model fit, shown as the posterior mean trajectory (red dashed line) with a 95% credible interval (shaded band). (a) Succinate uptake flux *J*_suc_ as a function of external succinate concentration [*S*_*c*_], with external malate and phosphate held at zero. (b) Malate uptake flux *J*_mal_ as a function of [*M*_*c*_] under competitive conditions, with external succinate fixed at 0.5 mM and external phosphate set to zero. (c) Phosphate uptake flux *J*_pho_ as a function of external phosphate concentration [*P*_*c*_] under non-competitive conditions (no external succinate or malate).

The proteoliposome validation experiments (Fig. 9a–e) show similarly strong agreement between model and data under multiple assay configurations. The model reproduces malate flux *J*_mal_ as a function of external malate concentration [*M*_*c*_] for several fixed intravesicular phosphate conditions (Fig. 9a, 9b, 9e), demonstrating that the model captures how internal phosphate availability modulates malate uptake in the reconstituted system. It also reproduces the dependence of *J*_mal_ on intravesicular phosphate [*P*_*m*_] at fixed external malate concentration (Fig. 9c), which is consistent with the expected phosphate-coupled antiport behaviour. Finally, the model matches the phosphate uptake data *J*_pho_ as a function of external phosphate concentration [*P*_*c*_] at fixed intravesicular malate (Fig. 9d), providing complementary support for the inferred phosphate-linked exchange step.

**Figure 9:**
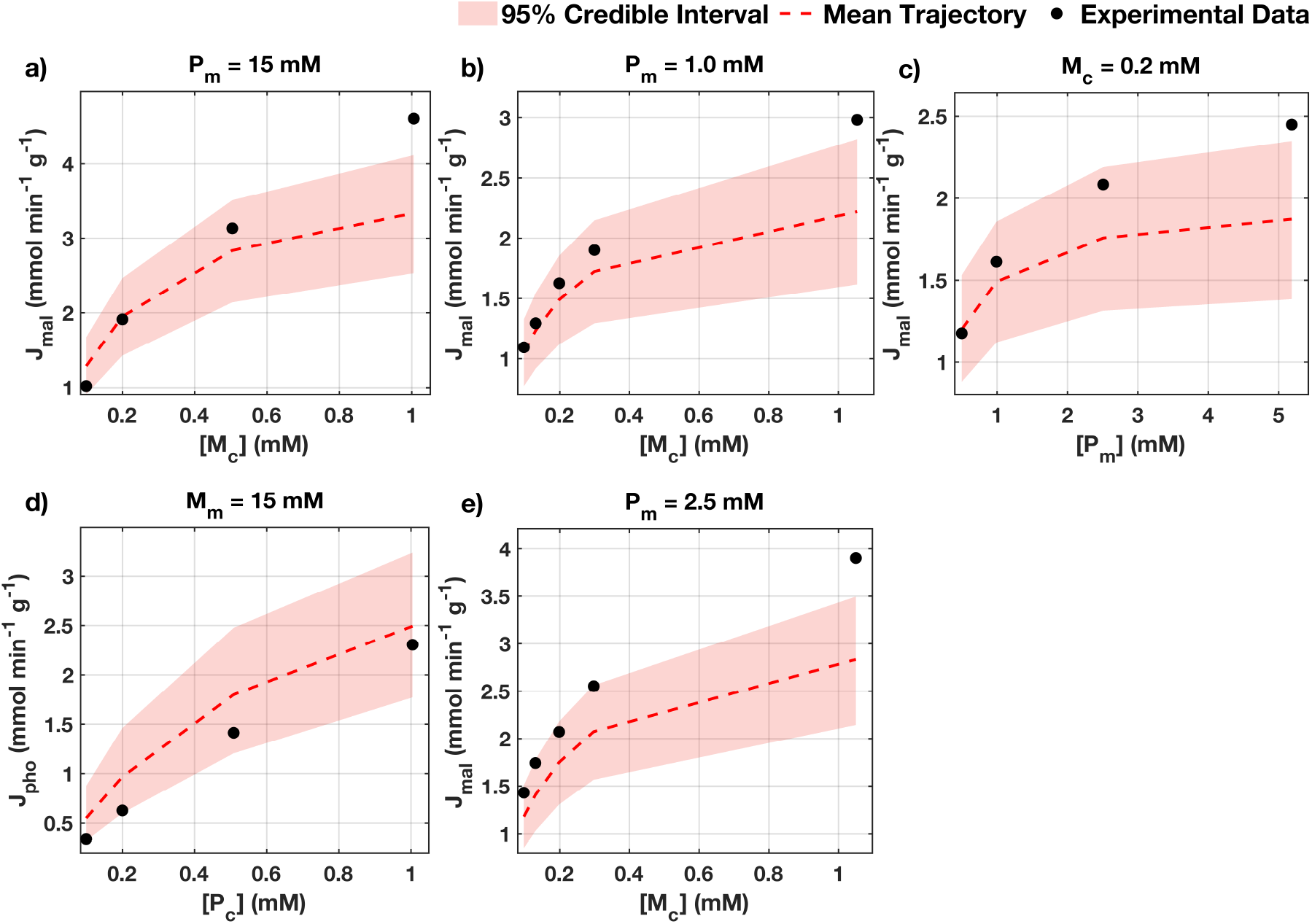
Model validation against proteoliposome uptake assays. Experimental flux measurements (black dots) are compared with the Bayesian model fit, shown as the posterior mean trajectory (red dashed line) with a 95% credible interval (shaded band). (a) Malate uptake flux *J*_mal_ as a function of external malate [*M*_*c*_] with intravesicular phosphate fixed at [*P*_*m*_] = 15 mM. (b) *J*_mal_ versus [*M*_*c*_] with [*P*_*m*_] = 1.0 mM. (c) *J*_mal_ versus intravesicular phosphate [*P*_*m*_] with external malate fixed at [*M*_*c*_] = 0.2 mM. (d) Phosphate uptake flux *J*_pho_ as a function of external phosphate [*P*_*c*_] with intravesicular malate fixed at [*M*_*m*_] = 15 mM. (e) *J*_mal_ versus [*M*_*c*_] with [*P*_*m*_] = 2.5 mM.

Overall, across both intact mitochondria and proteoliposome settings, the posterior predictive bands generally enclose the experimental measurements and the posterior mean trajectories reproduce the observed saturation and competitive effects. These results support that the Bayesian-calibrated model provides a quantitatively accurate and mechanistically consistent representation of SLC25A10-mediated dicarboxylate/phosphate exchange under the experimental assay conditions.

### 4.3 Thermodynamic Validation of the Kinetic Model

To ensure that the SLC25A10 model is not only mathematically consistent but also thermodynamically sound, we validated it by analysing flux trajectories, exchange-reaction free-energy dissipation, and equilibrium metabolite concentrations under defined initial conditions in both calibration settings.

Figures 10 and 11 show that the model satisfies thermodynamic constraints and reproduces the analytically predicted equilibrium partitioning between compartments. In the intact-mitochondria setting (Fig. 10), the antiport nature of the mechanism is reflected by the opposing signs of the succinate and phosphate fluxes, *J*_suc_ and *J*_pho_, which relax to zero as concentration gradients dissipate (Fig. 10a). Consistently, the exchange-reaction free-energy change Δ*G* increases toward zero and asymptotically approaches Δ*G* = 0, indicating relaxation to thermodynamic equilibrium (Fig. 10b). Simulated equilibrium concentrations for succinate and phosphate in the matrix and external medium agree closely with the closed-form equilibrium expressions derived from the reduced mass-balance formulation (Fig. 10c–f), providing an independent check that the reduced and full formulations are mutually consistent.

**Figure 10:**
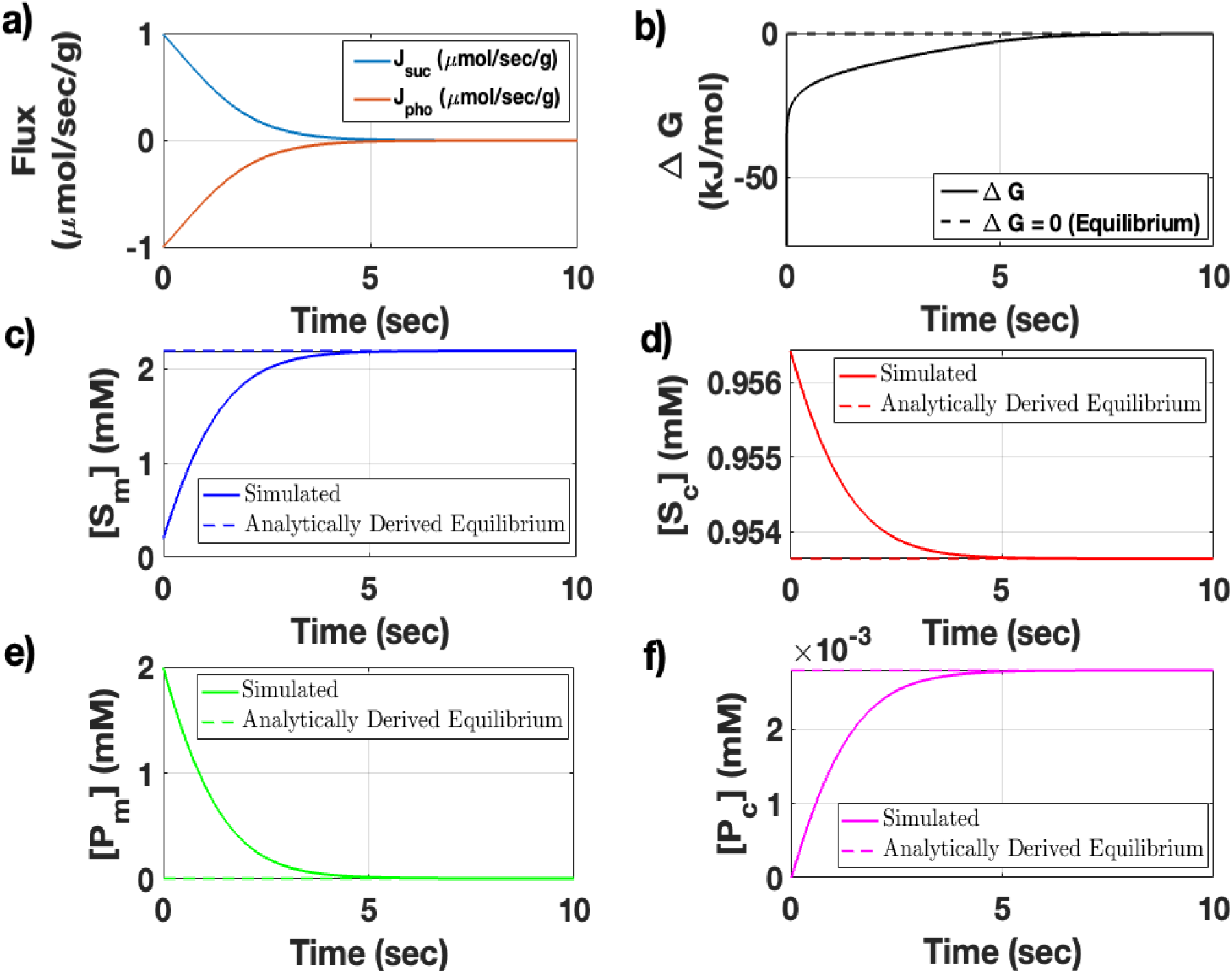
Thermodynamic analysis and equilibrium validation of the SLC25A10 antiport model in the intact mitochondria setting. (a) Time-dependent fluxes of succinate (*J*_suc_) and phosphate (*J*_pho_) showing relaxation toward equilibrium under electroneutral exchange stoichiometry. (b) Time-resolved free energy change (Δ*G*) for the succinate/phosphate exchange reaction S_*c*_+P_*m*_ ⇌ S_*m*_+P_*c*_; the dashed line indicates Δ*G* = 0 (thermodynamic equilibrium). (c–f) Simulated concentrations of succinate and phosphate in the matrix (*m*) and the external compartment (*c*; assay medium) compared with analytically derived equilibrium values (dashed lines). Initial conditions are [*S*_*c*_] = 0.1 mM and [*P*_*c*_] = 0 mM, with malate set to zero in both compartments; remaining initial concentrations are set according to Table 1. Simulations use the intact-mitochondria compartment volumes (per gram protein) reported in Table 1. Parameter values for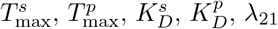, and *λ*_31_ are fixed at their posterior modes (Table 4).

**Figure 11:**
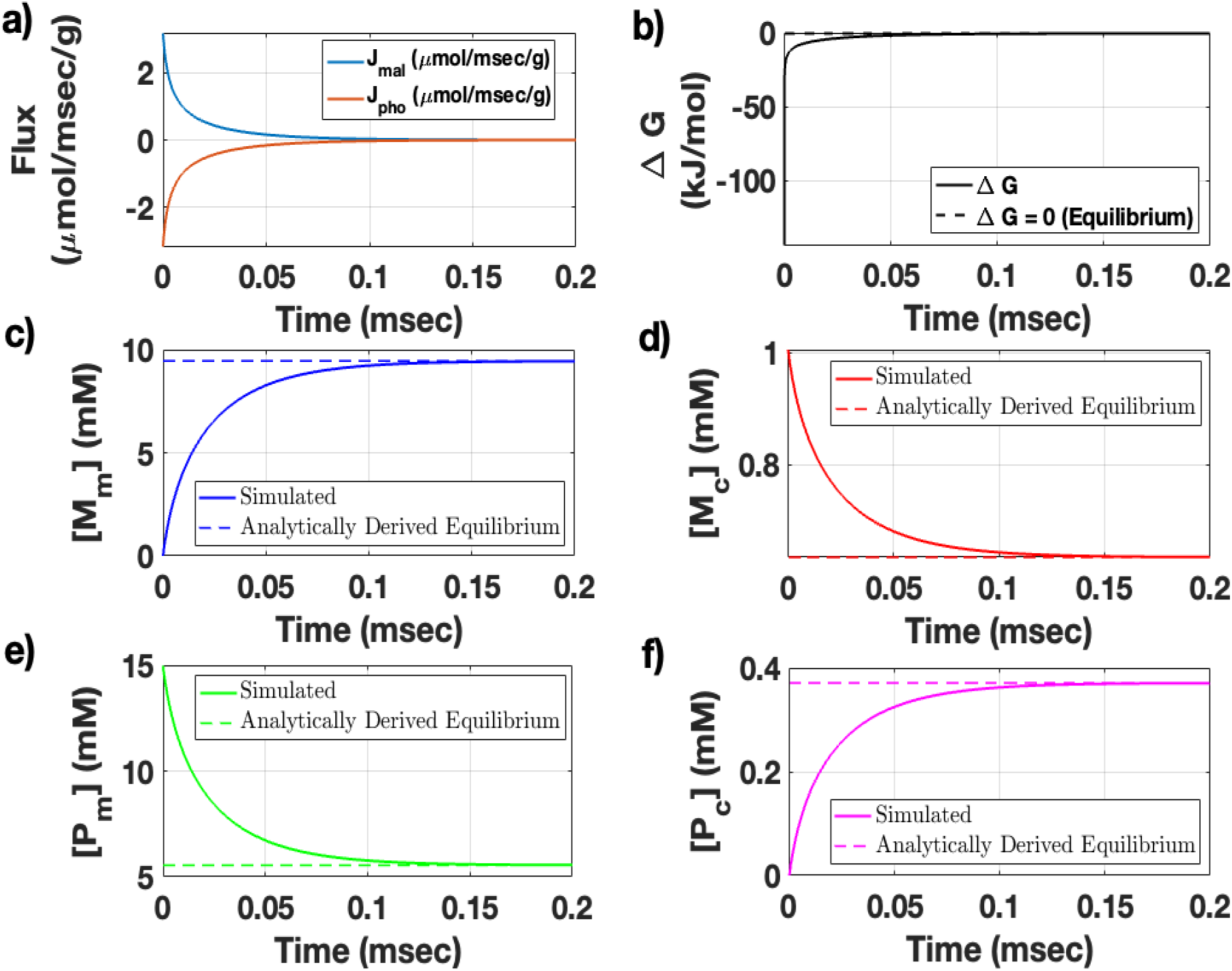
Thermodynamic analysis and equilibrium validation of the SLC25A10 antiport model in the proteoliposome setting. (a) Time-dependent fluxes of malate (*J*_mal_) and phosphate (*J*_pho_) showing rapid relaxation toward equilibrium under electroneutral exchange stoichiometry. (b) Time-resolved free-energy change (Δ*G*) for the malate/phosphate exchange reaction M_*c*_ +P_*m*_ ⇌ M_*m*_ +P_*c*_; the dashed line indicates Δ*G* = 0 (thermodynamic equilibrium). (c–f) Simulated concentrations of malate and phosphate in the internal compartment (*m*) and the external compartment (*c*; extraliposomal assay medium) compared with analytically derived equilibrium values (dashed lines). Initial conditions are [*M*_*c*_] = 1 mM and [*P*_*m*_] = 15 mM, with [*M*_*m*_] = 0 and [*P*_*c*_] = 0. Simulations use the proteoliposome compartment volumes (per gram protein) reported in Table 1. Parameter values for 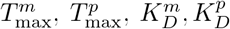, and *λ*_23_ are fixed at their posterior modes (Table 5).

An analogous validation holds in the proteoliposome setting (Fig. 11). Here, the malate and phosphate fluxes, *J*_mal_ and *J*_pho_, exhibit rapid relaxation to zero (Fig. 11a), and the corresponding Δ*G* trajectory approaches Δ*G* = 0 (Fig. 11b). As in the intactmitochondria case, the simulated equilibrium concentrations match the analytically derived equilibrium values for the malate/phosphate exchange system (Fig. 11c–f).

Together, these results demonstrate that the model implements a reversible, electroneutral 1:1 antiport constraint between one bound substrate and one counter-substrate, and that its dynamics are thermodynamically consistent: modelled fluxes decay as driving forces dissipate, and the system converges to the analytically predicted equilibrium state. In the intact-mitochondria configuration, this equilibration illustrates how a dicarboxylate–phosphate exchange mechanism can redistribute succinate (or malate) relative to the extra-matrix compartment while maintaining strict coupling to phosphate counter-transport, linking dicarboxylate handling relevant to TCA-cycle metabolism with phosphate exchange relevant to mitochondrial bioenergetics.

### 4.4 Non-equilibrium dynamics of the dicarboxylate carrier (SLC25A10)

To investigate the dynamic behaviour of the SLC25A10 transporter under experimentally realistic conditions, we simulated a competitive transport scenario analogous to the classical malate–succinate uptake experiments of Palmieri et al. [3]. The system was initialised with 0.908 mM malate, 0.5 mM succinate, and no phosphate in the external compartment, while the mitochondrial matrix concentrations and compartment volumes are set according to Table 1. These conditions mimic isolated mitochondrial uptake assays but allow us to resolve rapid transients that are experimentally inaccessible due to the sub-second timescale of equilibration and the technical difficulty of measuring compartment-resolved metabolite gradients in real time.

Figures 12–14 reveal that the simulated dynamics organize naturally into two sequential transport phases. In the early phase, the large initial matrix phosphate pool provides the dominant thermodynamic bias for dicarboxylate uptake. This is seen directly in the concentration panels of Fig. 12: matrix malate rises most strongly while external malate falls rapidly (Fig. 12a,d), indicating that malate is the dominant imported dicarboxylate under these conditions. Matrix succinate also increases and external succinate decreases (Fig. 12b,e), but with a smaller amplitude, showing that succinate participates in the same uptake program while competing less successfully than malate. At the same time, matrix phosphate collapses and external phosphate rises sharply (Fig. 12c,f), demonstrating that phosphate export is the principal exchange process through which the initial thermodynamic imbalance is relaxed. The gradient panels make this interpretation more explicit. Initially, the malate and succinate gradients (Δ*M* and Δ*S*) are positive, favoring uptake into the matrix, whereas the phosphate gradient (Δ*P* ) is strongly negative, favoring phosphate efflux (Fig. 12g–i). During Phase 1, these gradients are rapidly dissipated as transport proceeds, so the driving forces themselves visibly collapse over time rather than remaining fixed. Biologically, this is intuitive: when phosphate is initially abundant in the matrix, the prevailing phosphate gradient biases net SLC25A10 exchange toward uptake of external dicarboxylates, with malate favoured over succinate because it competes more effectively for transport.

**Figure 12:**
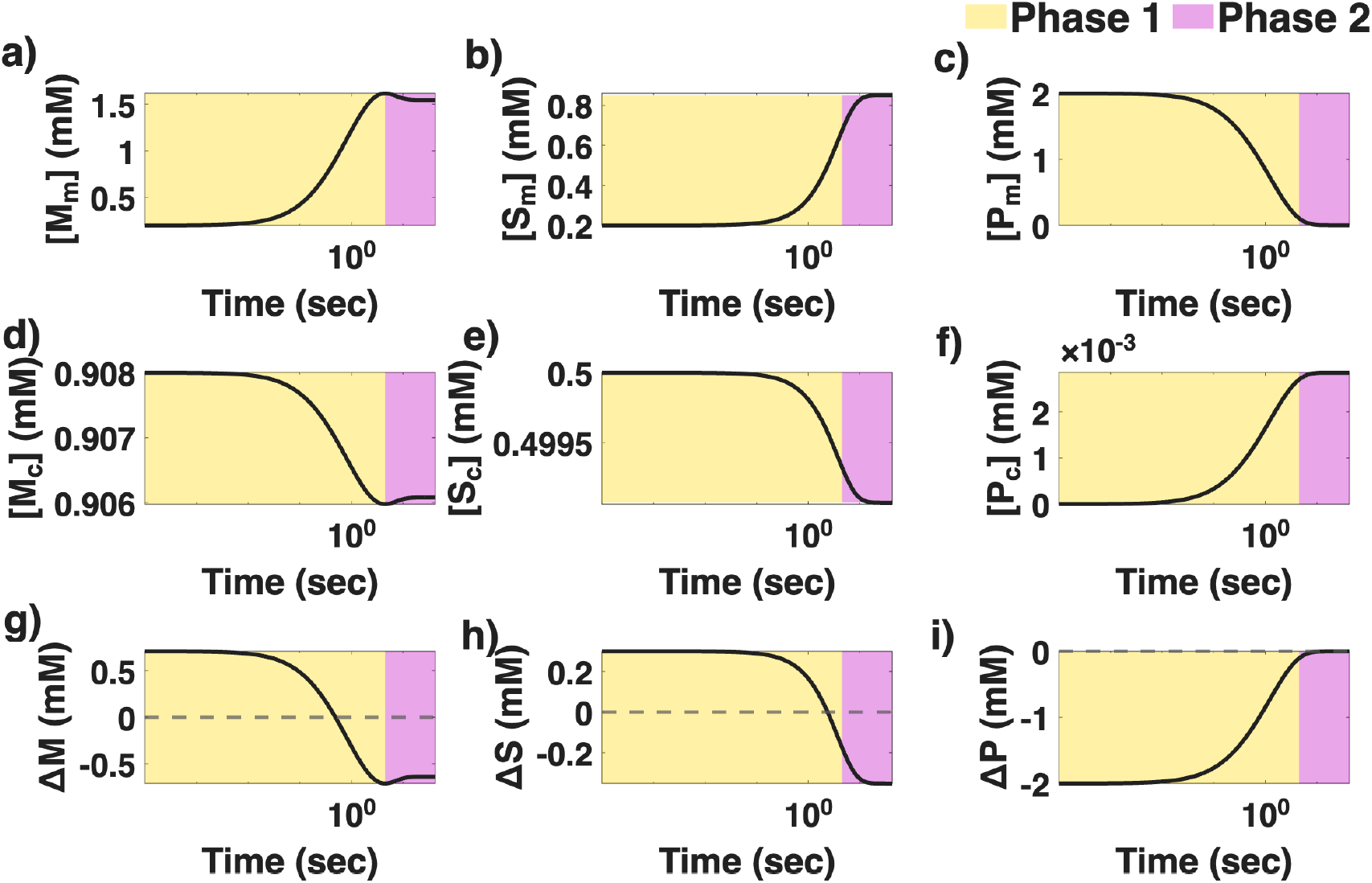
Time-dependent concentration and gradient dynamics of the SLC25A10 transporter in the intact-mitochondria setting, resolved into two transport phases. (a) Matrix malate concentration ([*M*_*m*_]); (b) matrix succinate concentration ([*S*_*m*_]); (c) matrix phosphate concentration ([*P*_*m*_]); (d) external malate concentration ([*M*_*c*_]); (e) external succinate concentration ([*S*_*c*_]); and (f) external phosphate concentration ([*P*_*c*_]). (g) malate concentration gradient (Δ*M* = [*M*_*c*_] − [*M*_*m*_]); (h) succinate concentration gradient (Δ*S* = [*S*_*c*_] − [*S*_*m*_]); and (i) phosphate concentration gradient (Δ*P* = [*P*_*c*_] − [*P*_*m*_]). The time axis is shown on a logarithmic scale. The shaded yellow region (Phase 1; 0– 4.5 sec) denotes the early phosphate-coupled heteroexchange regime, with malate/phosphate (M/P) and succinate/phosphate (S/P) exchange dominating as indicated in the respective panels. The shaded purple region (Phase 2; 4.5 sec to end) denotes the later malate/succinate (M/S) exchange regime that emerges after substantial depletion of matrix phosphate. Initial external concentrations are [*M*_*c*_] = 0.908 mM, [*S*_*c*_] = 0.50 mM, and [*P*_*c*_] = 0 mM. Initial matrix concentrations are [*M*_*m*_] = 0.20 mM, [*S*_*m*_] = 0.20 mM, and [*P*_*m*_] = 2.0 mM. The intact-mitochondria compartment volumes (per assay) are *V*_*m*_ = 1.42 *µ*L and *V*_*c*_ = 1 mL with protein-normalised volumes 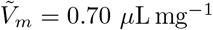 and 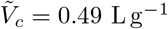. Parameter values are set to the intact-mitochondria posterior summary (Table 4).

Figure 13 clarifies the thermodynamic origin of this phase structure. In the early phase, both the malate/phosphate and succinate/phosphate exchange reactions have strongly displaced free energies, indicating that phosphate-coupled exchange is far from equilibrium and therefore strongly favoured. By contrast, the malate/succinate exchange reaction follows a distinct trajectory and changes sign across the simulation, implying that the preferred direction of malate–succinate redistribution reverses as the phosphate gradient is exhausted. This sign change is important because it shows that the later dynamics are not simply a weaker continuation of the initial regime. Instead, once matrix phosphate has been substantially depleted, phosphate-coupled transport loses its thermodynamic leverage, phosphate flux approaches zero, and the carrier enters a second, slower redistribution phase in which malate and succinate continue to exchange mainly against each other. The gradient panels in Fig. 12 are consistent with this transition: Δ*P* moves strongly toward zero as the phosphate reservoir is spent, while Δ*M* and Δ*S* continue to evolve and can partially reverse, indicating that the balance of dicarboxylate driving forces has been reorganized. In concentration space, this appears as continued readjustment of malate and succinate with only small further phosphate changes; in gradient space, it appears as depletion and partial reversal of the transmembrane concentration differences; in flux space, it appears as collapse of all flux magnitudes toward zero; and in thermodynamic space, it appears as all three exchange modes relaxing toward Δ*G* = 0.

**Figure 13:**
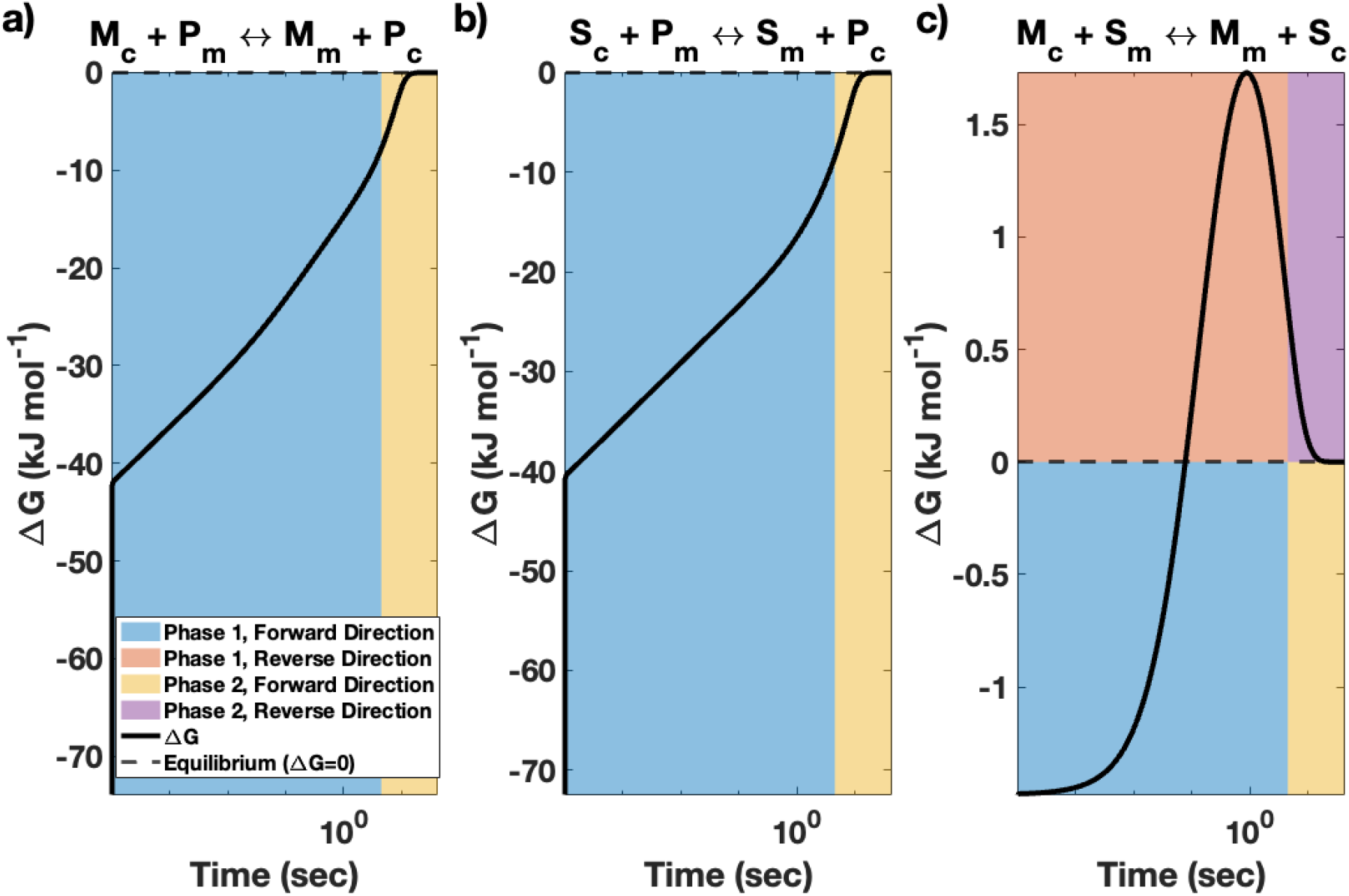
Free-energy trajectories for the three dominant SLC25A10 exchange modes in the intact-mitochondria setting. (a) Free energy for the malate/phosphate exchange reaction M_*c*_ +P_*m*_ ⇌ M_*m*_ + P_*c*_. (b) Free energy for the succinate/phosphate exchange reaction S_*c*_ + P_*m*_ ⇌ S_*m*_ + P_*c*_. (c) Free energy for the malate/succinate exchange reaction M_*c*_ +S_*m*_ ⇌ M_*m*_ +S_*c*_. The dashed horizontal line marks thermodynamic equilibrium (Δ*G* = 0). The time axis is shown on a logarithmic scale. Shaded regions indicate the thermodynamically favoured direction in each phase: blue, Phase 1 forward direction; red, Phase 1 reverse direction; yellow, Phase 2 forward direction; purple, Phase 2 reverse direction. Initial external concentrations are [*M*_*c*_] = 0.908 mM, [*S*_*c*_] = 0.50 mM, and [*P*_*c*_] = 0 mM. Initial matrix concentrations are [*M*_*m*_] = 0.20 mM, [*S*_*m*_] = 0.20 mM, and [*P*_*m*_] = 2.0 mM. The intact-mitochondria compartment volumes (per assay) are *V*_*m*_ = 1.42 *µ*L and *V*_*c*_ = 1 mL with protein-normalised volumes 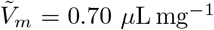 and 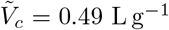. Parameter values are set to the intact-mitochondria posterior summary (Table 4).

The flux trajectories in Fig. 14 show the same regime in mechanistic terms. During the early phase, *J*_mal_ *>* 0 and *J*_suc_ *>* 0, whereas *J*_pho_ *<* 0, so the carrier is executing coordinated dicarboxylate influx coupled to phosphate efflux (Fig. 14a). The much larger magnitude of *J*_mal_ relative to *J*_suc_ explains why malate accumulates more strongly in the matrix than succinate. The total transport magnitude *J*_tot_ is also maximal at early times and then falls abruptly (Fig. 14b), showing that most carrier turnover is concentrated in an initial burst rather than being sustained uniformly over time. Biologically, this means that SLC25A10 can respond rapidly to an imposed substrate imbalance, but this response is self-limiting because the very phosphate gradient that powers the early burst is consumed by transport itself.

**Figure 14:**
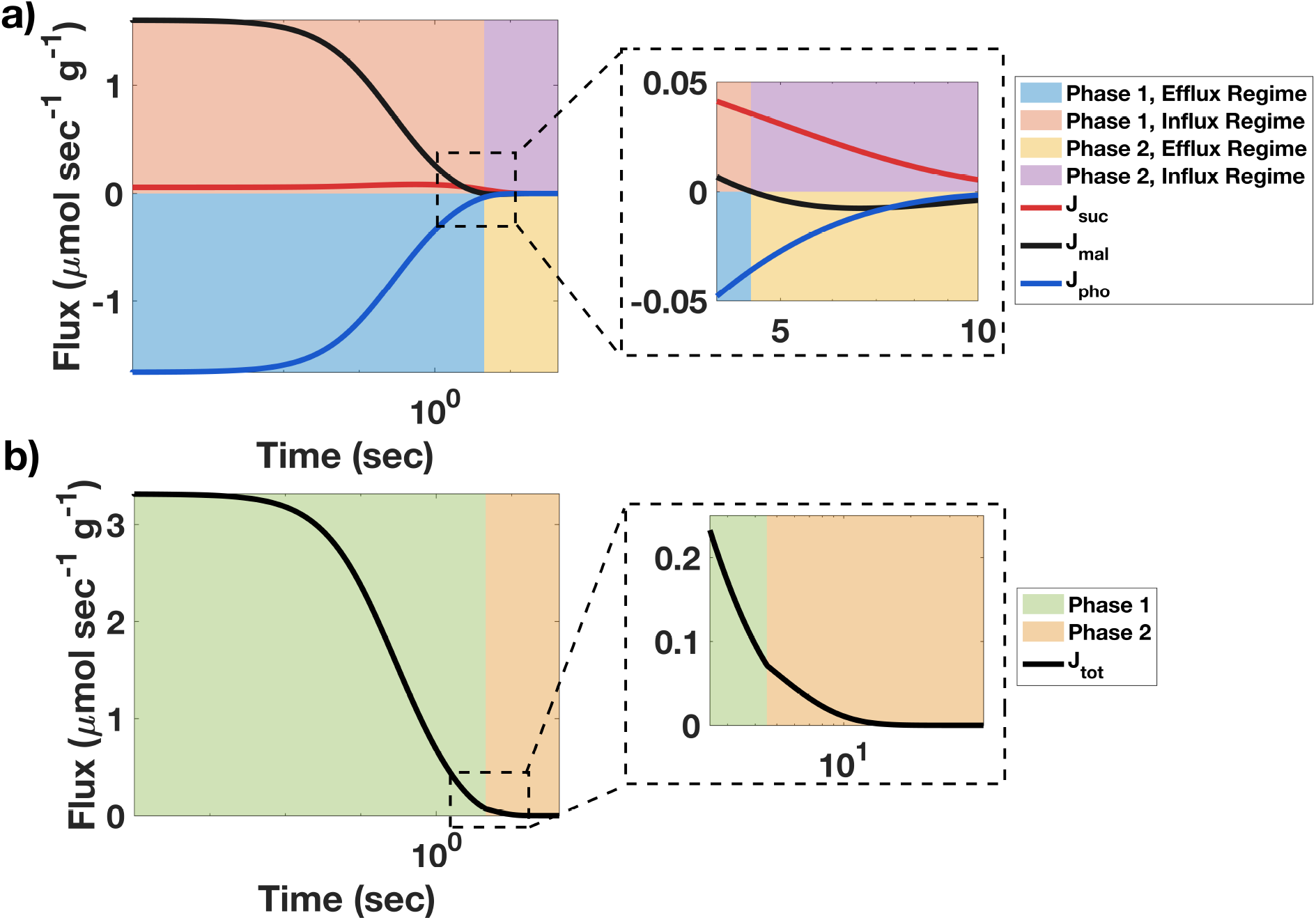
Flux and transport-magnitude dynamics of the SLC25A10 transporter in the intact-mitochondria setting. (a) Individual flux components for malate (*J*_mal_), succinate (*J*_suc_), and phosphate (*J*_pho_) as functions of time. Positive flux denotes net transport into the matrix (influx regime), whereas negative flux denotes net transport into the external compartment (efflux regime). The shaded backgrounds distinguish influx and efflux regimes across the two phases: blue, Phase 1 efflux regime; peach, Phase 1 influx regime; yellow, Phase 2 efflux regime; purple, Phase 2 influx regime. The early trajectory is dominated by strong malate influx and phosphate export, with a smaller succinate influx contribution, whereas after the phase transition the flux magnitudes decay toward zero as the system relaxes. (b) Total transport magnitude *J*_tot_ = |*J*_suc_| + |*J*_mal_| + |*J*_pho_|. The green shaded region denotes Phase 1, in which transport activity is maximal and dominated by phosphate-coupled dicarboxylate uptake, with malate and succinate influx coupled to phosphate export, while the orange shaded region denotes Phase 2, in which the overall transport magnitude drops sharply and approaches zero. The time axis is shown on a logarithmic scale in both panels. Initial external concentrations are [*M*_*c*_] = 0.908 mM, [*S*_*c*_] = 0.50 mM, and [*P*_*c*_] = 0 mM. Initial matrix concentrations are [*M*_*m*_] = 0.20 mM, [*S*_*m*_] = 0.20 mM, and [*P*_*m*_] = 2.0 mM. The intact-mitochondria compartment volumes (per assay) are *V*_*m*_ = 1.42 *µ*L and *V*_*c*_ = 1 mL with protein-normalised volumes 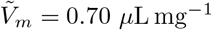 and 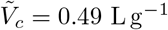. Parameter values are fixed at their posterior modes (Table 4).

Together, these three figures provide a biologically intuitive picture of SLC25A10 function under competitive conditions. The transporter first operates in a phosphate-coupled uptake regime that rapidly draws external dicarboxylates into the matrix, especially malate, and then transitions into a low-flux equilibration regime once the phosphate reservoir has been spent. The added gradient panels sharpen this interpretation because they show directly how the relevant transmembrane concentration differences are consumed by transport and thereby encode the transition between phases. This two-stage behaviour highlights that SLC25A10 is not governed by a single fixed exchange mode; rather, the dominant mode emerges dynamically from the evolving substrate pools and their associated gradients. In physiological terms, the model suggests that transient phosphate availability can strongly amplify early dicarboxylate uptake, whereas depletion of matrix phosphate suppresses further high-flux exchange and leaves the carrier to mediate a slower residual redistribution between remaining dicarboxylate gradients. Overall, these non-equilibrium dynamics illustrate how SLC25A10 can rapidly reshape matrix–external dicarboxylate and phosphate distributions following changes in substrate availability during experimental perturbations or metabolic stress [5, 4].

### 4.5 Effect of Mitochondrial Morphology on Initial SLC25A10 Fluxes

To quantify how mitochondrial morphology modulates dicarboxylate exchange, we investigated the sensitivity of the initial SLC25A10 fluxes to protein-normalised compartment volumes in the intact mitochondria setting. We considered two coupled scenarios: matrix swelling, in which 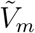 increases while 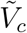 decreases, and matrix condensation, in which 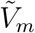 decreases while 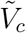 increases. The combined matrix and external (medium) specific volume per gram protein was held fixed by enforcing 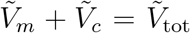, such that increases in 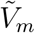 are accompanied by compensatory decreases in 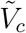, and vice versa. Here, 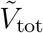 is defined as the sum of the baseline 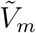 and 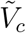 values reported in Table 1. For each condition, we recorded the succinate flux (*J*_suc_), malate flux (*J*_mal_), phosphate flux (*J*_pho_), and the total transport magnitude (*J*_tot_) at *t* = 1 s (Fig. 15).

**Figure 15:**
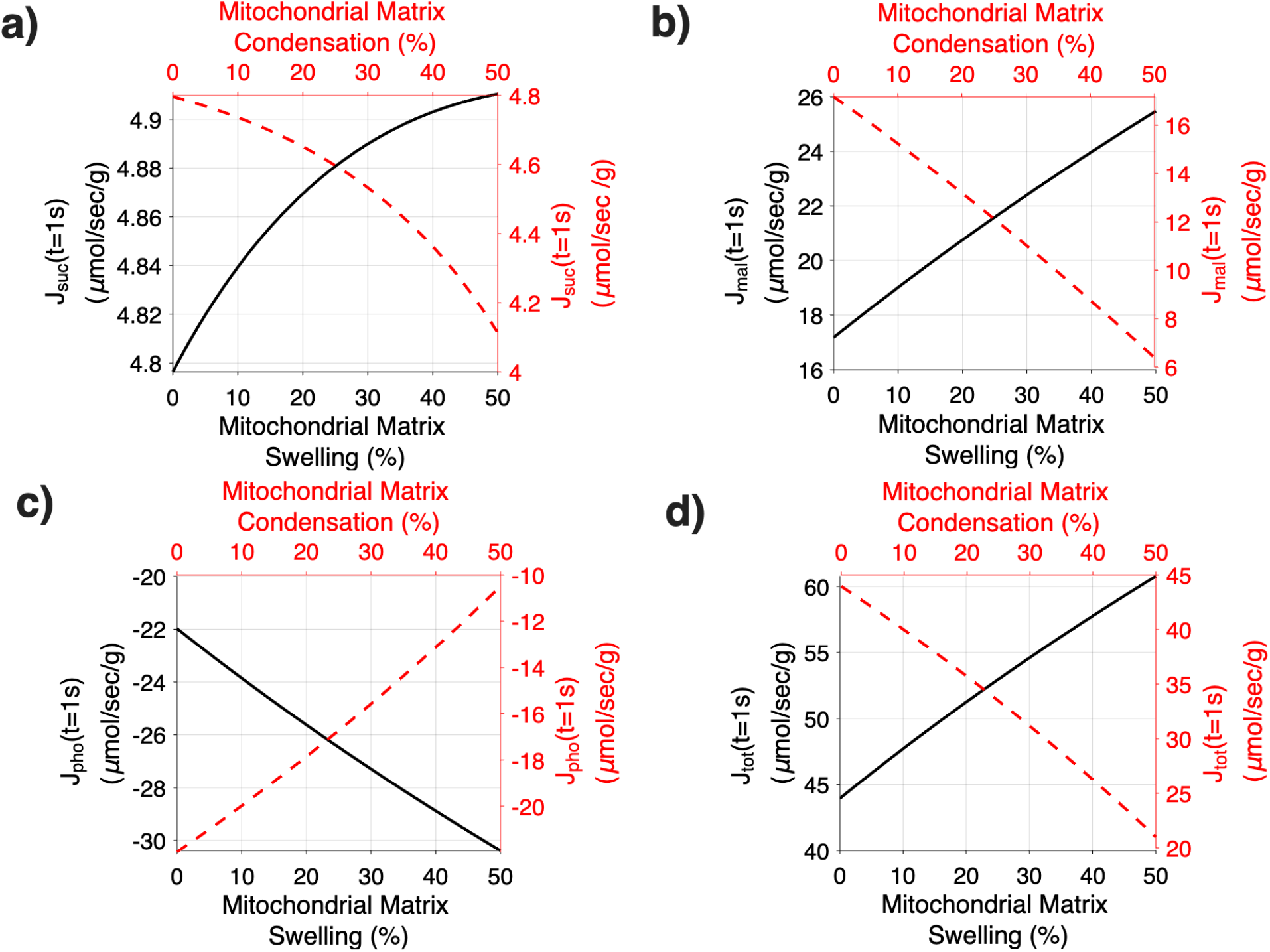
Effect of matrix swelling and condensation on initial SLC25A10 fluxes under a coupled-volume constraint in intact mitochondria. Changes in the initial fluxes of (a) succinate *J*_suc_, (b) malate *J*_mal_, (c) phosphate *J*_pho_, and (d) the total transport magnitude *J*_tot_, evaluated at *t* = 1 s, as the matrix normalised volume 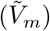 is varied between −50% (condensation; red dashed curves) and +50% (swelling; black solid curves) relative to the baseline value in Table 1. Initial concentrations are set to [*M*_*c*_] = 0.91 mM, [*S*_*c*_] = 0.5 mM, and [*P*_*c*_] = 0.1 mM on the external side, while [*M*_*m*_], [*S*_*m*_], and [*P*_*m*_] in the matrix are set according to Table 1. Parameter values for 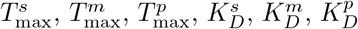, *λ*_21_, and *λ*_31_ are fixed at their posterior modes (Table 4).

Figure 15 shows the dependence of initial carrier activity on morphology under the coupled-volume constraint. Increasing the matrix volume (swelling) modestly increases both dicarboxylate flux components, *J*_suc_ and *J*_mal_ (Fig. 15a–b), while simultaneously making the phosphate flux *J*_pho_ more negative (Fig. 15c), indicating stronger phosphate export from the matrix. This coordinated change is consistent with the antiport stoichiometry: enhanced dicarboxylate influx is accompanied by an increased counter-flux of phosphate in the opposite direction. The net effect is an increase in the overall transport magnitude *J*_tot_ with swelling (Fig. 15d), indicating faster initial carrier turnover when the matrix swells and the opposing compartment contracts.

Conversely, matrix condensation produced the opposite behaviour: *J*_suc_ and *J*_mal_ decreased, *J*_pho_ became less negative (weaker counter-transport in the negative direction), and *J*_tot_ declined. Mechanistically, these trends are not driven by changes in the intrinsic binding parameters 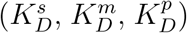, which are held fixed, but rather by how compartment volumes scale the early-time evolution of concentration gradients. In particular, swelling reduces the rate of concentration change on the matrix side while amplifying it on the external side, thereby altering the short-time gradients and the effective competition and saturation factors entering the REA flux expressions. Under the coupled-volume constraint, these effects bias the initial exchange toward larger dicarboxylate fluxes (and a correspondingly larger phosphate counter-flux) when the matrix swells.

Overall, the dependence on morphology is monotonic over the tested range: swelling systematically increases *J*_tot_, whereas condensation decreases it. Among the individual components, malate shows the largest absolute change, consistent with its larger baseline magnitude under the assay conditions. These results emphasise that morphology-induced volume redistribution—when the total matrix-plus-external volume per protein is constrained—can modulate the initial transport capacity of SLC25A10 by reshaping how rapidly gradients evolve, rather than by altering intrinsic affinity parameters. This is consistent with the general biophysical principle that compartment volumes buffer concentration changes and thereby influence transport-driven transients and effective driving forces [25, 26, 27, 28, 29]. Physiologically, moderate, transient changes in matrix volume could therefore tune early exchange rates, whereas sustained condensation could dampen initial dicarboxylate/phosphate turnover and slow the redistribution of metabolites across the inner mitochondrial membrane.

## 5 Discussion and Conclusion

In this study, we developed the first mechanistically derived and thermodynamically consistent kinetic model of the mitochondrial dicarboxylate carrier SLC25A10, based on the ping–pong mechanism. Our framework incorporates competitive binding, conformational bias weights, and thermodynamic constraints, allowing the model to quantitatively reproduce experimental transport assays while providing mechanistic insights into regimes that remain experimentally inaccessible.

The strengths of the framework extend beyond parameter estimation. Structural identifiability analysis clarified which parameters can be determined from experimental outputs and guided reparameterisation. Bayesian inference enabled statistically rigorous calibration of kinetic parameters with credible intervals, separately for intact-mitochondria uptake assays and reconstituted proteoliposome experiments, that quantify uncertainty. Thermodynamic validation ensured compliance with physical laws, confirming that fluxes, exchange reaction free energy, and equilibrium concentrations converge consistently. Together, these analyses establish a robust and generalisable modelling platform for mitochondrial carriers of the SLC25 family.

Several kinetic models of SLC25A10 have been proposed previously, including those by Bazil et al. [15] and Zhang et al. [16]. These models incorporate competitive binding, reversibility, heteroexchange, and thermodynamic consistency, but they rely on the assumption of a sequential transport mechanism. Such models are calibrated using classical transport assays [23, 20] and employ the King–Altman method [30] to derive rate equations. While conceptually valuable, this approach simplifies kinetics and lacks mechanistic depth. Recent structural and biophysical evidence, however, demonstrates that SLC25A10 is monomeric with a single binding site, consistent with a ping–pong mechanism [1]. Our work therefore advances the field by explicitly embedding this mechanism into a thermodynamically consistent kinetic framework.

A central innovation of the model is the introduction of conformational bias weights, *λ*_21_ and *λ*_31_, which quantify the relative influence of malate versus succinate and phosphate versus succinate, respectively. Together with the ratio *ϕ* (Eq. (4)), these parameters determine the directional preference of SLC25A10 by defining which substrate and which side of the membrane most strongly bias initiation of the transport cycle. Values near unity indicate equal competition, values below unity indicate succinate dominance, and values above unity indicate bias toward malate or phosphate. From a flux perspective, *J*_suc_ and *J*_mal_ depend directly on *λ*_21_, while phosphate flux is shaped by *λ*_31_. These bias weights thus provide a mechanistic lens for dissecting competitive exchange dynamics beyond what can be inferred from King–Altman–based formulations.

The inferred posterior summaries show that the kinetic parameters are constrained differently across the two assay settings. In the intact-mitochondria calibration, the maximal transport capacities for succinate 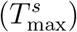, malate 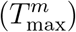, and phosphate 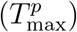, together with the dissociation constants for succinate 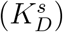, malate 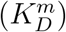, and phosphate 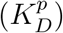, all exhibit well-behaved and comparatively concentrated posterior distributions (Figs. 6 and 4), indicating that the uptake data are informative for constraining both transport-capacity and affinity effects in the reduced model. In the proteoliposome calibration, inference is restricted to the malate–phosphate branch, and the posterior distributions of 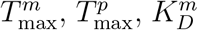, and 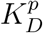 show more heterogeneous behaviour (Figs. 7 and 5). In particular, the transport-capacity parameters show clearer posterior concentration, whereas the dissociation constants, especially 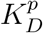, remain more strongly influenced by the prior specification, consistent with more limited information content in the reconstituted assay for separating phosphate-related effects. Biologically, these posterior patterns support robust inference of the overall transport scale in the intact-mitochondria setting, while indicating that phosphate-dependent effects remain more difficult to isolate in the proteoliposome system without additional targeted perturbations.

Another strength of our framework is its ability to provide mechanistic insight into heteroexchange behaviour that is difficult to infer from classical uptake assays [3, 23]. Conventional experiments report bulk uptake or exchange rates and do not directly resolve compartment concentrations or the microscopic carrier-state transients. In our model, flux trajectories are therefore computed as time-varying outputs of the reduced (rapid-equilibrium) transporter formulation, driven by the evolving matrix and external concentrations in the coupled mass-balance system (Fig. 12 and Fig. 14). Under these assumptions, simulations predict that the intact-mitochondria dynamics naturally separate into two phases (Figs. 12–14). In the first phase, the large initial matrix phosphate pool makes phosphate-coupled malate and succinate uptake thermodynamically favourable while phosphate is exported. The second phase emerges once that phosphate reservoir has been largely depleted. At that point, phosphate flux approaches zero, the phosphate gradient collapses, and transport shifts into a slower redistribution regime in which malate and succinate continue to readjust primarily against each other. Together, these results support a biologically intuitive picture in which matrix phosphate provides a transient thermodynamic bias for early dicarboxylate uptake, while also showing that a three-substrate ping–pong carrier need not operate in a single fixed mode over an entire trajectory. Rather, the dominant exchange pathway can switch dynamically as the transmembrane gradients are reshaped by the transport process itself.

The morphology analysis extends this logic by showing that SLC25A10 activity depends not only on intrinsic kinetic parameters but also on how compartment geometry modulates the evolution of gradients. Under the intact-mitochondria coupled-volume constraint, matrix swelling modestly increases the initial dicarboxylate influxes and increases the total transport magnitude *J*_tot_, whereas matrix condensation has the opposite effect (Fig. 15). Mechanistically, these trends arise from how coupled volume redistribution reshapes the early-time evolution of concentration gradients and saturation terms in the reduced flux expressions, rather than from changes in intrinsic binding parameters (which are held fixed). These results are consistent with the broader view that crista geometry and compartment volumes influence carrier-driven transients and effective driving forces [26, 29], linking mitochondrial structure to short-timescale metabolic exchange capacity.

These results have important implications for pathological settings involving excessive succinate accumulation in the mitochondrial matrix, which has been associated with multiple disease states, including cancer and metabolic disorders [5]. In mitochondria, succinate levels are regulated by enzymes such as succinyl-CoA synthetase (SCAS), which generates succinate, and succinate dehydrogenase (SDH), which consumes it [14, 15, 16, 17]. Inactivating mutations in SDH impair succinate oxidation and can lead to pathological succinate accumulation [14, 12, 13, 5, 54]. In this biological context, an important mechanistic question is how SLC25A10-mediated exchange may respond when the usual metabolic sink for succinate is compromised.

Figure 12 shows that succinate transport is highly sensitive to phosphate-coupled exchange during the initial high-flux regime, while the concentration-gradient dynamics (Fig. 12g–i) and the corresponding free-energy analyses (Fig. 13) and flux (Fig. 14) indicate that this regime is transient because the relevant driving forces are rapidly dissipated. This suggests that, in metabolic settings where matrix succinate accumulates, SLC25A10-mediated succinate redistribution will depend not only on succinate abundance itself, but also on the availability of counter-substrates, especially phosphate, and on competition from other dicarboxylates such as malate. Thus, succinate accumulation alone is unlikely to determine transport behaviour; rather, the dominant exchange mode is expected to depend on the broader metabolic state that sets the transmembrane substrate gradients.

These observations motivate a natural next step beyond the present reduced carrier model. In particular, the reduced ping–pong transport-rate equations for SLC25A10 developed here could be embedded in a larger mitochondrial network model that includes additional SLC25-family carriers, the phosphate carrier, and central metabolic pathways such as the TCA cycle and the electron transport chain [15, 16, 17]. Within such a framework, one could investigate how the surrounding transport and metabolic network determines the effective exchange mode of SLC25A10 under conditions of pathological succinate accumulation. We emphasise, however, that the present model does not explicitly include these additional transporters and pathways, and therefore such questions remain beyond the scope of the current study.

A second and complementary future direction is to study the effect of SLC25A10 perturbation on broader cellular metabolite dynamics. Our targeted metabolomic measurements reported in Appendix D show that perturbation of SLC25A10 is associated with changes in succinate and related dicarboxylates at the whole-cell level. While these data do not resolve mitochondrial transport directionality and therefore cannot be used as mechanistic validation of the transporter model, they do indicate that SLC25A10 perturbation has measurable downstream consequences for cellular metabolic state. This is particularly relevant for succinate, which is an established oncometabolite [5]. A broader cellular transport–metabolism model that couples mitochondrial transport to cytosolic and whole-cell metabolic dynamics would therefore provide a more appropriate framework for investigating how SLC25A10 perturbation influences metabolite accumulation, redistribution, and metabolic rewiring at the cellular level [14].

Despite these advances, the model has several limitations. Our Bayesian calibration adopts simplifying assumptions about both the likelihood and the priors. For the likelihood, we assume independent, identically distributed Gaussian errors on the flux measurements. In practice, however, experimental errors and model–data discrepancy may be heteroskedastic (for example, larger fluxes may exhibit larger variance), correlated across conditions within an assay, or non-Gaussian owing to detection limits or instrumental drift. Such effects could influence posterior uncertainty and, in principle, bias parameter estimates. Nevertheless, we adopt this Gaussian error model here to enable tractable likelihood evaluation and parameter inference, and we view the present work as a foundation for future calibration methods that incorporate more realistic error structures.

The prior specification is also simplified and assay-dependent. For the intact-mitochondria calibration, we use uniform priors for the inferred kinetic parameters, reflecting the absence of strong assay-specific prior constraints. For the proteoliposome calibration, where inference is restricted to the malate–phosphate branch, we use informative Gaussian priors for the corresponding transport-capacity and dissociation parameters based on literature values. While these choices are pragmatic and computationally convenient, they may influence posterior concentration in settings where the data are only weakly informative. This issue is especially important for parameters that also exert strong influence on model outputs, as indicated by the sensitivity analysis.

A related limitation is that the impact of prior specification is not uniform across parameters. The sensitivity analysis shows that several transport-capacity and phosphate-related parameters exert strong influence on the predicted fluxes, whereas the posterior analysis indicates that some of these same parameters remain only weakly constrained by the available data, particularly in the proteoliposome calibration. As a result, prior assumptions may have a disproportionate effect on the uncertainty and interpretation of the most influential model components. This motivates future work combining more informative experiments with formal prior-sensitivity analysis to determine more clearly which conclusions are data-driven and which remain sensitive to prior specification.

Additionally, the distribution of dicarboxylates and phosphate among their protonation (and, where relevant, metal-bound) species depends on pH and ionic conditions, which can influence the concentrations of the transporter-relevant charged forms (e.g. HPO_4_^2−^). In the present work, we model transport under the experimental assay conditions, where pH and buffer composition are controlled and effectively constant; accordingly, speciation is treated as fixed and its effect is absorbed into the inferred effective kinetic parameters (e.g., *K*_*D*_ parameters and *T*_max_). Incorporating explicit pH-dependent speciation (and cation binding) would be most informative when datasets spanning multiple pH values and ionic conditions are available, enabling those additional degrees of freedom to be constrained rather than introducing non-identifiable parameters.

Moreover, several modelling assumptions limit the scope of the present framework. First, while the model is mechanistically grounded in a ping–pong scheme, we employ a reduced rapid-equilibrium treatment of the carrier complex (fast binding/unbinding), which collapses intermediate carrier-state transients and may miss finer regulatory steps or additional conformational intermediates. Relatedly, parameter lumping in the reduced flux expressions can mask asymmetries between binding and release kinetics that would be resolvable only with richer time-resolved datasets.

Second, compartments are treated as well-mixed volumes, neglecting sub-mitochondrial microdomains, diffusion limitations, and interactions with other carriers that could be important in highly structured mitochondria. Finally, calibration relies on isolated uptake assays (intact mitochondria and proteoliposomes) under controlled buffer conditions; tissue-specific regulation, post-translational modifications, and in vivo substrate gradients are not represented, which limits direct translatability to physiological and pathological settings. Future work should integrate additional carrier states, broader metabolic coupling, and spatially resolved compartmentalisation within whole-mitochondrion models, supported by more diverse experimental measurements.

In conclusion, we present a mechanistically derived and thermodynamically consistent kinetic model of the mitochondrial dicarboxylate carrier SLC25A10 grounded in a ping– pong, single-binding-site mechanism. Using a reduced rapid-equilibrium flux formulation with Bayesian calibration to both intact-mitochondria uptake assays and proteoliposome data, the model quantitatively reproduces the measured exchange kinetics. The calibrated framework captures competitive coupling among succinate, malate, and phosphate, reveals two-phase non-equilibrium transport dynamics under intact-mitochondria assay conditions, and shows that morphology-induced volume redistribution can systematically modulate early exchange capacity by reshaping the evolution of substrate gradients.

Beyond reproducing classical transport assays, the framework provides a mechanistically interpretable and transferable basis for studying how substrate competition, thermodynamic driving forces, and compartment geometry interact to shape mitochondrial carrier behaviour. More broadly, our approach—combining mechanistic reduction, identifiability analysis, Bayesian inference, and thermodynamic validation—offers a general workflow that can be adapted to other SLC25 carriers to connect structure-informed transport mechanisms with quantitative, experimentally grounded mitochondrial transport dynamics.

## Data Availability

The core MATLAB codes supporting this study are publicly available on GitHub: https://github.com/rxn315/MATHEMATICAL_MODEL_SLC25A10_REVISED.git. Due to file-size limitations, the full reproducibility package (including scripts used to generate all figures and tables in the revised manuscript) is archived on Zenodo: (DOI: https://doi.org/10.5281/zenodo.18664645).

## Acknowledgments

ES was supported by a Cancer Research UK grant (C42109/A24757). YL was supported by The Paradifference Foundation. FS was supported by a UKRI Future Leaders Fellowship (MR/T043571/1). We acknowledge that we used ChatGPT to refine and enhance the clarity of this article. However, all ideas and content are our own, and we take full responsibility for the final submission. Additionally, I acknowledge the use of BioRender in creating some of the scientific illustrations for this article.

## Author Contributions

**Ramin Nashebi:** Mathematical model conceptualisation, computational methodology, software, visualisation, model validation, model analysis, writing, and editing.

**Yingying Lyu:** Experimental data procurement and writing the description of the experimental work.

**Elías Vera-Sigüenza:** Model validation, model analysis, article review.

**Daniel A. Tennant:** Conceptualisation, experimental data validation, article review, and editing.

**Fabian Spill:** Mathematical model conceptualisation, model validation, article review and editing, supervised and managed the study.

## Appendix A Full ODE System for SLC25A10 Ping–Pong Model

The complete set of ordinary differential equations (ODEs) governing transporter states, complexes, and metabolite concentrations is:

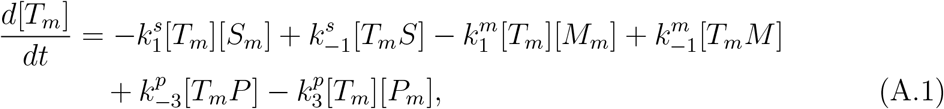

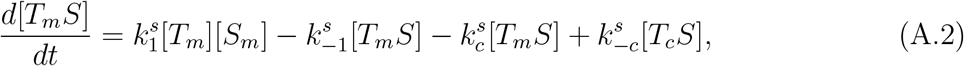

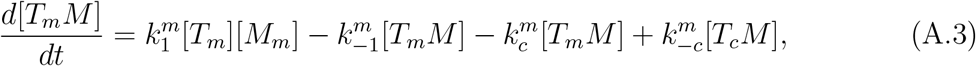

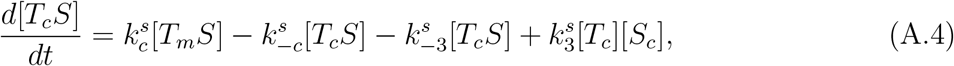

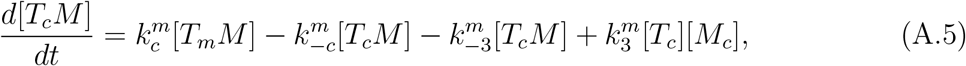

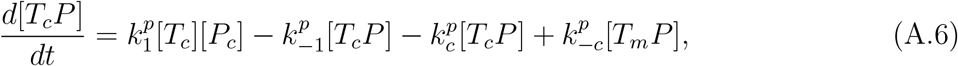

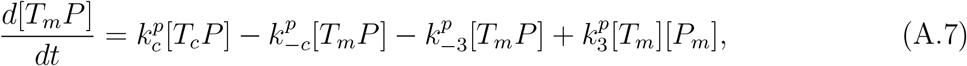

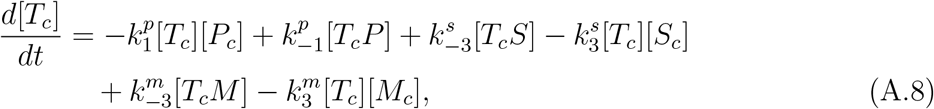

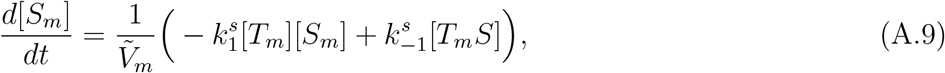

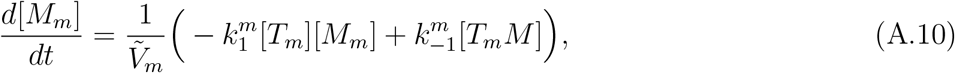

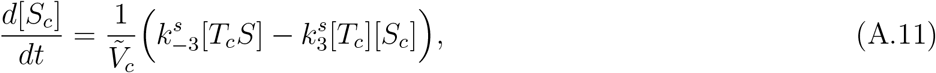

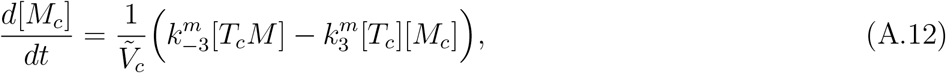

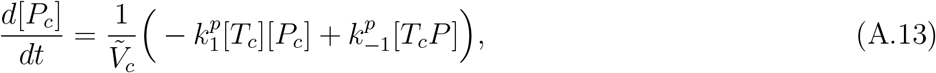

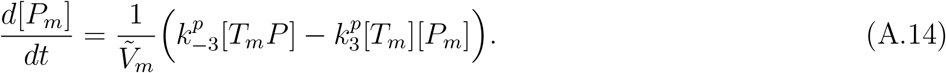

## Appendix B Symmetric Transport Assumption

We assume that the SLC25A10 transporter operates with symmetric kinetics [31, 32, 33]. Under this assumption, the elementary rate constants governing substrate binding/unbinding, conformational transitions, and substrate release have the same magnitudes when the carrier operates toward the mitochondrial matrix (*m*) or toward the external compartment (*c*).

For example, for succinate, the matrix-side binding and release rates 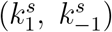 are assumed equal to the corresponding rates on the external side 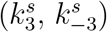:

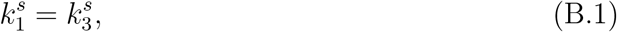

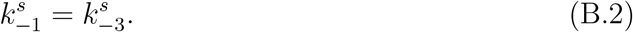

Analogous relations hold for malate and phosphate, and for the associated conformational transition steps. This symmetry reduces the number of independent kinetic parameters, facilitates parameter estimation, and ensures that the transporter satisfies microscopic reversibility and detailed balance at equilibrium.

## Appendix C Rapid–Equilibrium Assumption and Derivation of the Rate Equation

### Appendix C.1 Forward Cycle

To derive the rate equations, we invoke the rapid–equilibrium assumption (REA) for substrate binding. In the context of carrier-mediated transport, REA assumes that substrate binding and release to/from the carrier are much faster than the conformational transitions of the loaded carrier, so that the binding steps can be treated as being at thermodynamic equilibrium [37]. The slow, rate-limiting steps are the translocation (ping–pong) transitions of the loaded carrier.

For succinate on the matrix side, the binding reaction

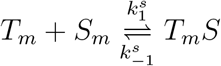

is assumed to be at equilibrium on the timescale of transport. Thus, the net rate of this binding step is zero:

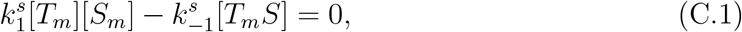

which yields

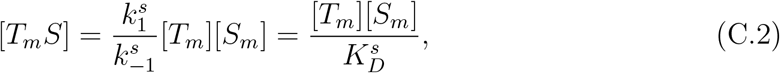

where

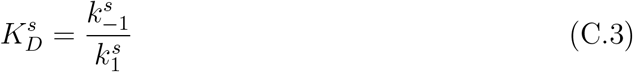

is the equilibrium dissociation constant for succinate on the matrix side. By symmetry of the carrier, the same dissociation constant applies on the external side,

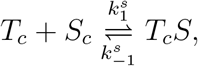

so that

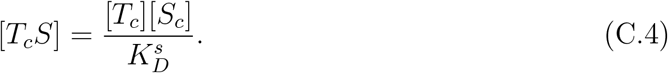

The same reasoning applies to malate and phosphate. For malate,

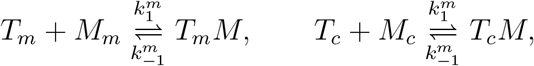

we obtain

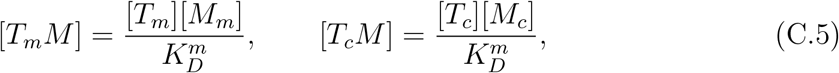

with

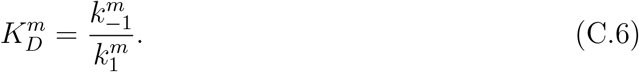

For phosphate,

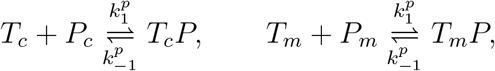

we obtain

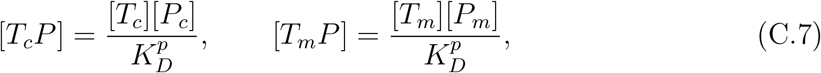

with

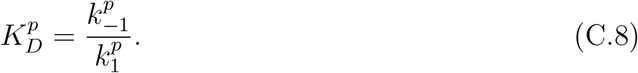

Under REA, the concentrations of the transporter–substrate complexes are therefore algebraic functions of the free transporter and substrate concentrations, as in (C.2), (C.5), and (C.7). The slow, rate–limiting steps in the forward cycle are the conformational transitions of the loaded carrier:

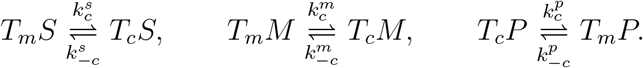

#### Appendix C.1.1 Intermediate product formation: 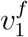

Under REA, the reversible binding and unbinding of substrates to the carrier on either side of the membrane occur much faster than the conformational (ping–pong) transition of the loaded carrier. Thus, the conformational change is the rate-limiting step, and the dicarboxylate flux is governed by the net conversion of the matrix-facing complexes (*T*_*m*_*S, T*_*m*_*M* ) into their external-facing counterparts (*T*_*c*_*S, T*_*c*_*M* ).

Accordingly, the forward-cycle flux through the dicarboxylate leg can be written as the rate of formation of the external-facing loaded transporter states:

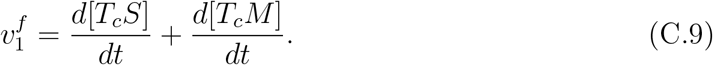

Substituting the conformational transition rates for these processes yields

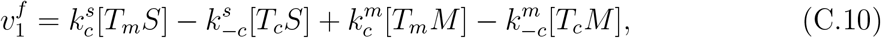

which represents the net forward flux through the rate-limiting conformational transitions of the loaded carrier.

Substituting the rapid–equilibrium expressions (C.2) and (C.5), we obtain

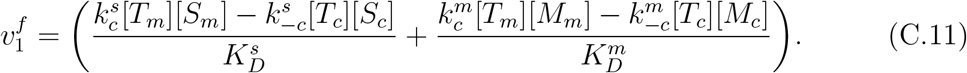

Under the symmetry assumption for the conformational transitions, this expression can be written in the compact form

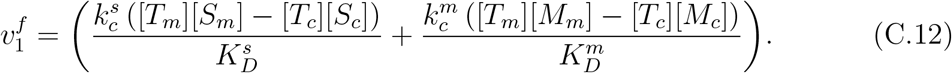

We define

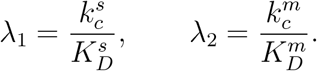

Factoring out [*T*_*m*_] and [*T*_*c*_] gives

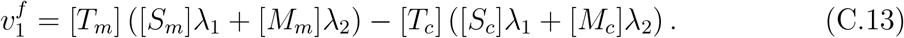

#### Appendix C.1.2 Final product formation: 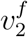

In the forward direction of the transport cycle, phosphate is released into the matrix as a consequence of the conformational (ping–pong) transition of the phosphate-loaded carrier. Under REA, phosphate binding and unbinding occur rapidly relative to this conformational change, which therefore constitutes the rate limiting step for phosphate transport in the forward direction.

Accordingly, the forward phosphate flux 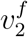 can be written as the rate at which the matrix-facing phosphate-loaded transporter (*T*_*m*_*P* ) is formed:

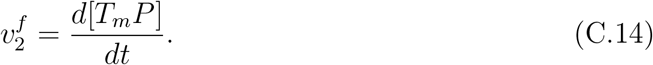

Substituting the microscopic conformational transition rates gives

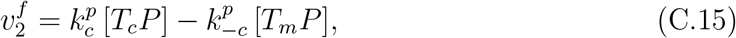

which represents the net forward flux through the rate-limiting conformational transition of the phosphate-loaded carrier. Using the rapid–equilibrium expressions (C.7),

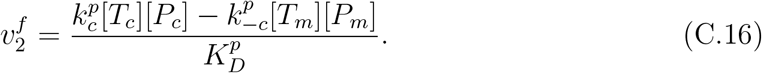

Under symmetry of the conformational rates,

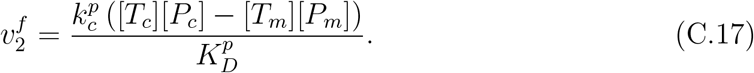

Defining

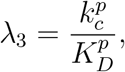

we can write

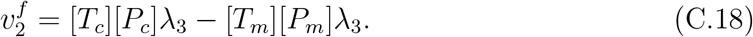

#### Appendix C.1.3 Cycle flux balance and relation between *T*_*m*_ **and** *T*_*c*_

In a cyclic ping–pong mechanism, the net flux through the cycle must be constant at steady state [55]. Thus, in the forward cycle, the rate of intermediate production 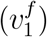 must equal the rate of final product formation 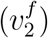:

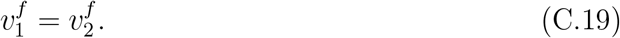

This condition also enforces the 1:1 antiport stoichiometry: each complete transport cycle exchanges exactly one bound substrate in one direction for one counter-substrate in the opposite direction at steady state.

Equating (C.13) and (C.17), we obtain

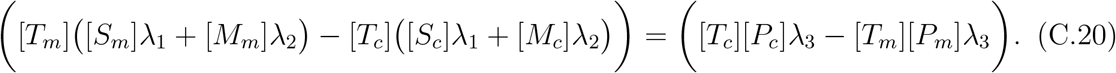

Rearranging terms,

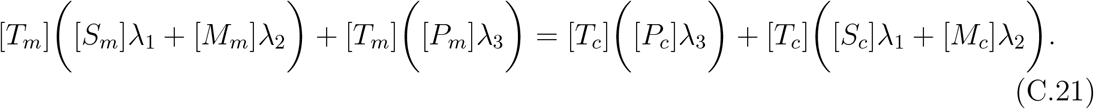

We can rewrite this as

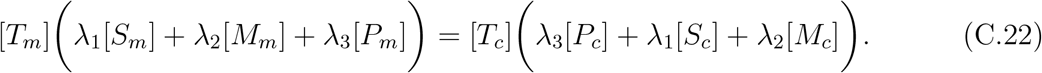

Solving for [*T*_*c*_] gives

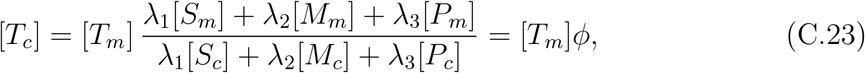

where we have defined

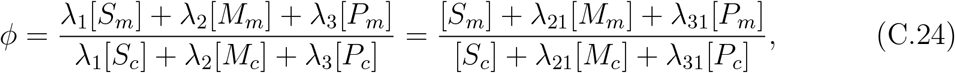

with *λ*_21_ = *λ*_2_*/λ*_1_ and *λ*_31_ = *λ*_3_*/λ*_1_.

### Appendix C.2 Reverse Cycle

In the reverse direction, the net substrate fluxes reverse relative to the chosen forward orientation. Under the three-substrate formulation, the dominant counter-substrate is not fixed a priori; depending on substrate availability, concentration gradients, and kinetic parameters, the reverse flux can involve phosphate, malate, or succinate. The same rapid-equilibrium relations for binding (Eqs. (C.2)–(C.7)) apply; only the direction of the net flux is reversed.

#### Appendix C.2.1 Intermediate product formation: 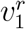

In the reverse direction of the transport cycle, dicarboxylates are delivered into the mitochondrial matrix through the conformational (ping–pong) transition of the substrate-loaded carrier. Under the rapid–equilibrium assumption, binding and unbinding of dicarboxylates occur on a much faster timescale than the conformational change; therefore, the reverse conformational transition constitutes the *rate-limiting step* for dicarboxylate transport into the matrix.

The reverse flux 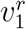 can be expressed as the rate of formation of the matrix-facing transporter–substrate complexes:

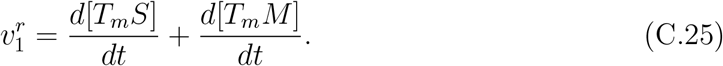

Substituting the microscopic conformational transition rates yields

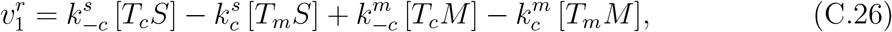

which represents the net reverse flux of succinate and malate delivered to the matrix via the rate limiting conformational transitions of the substrate loaded carrier. Using the rapid equilibrium relations

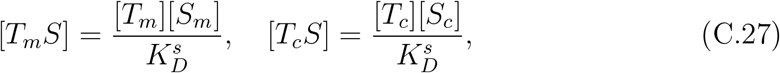

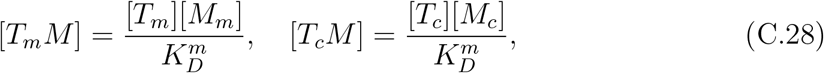

we obtain

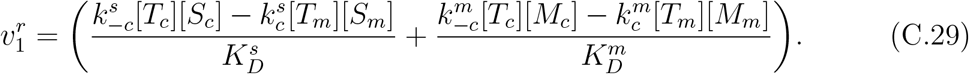

Under the symmetry assumption for the conformational transitions we write

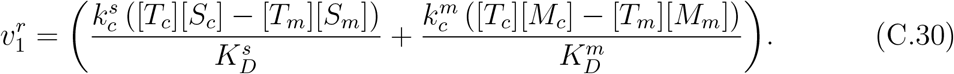

Introducing the effective transport efficiencies

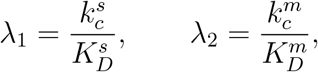

we can express 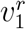 compactly as

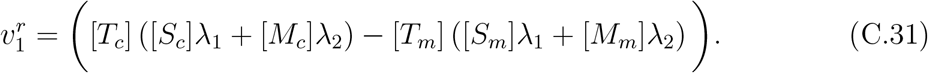

#### Appendix C.2.2 Final product formation: 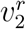

In the reverse direction of the transport cycle, phosphate is released into the external compartment as a consequence of the conformational (ping–pong) transition of the phosphate-loaded carrier. As in the forward cycle, phosphate binding and dissociation occur much faster than the conformational change; thus, the reversible conformational transition represents the rate limiting step governing phosphate export to the external side.

Accordingly, the reverse phosphate flux 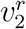 can be written as the rate of formation of the external-facing phosphate-loaded transporter:

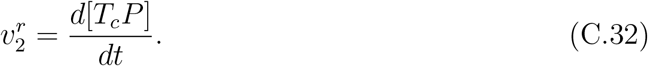

Substituting the microscopic conformational transition rates gives

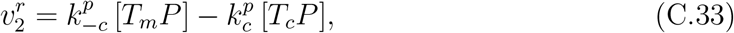

which represents the net reverse flux of phosphate delivered to the external compartment via the rate limiting conformational transition of the phosphate-loaded carrier. Using the rapid equilibrium expressions

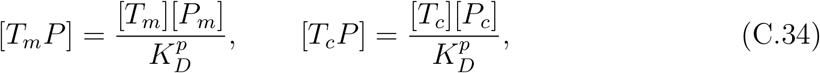

we obtain

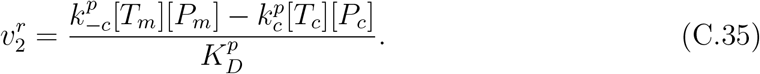

Under symmetry of the conformational rates this becomes

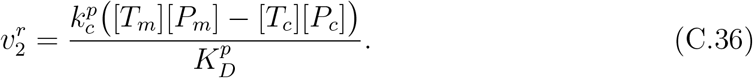

Defining

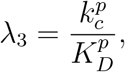

we can write

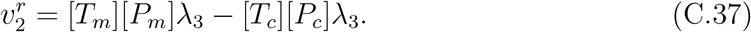

#### Appendix C.2.3 Reverse-cycle flux balance and relation between *T*_*m*_ **and** *T*_*c*_

As in the forward cycle, the net flux through the cycle must be constant at steady state [55]. In the reverse direction, this implies that the substrate flux through one half-cycle equals the corresponding counter-substrate flux through the return half-cycle, preserving the closed 1:1 antiport cycle:

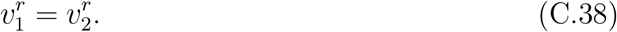

Equating (C.31) and (C.37) gives

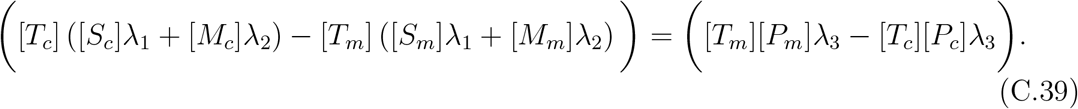

Rearranging terms,

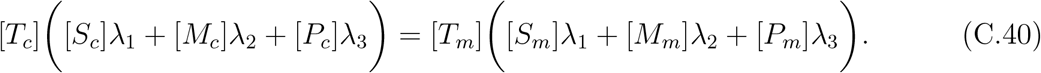

Solving for [*T*_*c*_] yields

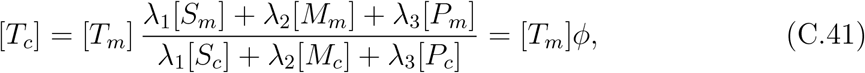

where we define

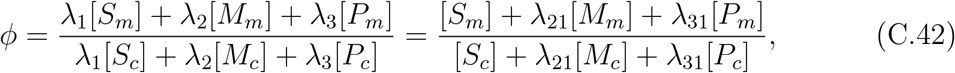

with *λ*_21_ = *λ*_2_*/λ*_1_ and *λ*_31_ = *λ*_3_*/λ*_1_.

### Appendix C.3 Total transporter conservation

The total transporter amount per gram protein [*T*_total_] is distributed among all transporter states. We write

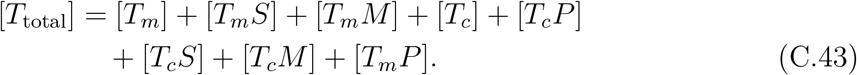

Using the rapid equilibrium relations (C.2), (C.5), and (C.7), this becomes

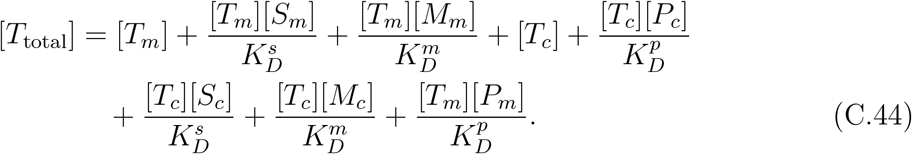

Equivalently,

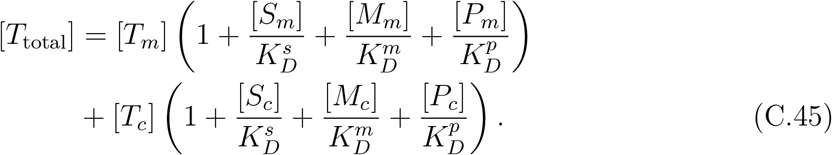

We define

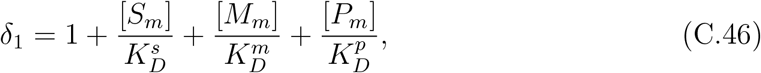

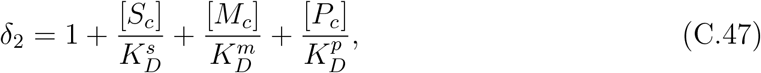

so that the total transporter conservation can be written as

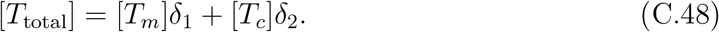

Substituting (C.23) into (C.48) yields

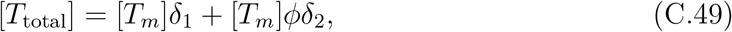

so that

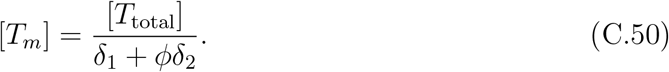

#### Appendix C.3.1 Final rate expressions for forward cycle

Substituting (C.50) into the forward flux expression (C.13) yields

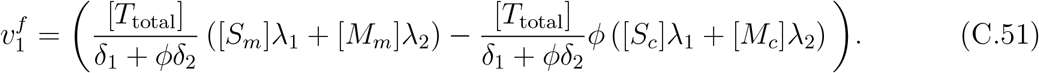

Equivalently,

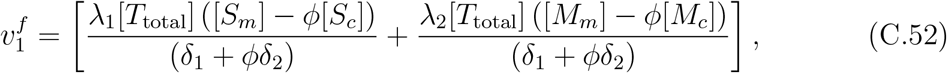

Recalling that 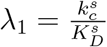 and 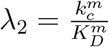, and defining 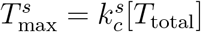 and 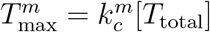, we obtain

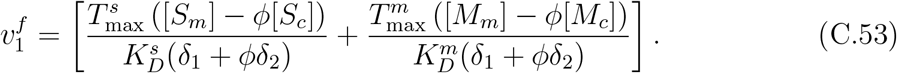

Similarly, substituting (C.50) into the phosphate flux expression (C.17) yields

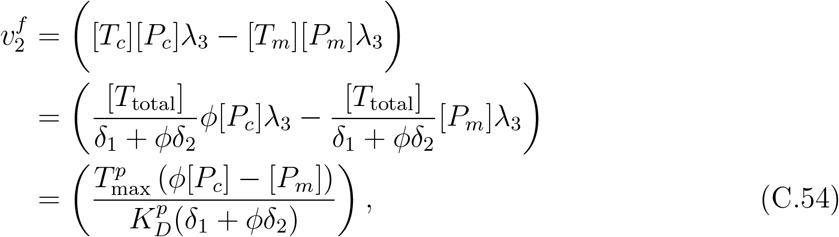

where 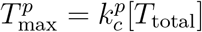.

#### Appendix C.3.2 Final rate expressions for reverse cycle

Substituting (C.50) into the reverse flux expression (C.31) yields

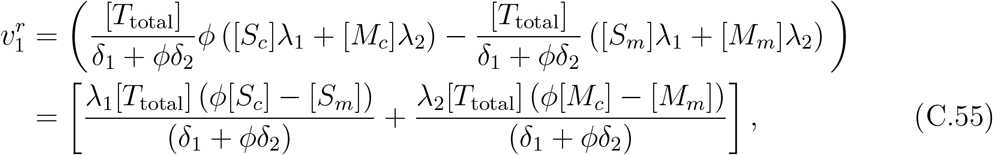

Recalling that 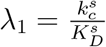 and 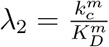, and defining 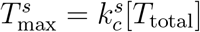 and 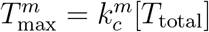, we obtain

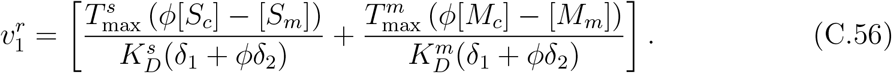

Similarly, substituting [*T*_*m*_] and [*T*_*c*_] = [*T*_*m*_]*ϕ* into (C.37), we obtain

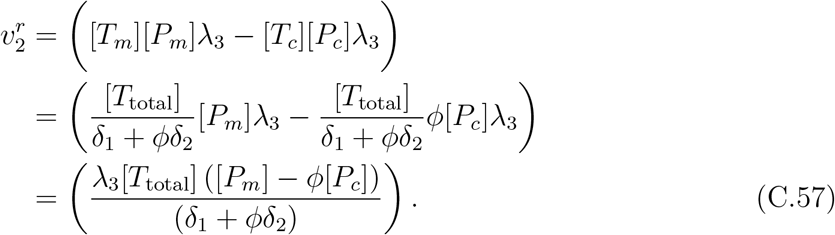

Defining 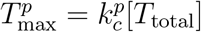, we obtain

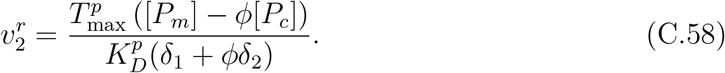

### Appendix C.4 Net Flux of the Complete Cycle

The complete transport cycle is reversible and can proceed in both the forward and reverse directions. Under REA, the forward-cycle fluxes of dicarboxylates and phosphate are

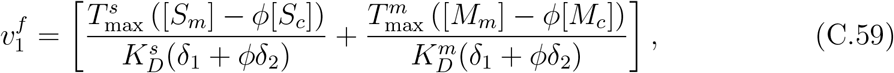

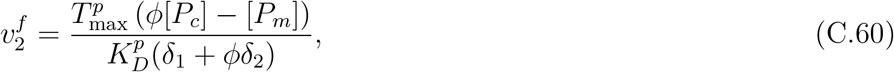

where

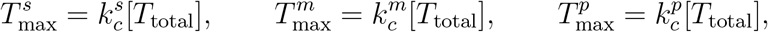

and

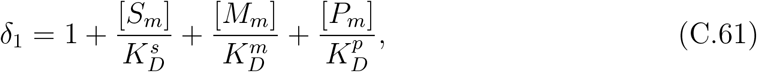

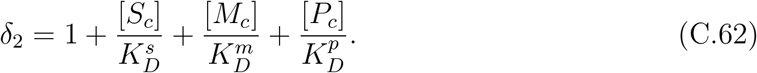

The corresponding reverse-cycle fluxes are

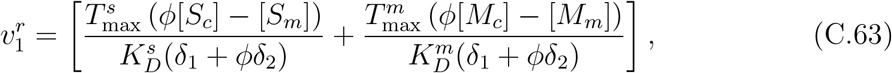

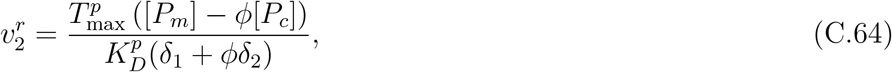

with

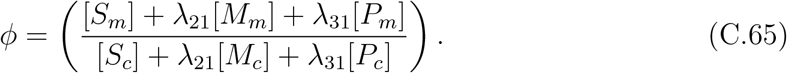

Recalling that 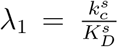 and 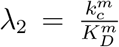, and defining 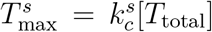 and 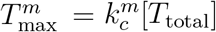, we can write that:

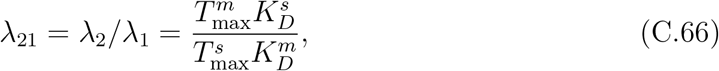

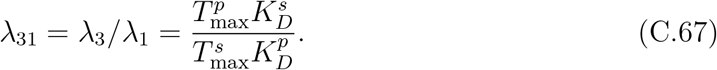

#### Appendix C.4.1 Net dicarboxylate flux

We define the net dicarboxylate exchange rate as the difference between the reverse and forward contributions,

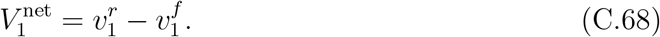

Substituting the expressions above for 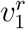 and 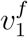 yields

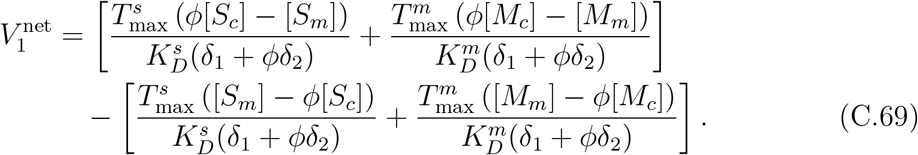

We decompose this into separate contributions from succinate and malate,

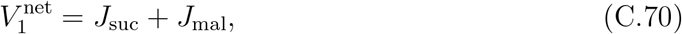

where

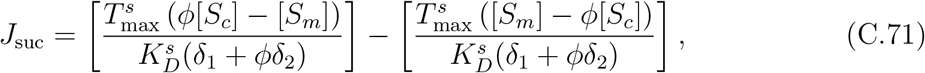

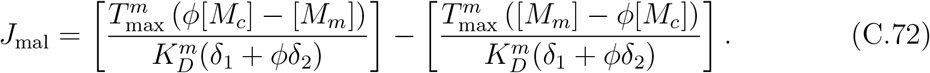

#### Appendix C.4.2 Net phosphate flux

Analogously, the net phosphate exchange rate is defined as

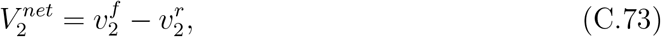

so that

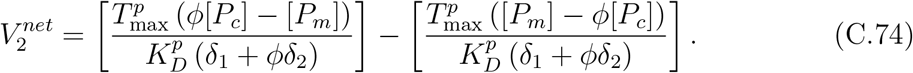

We denote the corresponding net phosphate flux by

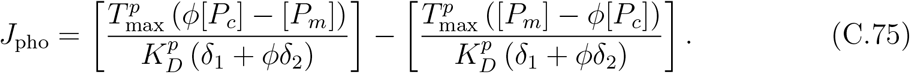

These expressions provide the net substrate fluxes for the complete reversible three-substrate ping–pong cycle under the rapid-equilibrium assumption, with competitive binding of succinate, malate, and phosphate at a single site. The dicarboxylate/phosphate grouping used in the algebraic derivation corresponds to the calibration orientation, whereas the resulting reduced flux expressions can support S/P, M/P, and M/S exchange regimes depending on substrate gradients and kinetic parameters.

## Appendix D Experimental Protocol and Supplementary Targeted Metabolomics Measurements in SLC25A10-Perturbed Chromaffin Cells

To elucidate the metabolic consequences of diminished SLC25A10 activity, we conducted a comparative metabolomic analysis. We used immortalized murine chromaffin cells (imCCs) wherein an inducible knockdown of Slc25a10 (SLC25A10-KD) was established, alongside corresponding scramble-control imCCs serving as controls. Subsequent to verification of the efficacy of SLC25A10 knockdown, extracts of cellular metabolites of both SLC25A10-KD and scramble-control imCCs were subjected to comprehensive metabolomic profiling based on liquid chromatography-mass spectrometry (LC-MS).

The metabolomic data revealed significant alterations in the cellular metabolic landscape attributable to SLC25A10 deficiency. In a targeted analysis (Fig. 16b), several dicarboxylates—including succinate, fumarate, and malonate—showed increased whole-cell abundance in SLC25A10-KD cells relative to controls. These measurements are consistent with altered dicarboxylate-related metabolism following SLC25A10 perturbation. However, because the measurements are not compartment-resolved, they do not identify mitochondrial efflux, exchange directionality, or the dominant SLC25A10 transport mode.

**Figure 16:**
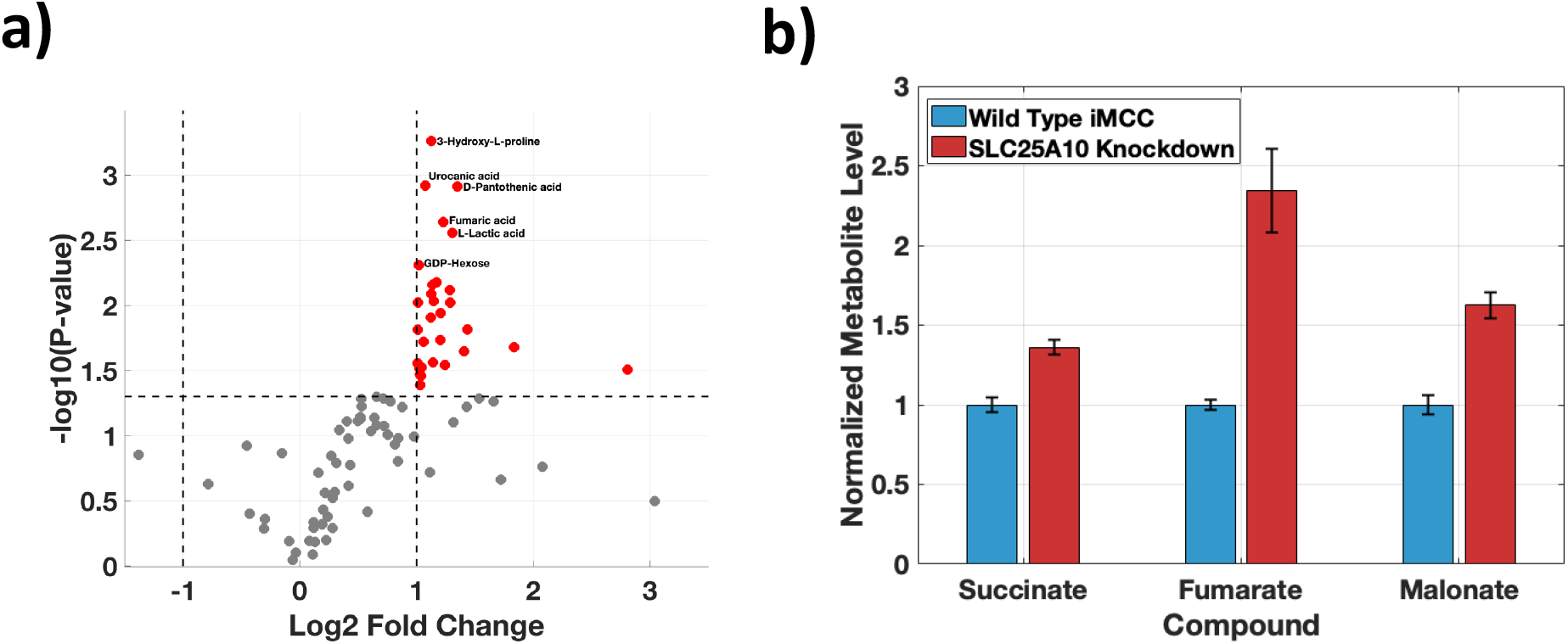
Targeted metabolomics shows increased dicarboxylate-related metabolite levels in SLC25A10 knockdown immortalized murine chromaffin cells (imCCs). (a) Volcano plot summarising differential metabolite analysis between SLC25A10 knockdown and wild-type imCCs. Red points indicate metabolites that increase significantly in the knockdown condition (e.g., succinate, fumarate, and malonate), while grey points denote non-significant changes (dashed lines indicate the fold-change and significance thresholds used in the analysis). (b) Normalised metabolite levels for succinate, fumarate, and malonate in wild-type (blue) and SLC25A10 knockdown (red) imCCs, showing higher levels in the knockdown condition.

Furthermore, a broader, untargeted view of the metabolome (Fig. 16a) underscored the extensive impact of SLC25A10 knockdown. This analysis highlighted a global metabolic reprogramming, with numerous metabolites exhibiting statistically significant changes in abundance. Among those notably elevated in SLC25A10-KD cells were L-lactic acid, 3-hydroxy-L-proline, urocanic acid, and D-pantothenic acid, in addition to fumaric acid. These widespread metabolic shifts suggest that SLC25A10 plays an integral role in maintaining broader cellular metabolic homeostasis, likely through its influence on the tricarboxylic acid (TCA) cycle and interconnected pathways.

We emphasise that these measurements reflect whole-cell metabolite pools and therefore conflate contributions from multiple subcellular compartments. Accordingly, we use the metabolomics results as qualitative context for the transporter modelling rather than as direct, compartment-resolved validation of transport directionality or thermodynamic driving forces.

### Appendix D.1 Method description

Cell Lines and Culture Conditions: Immortalized murine chromaffin cells (imCCs) [56] were used for all experiments. Cells were maintained in Dulbecco’s Modified Eagle Medium (DMEM) high glucose (Gibco, Cat# 11965), supplemented with 10% (v/v) fetal bovine serum (FBS; Sigma, Cat# 7524). Cultures were maintained at 37 °C in a humidified atmosphere containing 5% CO_2_. Cells were routinely passaged at 70–80% confluency using TrypLE and seeded for experiments.

Generation of Inducible Slc25a10 Knockdown imCCs: To generate imCCs with inducible knockdown of Slc25a10, a Tet-On 3G doxycycline-inducible shRNA lentiviral system (VectorBuilder) was used. This system involves sequential stable expression of the Tet regulatory protein and a Tet-On inducible knockdown sequence. Doxycycline (5 µg/mL) was added to the cells for 48 hours to induce knockdown, after which knockdown efficacy was confirmed via western blotting. Cells were then plated onto 6-well plates, and doxycycline was added the day after plating to induce knockdown. After 48 hours, metabolites were harvested from scramble and knockdown cells for downstream LC-MS analysis.

LC-MS Sample Preparation: Cells were washed with ice-cold saline and scraped into pre-cooled acetonitrile/methanol/water (40:40:20) containing 0.5% formic acid and 1 µg/mL D_6_-glutaric acid (internal standard), and immediately placed on ice. After 5 minutes, 8.8 µL of 15% NH_4_HCO_3_ per 100 µL of sample was added and mixed. Samples were immediately snap-frozen on dry ice for 15 minutes, then centrifuged at 14,800 g for 5 minutes at 4 °C. Supernatants were collected and stored at −80 °C until LC-MS analysis. LC-MS Analysis: Samples were analysed using an Agilent Infinity II HPLC coupled with an Agilent G6545A qTOF mass spectrometer. Metabolites were separated on a Waters Premier BEH Z-HILIC column (1.7 µm, 2.1 × 150 mm). Mobile phase A consisted of 20 mM ammonium bicarbonate with 0.1% ammonium hydroxide and 0.1% InfinityLab deactivator. Mobile phase B consisted of H_2_O:ACN (1:9 ratio) with 0.1% InfinityLab deactivator. The flow rate was 0.2 mL/min with the following gradient: 10% A held for 2 min, ramped to 35% A at 18 min, 70% A at 22 min, 90% A at 22.1 min (held for 2.9 min), followed by 10% A at 25.1 min and held for 5 min. Data processing was performed using Agilent Profinder 10. Normalisation to the internal standard and total protein content was performed using an in-house script.

## Appendix E Analytical Equilibrium Concentrations for SLC25A10

To derive analytical equilibrium expressions for the concentrations of mitochondrial succinate 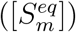, phosphate 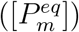, and malate 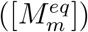, as well as external succinat 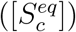, phosphate 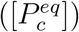, and malate 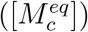, as functions of the initial mitochondrial concentrations 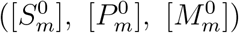 and external concentrations 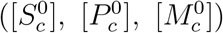, we employ conservation laws and exchange reaction free energy minimisation.

Here we derive closed-form expressions for the equilibrium concentrations of succinate and phosphate across the mitochondrial inner membrane mediated by the electroneutral antiporter SLC25A10. The derivation explicitly accounts for unequal compartment volumes and is based on mass conservation and thermodynamic equilibrium.

The analytical derivation relies on the following assumptions: the system consists of a well-mixed mitochondrial matrix and external side, each characterised by fixed volumes *V*_*m*_ and *V*_c_, respectively. Transport occurs via a one-to-one electroneutral antiport mechanism:

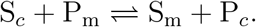

The system is considered at equilibrium, such that the free energy change of the transport reaction is zero. Only succinate and phosphate participate in the exchange reaction; malate concentration is assumed to be zero in this derivation. Volume changes due to osmotic effects are neglected. Total amounts of succinate and phosphate are conserved across compartments:

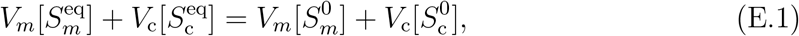

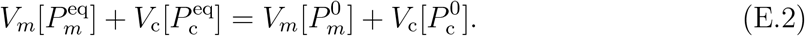

To enforce the strict 1:1 electroneutral antiport stoichiometry of SLC25A10, the transport process is parameterised by a single reaction extent *ξ*, which represents the net progress of the exchange cycle. Each transport event transfers exactly one succinate molecule in one direction and one phosphate molecule in the opposite direction, implying that changes in their concentrations are equal in magnitude and opposite in sign. Accordingly, the equilibrium concentrations in each compartment can be expressed as linear functions of the same exchange parameter, for example

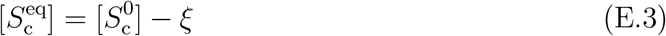

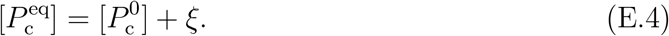

This formulation ensures mass conservation and electroneutrality by construction, while compartment volumes are incorporated separately when converting exchanged amounts to concentrations.

Substituting into Eqs. (E.1)–(E.2) yields:

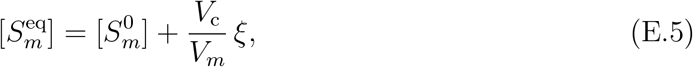

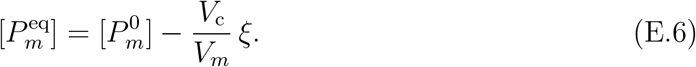

For an electroneutral exchange reaction, the free energy change is:

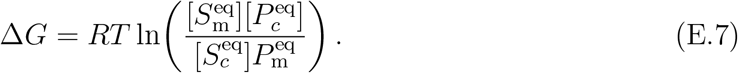

At equilibrium, Δ*G* = 0, which implies:

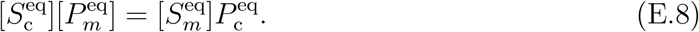

Substituting the expressions above into Eq. (E.8) and simplifying yields a linear equation for *ξ*, whose solution is:

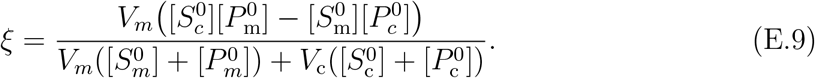

Substituting Eq. (E.9) into the expressions for the equilibrium concentrations yields:

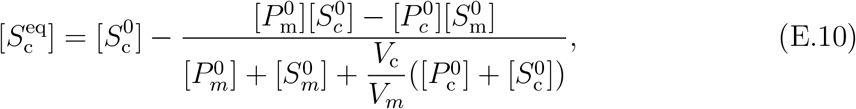

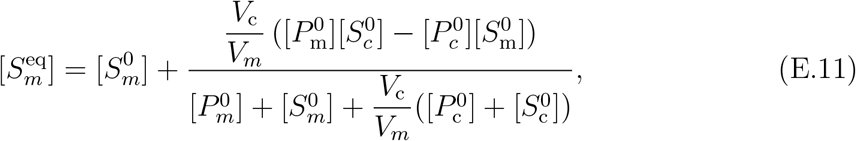

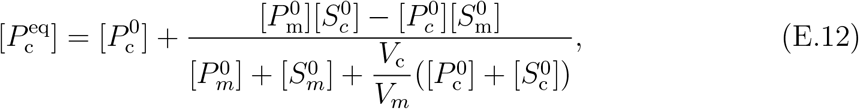

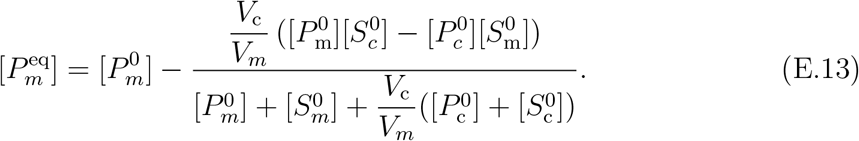

These expressions reduce to the classical concentration-only formulas in the special case *V*_*m*_ = *V*_c_. An analogous derivation can be carried out for malate–phosphate and malate–succinate exchange by assuming zero succinate and phosphate concentrations, respectively.

## Appendix F Proteoliposome Compartment Volumes

This appendix describes how the internal and external compartment volumes were parameterised for the proteoliposome assays. All primary quantities were taken directly from Indiveri et al. [23] (Table 10). The only non-tabulated value extracted from a figure is the intraliposomal (internal) volume at the optimised condition (13 Amberlite passages), which was read from Fig. 2 of Indiveri et al. [23] and treated as an approximate value.

**Table 9:** Model parameters, definitions, and units.

| Symbol | Description | Units |
| --- | --- | --- |
| $\tilde{V}_m$ | Matrix volume per gram protein ($V_m/g$) | $\text{L g}^{-1}$ |
| $\tilde{V}_c$ | External volume per gram protein ($V_c/g$) | $\text{L g}^{-1}$ |
| $V_m$ | Matrix compartment volume | $\text{L}$ |
| $V_c$ | External compartment volume | $\text{L}$ |
| $k_1^s$ | Succinate binding rate of matrix side | $\text{L mol}^{-1} \text{s}^{-1}$ |
| $k_{-1}^s$ | Succinate unbinding rate of matrix side | $\text{s}^{-1}$ |
| $k_3^s$ | Succinate binding rate of external side | $\text{L mol}^{-1} \text{s}^{-1}$ |
| $k_{-3}^s$ | Succinate unbinding rate of external side | $\text{s}^{-1}$ |
| $k_c^s$ | Conformational rate of succinate-loaded carrier | $\text{s}^{-1}$ |
| $k_{-c}^s$ | Reverse conformational rate of succinate-loaded carrier | $\text{s}^{-1}$ |
| $k_1^m$ | Malate binding rate of matrix side | $\text{L mol}^{-1} \text{s}^{-1}$ |
| $k_{-1}^m$ | Malate unbinding rate of matrix side | $\text{s}^{-1}$ |
| $k_3^m$ | Malate binding rate of external side | $\text{L mol}^{-1} \text{s}^{-1}$ |
| $k_{-3}^m$ | Malate unbinding rate of external side | $\text{s}^{-1}$ |
| $k_c^m$ | Conformational rate of malate-loaded carrier | $\text{s}^{-1}$ |
| $k_{-c}^m$ | Reverse conformational rate of malate-loaded carrier | $\text{s}^{-1}$ |
| $k_1^p$ | Phosphate binding rate of external side, | $\text{L mol}^{-1} \text{s}^{-1}$ |
| $k_{-1}^p$ | Phosphate unbinding rate of external side | $\text{s}^{-1}$ |
| $k_3^p$ | Phosphate binding rate of matrix side | $\text{L mol}^{-1} \text{s}^{-1}$ |
| $k_{-3}^p$ | Phosphate unbinding rate of matrix side | $\text{s}^{-1}$ |
| $k_c^p$ | Conformational rate of phosphate-loaded carrier | $\text{s}^{-1}$ |
| $k_{-c}^p$ | Reverse conformational rate of phosphate-loaded carrier | $\text{s}^{-1}$ |
| $K_D^x$ | Matrix side dissociation constant for $x \in \{s, m, p\}$ | $\text{mol L}^{-1}$ |
| $K_D^x$ | External side dissociation constant for $x \in \{s, m, p\}$ | $\text{mol L}^{-1}$ |
| $T_{\max}^x$ | Maximal Transport capacity for $x \in \{s, m, p\}$ | $\text{mol g}^{-1} \text{s}^{-1}$ |
| $\lambda_{21}$ | Conformational-bias ratio (malate vs. succinate weighting) | dimensionless |
| $\lambda_{31}$ | Conformational-bias ratio (phosphate vs. succinate weighting) | dimensionless |
| $\lambda_{23}$ | Conformational-bias ratio (phosphate vs. malate weighting) | dimensionless |
| $\phi$ | Weighted ratio of matrix to external substrate levels | dimensionless |
| $\delta_1$ | Matrix side saturation factor | dimensionless |
| $\delta_2$ | External side saturation factor | dimensionless |

*Note:* $L$ denotes liters, $s$ denotes seconds, $g$ denotes grams of protein, and $\text{mol}$ denotes moles.

**Table 10:** Primary quantities taken from Indiveri et al. [23] and used as inputs to the volume calculations.

| Symbol | Quantity (as reported) | Value used here |
| --- | --- | --- |
| $V_{\text{mix}}$ | Total reconstitution mixture volume | 0.68 mL = 680 $\mu\text{L}$ |
| $V_{\text{assay}}$ | Aliquot (reaction) volume used per transport assay | 100 $\mu\text{L}$ |
| $m_{\text{PL}}$ | Total phospholipid mass in the reconstitution mixture | 6.7 mg |
| $m_{\text{prot}}$ | Total protein mass in the reconstitution mixture | 7–15 $\mu\text{g}$ |
| $V_{\text{in}}^{\text{approx}}$ | Intraliposomal volume at 13 Amberlite passages | $\approx 4.0 \mu\text{L}/\text{mg PL}$ |

The external (extraliposomal) compartment volume in the transport assay is defined as the total assay volume *V*_assay_ minus the total internal volume *V*_*m*_. Therefore, the total external volume *V*_*c*_ is

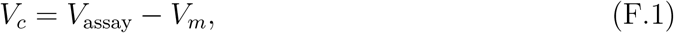

where *V*_assay_ = 100 *µ*L. To express the external volume in the same normalised units as the internal compartment (volume per mass of reconstituted carrier protein), we first compute the protein mass per assay from the mixture composition. The protein mass was reported as a range (7–15 *µ*g) in the full reconstitution mixture volume (*V*_mix_ = 680 *µ*L). We therefore define the average protein mass in the mixture as

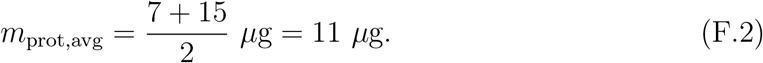

Because each assay used a 100 *µ*L aliquot of the 680 *µ*L mixture, the protein mass per assay is

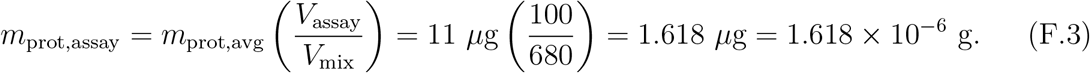

The total internal (intraliposomal) aqueous volume present in the full reconstitution mixture is obtained by multiplying the intraliposomal volume per lipid mass (read from Fig. 2 of Indiveri et al. [23] at the optimised condition) by the total lipid mass used in the mixture:

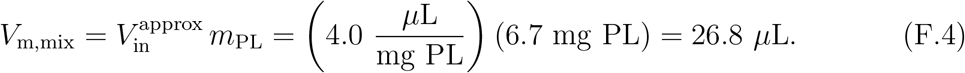

The specific internal volume, expressed as internal volume per gram of reconstituted protein, is then

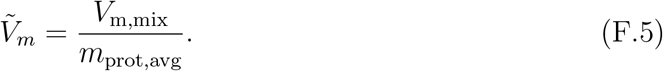

Using *V*_m,mix_ = 26.8 *µ*L and *m*_prot,avg_ = 11 *µ*g yields

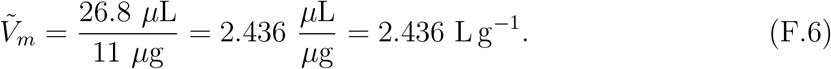

The total internal volume corresponding to a single 100 *µ*L assay aliquot is

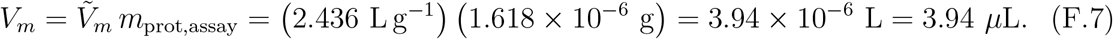

From (F.1), the total external volume is

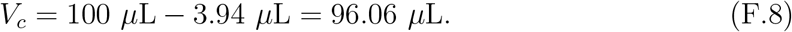

Finally, the specific external volume is defined by normalising the external volume by the protein mass present in the same assay aliquot:

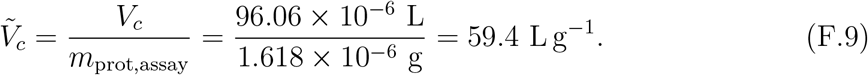

Together, these definitions provide consistent internal and external compartment volumes in both absolute units (*µ*L per assay) and normalised units (L g^−1^ protein).

## Appendix G Determination of Compartment Volumes in Intact Rat Liver Mitochondria Assays

The uptake assays using intact rat liver mitochondria in Palmieri et al. [3] follow an inhibitor-stop and rapid-centrifugation protocol. In these experiments, intramitochondrial concentrations are obtained by converting the measured intrapellet radioactivity into an amount (e.g., nmol) and dividing by the aqueous volume that is inaccessible to extracellular sucrose (the sucrose-inaccessible space). In parallel measurements, ^3^H_2_O and [^14^C]sucrose are used to estimate total pellet water and the sucrose-permeable space, respectively, which together determine the intramitochondrial aqueous space relevant for solute accumulation. Because Palmieri et al. [3] do not report a single numerical value for the sucrose-inaccessible space across all experiments, we adopt a literature value for rat liver mitochondria under comparable conditions for the specific internal volume [38].

The specific internal volume is defined as the sucrose-inaccessible aqueous space per unit mitochondrial protein. In our model notation, this corresponds to the matrix volume per protein, 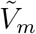. The total internal volume is the internal aqueous volume present in a single assay and equals the specific internal volume multiplied by the protein mass used in that assay; this corresponds to the matrix volume *V*_*m*_ in our model. The total external volume is the incubation-medium volume in which mitochondria are suspended; this corresponds to the external compartment volume *V*_*c*_ in our model. The specific external volume is the external volume normalised per unit mitochondrial protein; this corresponds to 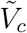 in our model.

Table 11 summarises the quantities reported in Palmieri et al. [3] and the literature value used for the sucrose-inaccessible aqueous space [38]. The incubation (external) reaction volume was not explicitly specified; therefore, we assume an incubation volume of 1.0 mL. In our model notation, this incubation medium corresponds to the external compartment volume *V*_*c*_.

**Table 11:** Primary quantities taken from Palmieri et al. [3] and Harris et al. [38] and used as inputs to the compartment-volume calculations for intact rat liver mitochondria uptake assays.

| Symbol | Quantity (as reported) | Value used here |
| --- | --- | --- |
| $m_p$ | Mitochondrial protein per assay (range) | 1.55–2.50 mg |
| $\tilde{V}_m$ | Sucrose-inaccessible water space (mean) | 0.70 mL g$^{-1}$ |
| $V_c$ | External incubation volume | Not stated (assume 1.0 mL) |

A literature mean value for the sucrose-inaccessible water space of rat liver mitochondria was used:

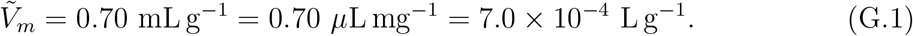

For an assay containing mitochondrial protein mass *m*_*p*_ (in mg), the total internal aqueous volume is

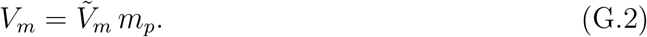

Since the *m*_*p*_ range is given, we used mean protein mass 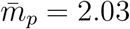 mg and (G.1),

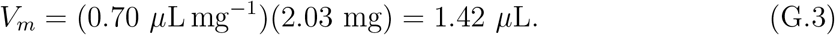

The specific external volume is the incubation volume normalised by mitochondrial protein mass:

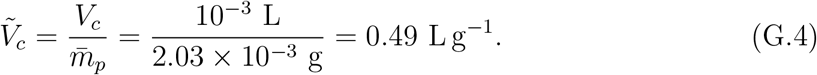

## Appendix H Finite-Difference versus Complex-Step Sensitivity Comparison

To assess numerical accuracy of the finite-difference sensitivity coefficients reported in Section 3.7, we compute the same local sensitivities using the complex-step method [42]. Figure 17 reports absolute and relative differences between the two estimates, showing small discrepancies across all parameters (maximum error *<* 8 × 10^−4^).

**Figure 17:**
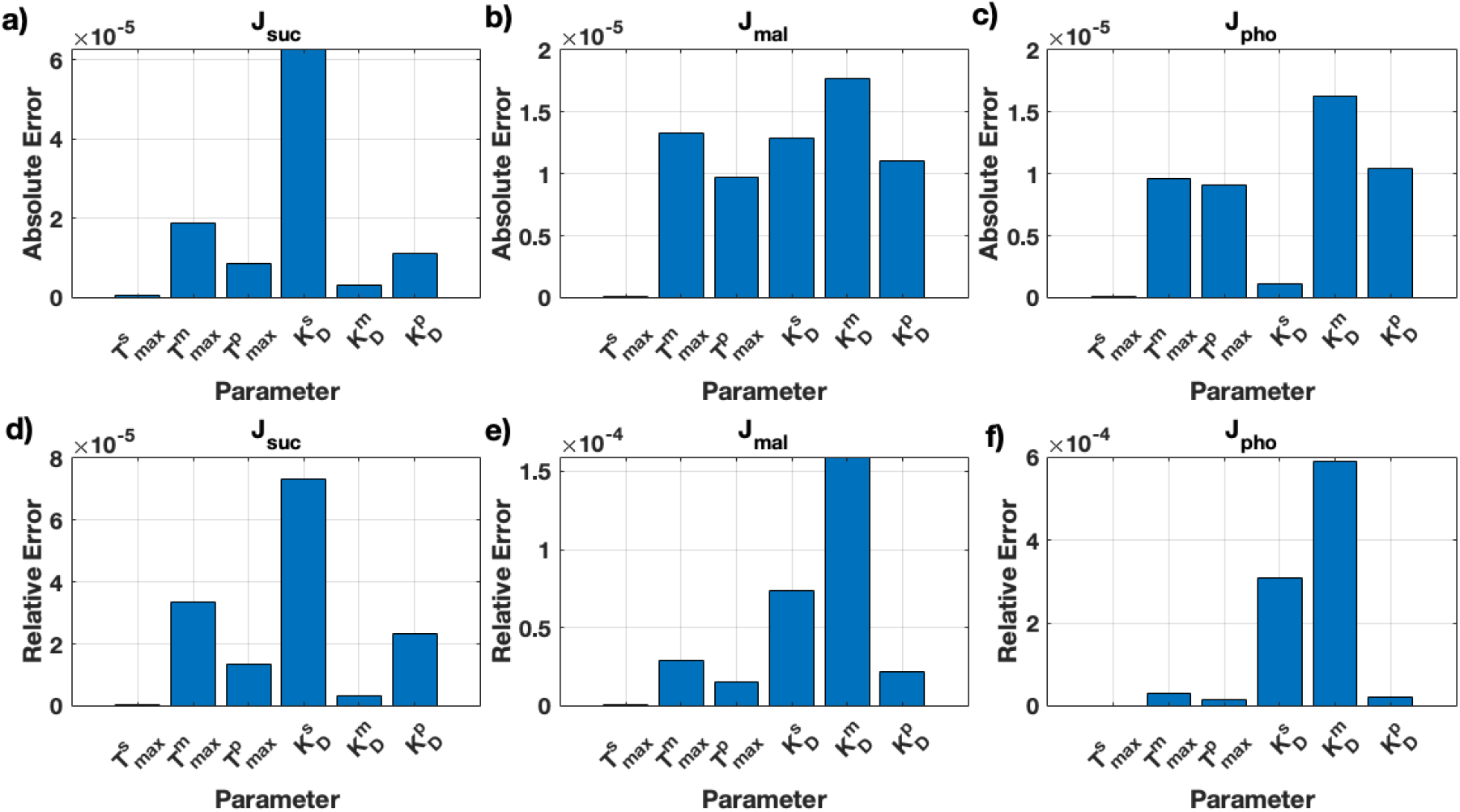
Comparison of finite-difference and complex-step sensitivity estimates. Absolute (a–c) and relative (d–f) differences between sensitivity coefficients computed by finite differences (using the perturbation described in Section 3.7) and the complex-step method.

## References

[1] Camila Cimadamore-Werthein et al. “Human mitochondrial carriers of the SLC25 family function as monomers exchanging substrates with a ping-pong kinetic mechanism”. In: EMBO Journal (2024). ISSN: 14602075. doi: 10.1038/s44318-024-00150–0.

[2] Edmund R.S. Kunji et al. “The SLC25 carrier family: Important transport proteins in mitochondrial physiology and pathology”. In: Physiology 35 (5 Sept. 2020), pp. 302–327. ISSN: 15489221. doi: 10.1152/physiol.00009.2020.

[3] Ferdinando Palmieri et al. “Kinetic Study of the Dicarboxylate Carrier in Rat Liver Mitochondria”. In: European Journal of Biochemistry 22 (1 1971), pp. 66–74. ISSN: 14321033. doi: 10.1111/j.1432-1033.1971.tb01515.x.

[4] Luc Rochette et al. Mitochondrial SLC25 carriers: Novel targets for cancer therapy. May 2020. doi: 10.3390/molecules25102417.

[5] Adrian Casas-Benito, Sonia Martínez-Herrero, and Alfredo Martínez. Succinate-Directed Approaches for Warburg Effect-Targeted Cancer Management, an Alternative to Current Treatments? May 2023. doi: 10.3390/cancers15102862.

[6] Amer Ahmed et al. The Role of Mitochondrial Solute Carriers SLC25 in Cancer Metabolic Reprogramming: Current Insights and Future Perspectives. Jan. 2025. doi: 10.3390/ijms26010092.

[7] Eva Pyrihová et al. “A mitochondrial carrier transports glycolytic intermediates to link cytosolic and mitochondrial glycolysis in the human gut parasite Blastocystis”. In: eLife 13 (May 2024). ISSN: 2050084X. doi: 10.7554/ELIFE.94187.

[8] Shinji Mizuarai et al. “Identification of dicarboxylate carrier Slc25a10 as malate transporter in de Novo fatty acid synthesis”. In: Journal of Biological Chemistry 280 (37 Sept. 2005), pp. 32434–32441. ISSN: 00219258. doi: 10.1074/jbc.M503152200.

[9] Charlotte Lussey-Lepoutre et al. Mitochondrial deficiencies in the predisposition to paraganglioma. June 2017. doi: 10.3390/metabo7020017.

[10] Julian Hlouschek et al. “Targeting SLC25A10 alleviates improved antioxidant capacity and associated radioresistance of cancer cells induced by chronic-cycling hypoxia”. In: Cancer Letters 439 (Dec. 2018), pp. 24–38. ISSN: 18727980. doi: 10.1016/j.canlet.2018.09.002.

[11] Ferdinando Palmieri, Pasquale Scarcia, and Magnus Monné. Diseases caused by mutations in mitochondrial carrier genes SLC25: A review. Apr. 2020. doi: 10.3390/biom10040655.

[12] Katarina Kluckova and Daniel A Tennant. “Metabolic implications of hypoxia and pseudohypoxia in pheochromocytoma and paraganglioma”. In: Cell and tissue research 372 (2018), pp. 367–378.

[13] Katarína Klučková et al. “Succinate dehydrogenase deficiency in a chromaffin cell model retains metabolic fitness through the maintenance of mitochondrial nadh oxidoreductase function”. In: FASEB Journal 34 (1 2020), pp. 303–315. ISSN: 15306860. doi: 10.1096/fj.201901456R.

[14] Elías Vera-Sigüenza et al. “A Mathematical Exploration of SDH-b Loss in Chromaffin Cells”. In: Bulletin of Mathematical Biology 87 (2024). url: https://api.semanticscholar.org/CorpusID:271269243.

[15] Jason N. Bazil, Gregery T. Buzzard, and Ann E. Rundell. “Modeling mitochondrial bioenergetics with integrated volume dynamics”. In: PLoS Computational Biology 6 (1 2010). ISSN: 15537358. doi: 10.1371/journal.pcbi.1000632.

[16] Xiao Zhang et al. “Integrated computational model of the bioenergetics of isolated lung mitochondria”. In: PLoS ONE 13 (6 June 2018). ISSN: 19326203. doi: 10.1371/journal.pone.0197921.

[17] Shima Sadri et al. “Computational Modeling of Substrate-Dependent Mitochondrial Respiration and Bioenergetics in the Heart and Kidney Cortex and Outer Medulla”. In: Function 4 (5 2023). ISSN: 26338823. doi: 10.1093/function/zqad038.

[18] Jonathan J. Ruprecht and Edmund R. S. Kunji. “Structural Mechanism of Transport of Mitochondrial Carriers.” In: Annual review of biochemistry (2021). URL: https://api.semanticscholar.org/CorpusID:231874184.

[19] Vasiliki Mavridou et al. “Substrate binding in the mitochondrial ADP/ATP carrier is a step-wise process guiding the structural changes in the transport cycle”. In: Nature Communications 13 (1 Dec. 2022). ISSN: 20411723. doi: 10.1038/s41467-022-31366-5.

[20] F. Palmieri. “Mitochondrial carrier proteins”. In: FEBS Letters 346 (1 June 1994), pp. 48–54. ISSN: 00145793. doi: 10.1016/0014-5793(94)00329-7.

[21] R. N. Johnson and J. Brian Chappell. “The transport of inorganic phosphate by the mitochondrial dicarboxylate carrier.” In: The Biochemical journal 134 3 (1973), pp. 769–74. URL: https://api.semanticscholar.org/CorpusID:22142227.

[22] Faustino Bisaccia, Cesare Indiveri, and Ferdinando Palmieri. “Purification and reconstitution of two anion carriers from rat liver mitochondria: the dicarboxylate and the 2-oxoglutarate carrier.” In: Biochimica et biophysica acta 933 2 (1988), pp. 229–40. url: https://api.semanticscholar.org/CorpusID:25273025.

[23] Cesare Indiveri et al. “Kinetics of the reconstituted dicarboxylate carrier from rat liver mitochondria.” In: Biochimica et biophysica acta 977 2 (1989), pp. 187–93. URL: https://api.semanticscholar.org/CorpusID:1667771.

[24] Cesare Indiveri et al. “Kinetic discrimination of two substrate binding sites of the reconstituted dicarboxylate carrier from rat liver mitochondria.” In: Biochimica et biophysica acta 977 2 (1989), pp. 194–9. URL: https://api.semanticscholar.org/CorpusID:40809880.

[25] Emmet A Francis et al. “Spatial modeling algorithms for reactions and transport in biological cells”. en. In: Nat. Comput. Sci. 5.1 (Jan. 2025), pp. 76–89.

[26] Guadalupe C Garcia et al. “Mitochondrial morphology provides a mechanism for energy buffering at synapses”. en. In: Sci. Rep. 9.1 (Dec. 2019), p. 18306.

[27] Sabzali Javadov, Xavier Chapa-Dubocq, and Vladimir Makarov. “Different approaches to modeling analysis of mitochondrial swelling”. In: Mitochondrion 38 (Jan. 2018), pp. 58–70.

[28] Raquel Adams et al. “How the topology of the mitochondrial inner membrane modulates ATP production”. en. In: Cells 14.4 (Feb. 2025), p. 257.

[29] Nasrin Afzal et al. “Effect of crista morphology on mitochondrial ATP output: A computational study”. en. In: Curr. Res. Physiol. 4 (Apr. 2021), pp. 163–176.

[30] Edward L. King and Carl Altman. “A Schematic Method of Deriving the Rate Laws for Enzyme-Catalyzed Reactions”. In: The Journal of Physical Chemistry 60 (1956), pp. 1375–1378. URL: https://api.semanticscholar.org/CorpusID:101759412.

[31] Alex Berlaga and Anatoly B. Kolomeisky. “Molecular Mechanisms of Active Transport in Antiporters: Kinetic Constraints and Efficiency”. In: Journal of Physical Chemistry Letters 12 (39 Oct. 2021), pp. 9588–9594. ISSN: 19487185. doi: 10.1021/acs.jpclett.1c02846.

[32] Alex Berlaga and Anatoly B. Kolomeisky. “Understanding Mechanisms of Secondary Active Transport by Analyzing the Effects of Mutations and Stoichiometry”. In: Journal of Physical Chemistry Letters 13 (24 June 2022), pp. 5405–5412. ISSN: 19487185. doi: 10.1021/acs.jpclett.2c01232.

[33] Alex Berlaga and Anatoly B. Kolomeisky. “Theoretical study of active secondary transport: Unexpected differences in molecular mechanisms for antiporters and symporters”. In: Journal of Chemical Physics 156 (8 Feb. 2022). ISSN: 10897690. doi: 10.1063/5.0082589.

[34] Lucy R. Forrest, Reinhard Krämer, and Christine Ziegler. “The structural basis of secondary active transport mechanisms.” In: Biochimica et biophysica acta 1807 2 (2011), pp. 167–88. URL: https://api.semanticscholar.org/CorpusID:10803966.

[35] Jonathan J. Ruprecht and Edmund R.S. Kunji. The SLC25 Mitochondrial Carrier Family: Structure and Mechanism. Mar. 2020. doi: 10.1016/j.tibs.2019.11.001.

[36] Roger Springett et al. “Modelling the free energy profile of the mitochondrial ADP/ATP carrier”. In: Biochimica et Biophysica Acta - Bioenergetics 1858 (11 Nov. 2017), pp. 906–914. ISSN: 18792650. doi: 10.1016/j.bbabio.2017.05.006.

[37] James Keener and James Sneyd. Mathematical Physiology II: Systems Physiology. Springer, 2009.

[38] Eric J. Harris and Karel van Dam. “Changes of total water and sucrose space accompanying induced ion uptake or phosphate swelling of rat liver mitochondria.” In: The Biochemical journal 106 3 (1968), pp. 759–66. URL: https://api.semanticscholar.org/CorpusID:37995416.

[39] Cesare Indiveri et al. “Kinetic characterization of the reconstituted dicarboxylate carrier from mitochondria: a four-binding-site sequential transport system.” In: Biochimica et biophysica acta 1143 3 (1993), pp. 310–8. URL: https://api.semanticscholar.org/CorpusID:8502488.

[40] Alejandro F. Villaverde et al. “A protocol for dynamic model calibration”. In: Briefings in Bioinformatics 23 (1 Jan. 2022). ISSN: 14774054. doi: 10.1093/bib/bbab387.

[41] Alejandro F. Villaverde, Antonio Barreiro, and Antonis Papachristodoulou. “Structural Identifiability of Dynamic Systems Biology Models”. In: PLoS Computational Biology 12 (10 Oct. 2016). ISSN: 15537358. doi: 10.1371/journal.pcbi.1005153.

[42] William Squire and George Trapp. “Using complex variables to estimate derivatives of real functions”. In: SIAM Rev. Soc. Ind. Appl. Math. 40.1 (Jan. 1998), pp. 110–112.

[43] Hanwen Huang, Andreas Handel, and Xiao Song. “A Bayesian approach to estimate parameters of ordinary differential equation”. In: Computational Statistics 35 (3 Sept. 2020), pp. 1481–1499. ISSN: 16139658. doi: 10.1007/s00180-020-00962-8.

[44] Christophe Andrieu et al. “On the utility of Metropolis-Hastings with asymmetric acceptance ratio”. In: (Mar. 2018). URL: http://arxiv.org/abs/1803.09527.

[45] Aki Vehtarh et al. “Rank-Normalization, Folding, and Localization: An Improved (Formula presented) for Assessing Convergence of MCMC (with Discussion)*†”. In: Bayesian Analysis 16 (2 2021), pp. 667–718. ISSN: 19316690. doi: 10.1214/20-BA1221.

[46] Andrew Gelman et al. Bayesian Data Analysis. 3rd. London: Chapman & Hall/CRC, 2013.

[47] Bob Carpenter et al. “Stan: A probabilistic programming language”. In: Journal of Statistical Software 76 (1 2017). ISSN: 15487660. doi: 10.18637/jss.v076.i01.

[48] Ying Liu, Dootika Vats, and James M. Flegal. “Batch Size Selection for Variance Estimators in MCMC”. In: Methodology and Computing in Applied Probability 24 (1 Mar. 2022), pp. 65–93. ISSN: 15737713. doi: 10.1007/s11009-020-09841-7.

[49] Andrew Gelman and Donald B. Rubin. “Inference from Iterative Simulation Using Multiple Sequences”. In: Statistical Science 7 (1992), pp. 457–472. URL: https://api.semanticscholar.org/CorpusID:14661921.

[50] P. Stephen et al. “General methods for monitoring convergence of iterative simulations”. In: Journal of Computational and Graphical Statistics 7 (1998), pp. 434–455. URL: https://api.semanticscholar.org/CorpusID:7300890.

[51] Stefan J. Jol et al. “Thermodynamic calculations for biochemical transport and reaction processes in metabolic networks”. In: Biophysical Journal 99 (10 Nov. 2010), pp. 3139–3144. ISSN: 15420086. doi: 10.1016/j.bpj.2010.09.043.

[52] N. Tepper et al. “Steady-State Metabolite Concentrations Reflect a Balance between Maximizing Enzyme Efficiency and Minimizing Total Metabolite Load”. In: PLoS ONE 8 (2013). URL: https://api.semanticscholar.org/CorpusID:1931484.

[53] X. Li et al. “A database of thermodynamic quantities for the reactions of glycolysis and the tricarboxylic acid cycle”. In: Journal of Physical Chemistry B 114 (49 Dec. 2010), pp. 16068–16082. ISSN: 15205207. doi: 10.1021/jp911381p.

[54] G. M. Tannahill et al. “Succinate is an inflammatory signal that induces IL-1β through HIF-1α”. In: Nature 496 (7444 Apr. 2013), pp. 238–242. ISSN: 00280836. doi: 10.1038/nature11986.

[55] W. Wallace Cleland. “Derivation of rate equations for multisite ping-pong mechanisms with ping-pong reactions at one or more sites.” In: The Journal of biological chemistry 248 24 (1973), pp. 8353–5. URL: https://api.semanticscholar.org/CorpusID:26861029.

[56] Eric Letouzé et al. “SDH mutations establish a hypermethylator phenotype in paraganglioma”. In: Cancer cell 23.6 (2013), pp. 739–752.

